# A conserved tetrapolar mating system permits sexual hybridization and recombinant host phenotypes in wheat- and barley-adapted *Puccinia striiformis*

**DOI:** 10.64898/2026.05.01.721896

**Authors:** Shideh Mojerlou, Zhenyan Luo, Rita Tam, Mareike Möller, Ashley Jones, Benjamin Schwessinger, Julian Rodriguez-Algaba

## Abstract

Host specialization can restrict gene flow in fungal pathogens, yet sexual compatibility may persist when mating occurs on a shared alternate host. We tested this possibility by crossing wheat-adapted *Puccinia striiformis* f. sp. *tritici* and barley-adapted *P. striiformis* f. sp. *hordei* on *Berberis vulgaris*. Despite contrasting host specialization and genome-wide sequence differentiation, reciprocal crosses produced 18 viable aeciospore-derived progeny. Chromosome-scale parental assemblies revealed physically unlinked homeodomain (HD) and pheromone receptor (PR) mating-type loci, consistent with a conserved tetrapolar mating system. All progeny inherited one HD haplotype from each parent, representing all four parental HD haplotype combinations. Phenotyped progeny displayed host-interaction profiles distinct from either parent, including compatibility with selected genotypes of both wheat and barley. These findings demonstrate that host-associated differentiation in *P. striiformis* can coexist with incomplete intrinsic reproductive isolation between the tested parental isolates, allowing sexual hybridization to generate novel host phenotypes.

## Introduction

Host specialization is widespread among fungal plant pathogens, where closely related lineages become associated with particular host species. Such specialization can involve adaptive interactions between pathogen effectors and host immune receptors, as well as alleles that influence pathogen performance in different host environments (Stukenbrock 2013; Thines 2019). Host-associated adaptation can reduce effective gene flow through ecological barriers, including poor infection success on non-adapted primary hosts and reduced fitness of recombinant genotypes. However, ecological barriers to gene flow do not necessarily imply complete intrinsic reproductive isolation, particularly in pathogens whose sexual stage occurs outside the primary host environment. In such systems, recombination on shared sexual hosts can generate new combinations of host-interaction loci despite long periods of clonal growth on primary hosts (Stukenbrock 2016; Zhao et al. 2016).

Population genomic and experimental studies show that hybridization and recombination can generate novel pathogen genotypes and alter host compatibility, as illustrated by *Zymoseptoria pseudotritici* and triticale powdery mildew (*Blumeria graminis* f. sp. *triticale*) (Stukenbrock et al. 2012; Menardo et al. 2016; Müller et al. 2021). Studies of closely related host-specialized fungal pathogens, including *Venturia inaequalis*, *Ascochyta* spp. and citrus-associated *Alternaria* pathotypes, further show that host specialization can structure pathogen populations and contribute to ecological reproductive barriers (Peever et al. 1999; Hernandez-Bello et al. 2006; Gladieux et al. 2010). In cereal rust fungi (*Pucciniales*), historical crossing studies similarly showed that sexual crosses between differently adapted pathogen forms on a common sexual host can yield progeny with altered host compatibility (reviewed by Anikster 1984). However, the genomic basis and evolutionary persistence of this reproductive compatibility have remained largely untested.

Many cereal-infecting rust fungi undergo asexual infection on their primary hosts but reproduce sexually on phylogenetically distant alternate hosts, allowing mating to occur outside the selective environment imposed by the primary cereal host (Zhao et al. 2016; Schwessinger 2017; Aime et al. 2018; Rodriguez-Algaba et al. 2022). During the sexual cycle, nuclear fusion (karyogamy) in teliospores is followed by meiosis, producing haploid basidiospores that infect the alternate host and develop pycnia. Fusion of compatible pycniospores and receptive hyphae (plasmogamy) then establishes the dikaryotic stage, which produces aeciospores capable of infecting cereal hosts (Mojerlou et al. 2025). In grass-infecting rusts and powdery mildews, host specialization is often described using *formae speciales*, i.e., morphologically similar pathogen forms specialized to different host species (Eriksson 1894). Yet some isolates assigned to a single *forma specialis* can infect multiple cereal species, indicating that host compatibility does not always map strictly onto *forma specialis* boundaries (Bettgenhaeuser et al. 2014; Dracatos et al. 2018). This separation between primary-host specialization and sexual reproduction makes rust fungi suitable for testing whether host specialization and intrinsic reproductive compatibility can remain decoupled.

The wheat-adapted *P. striiformis* f. sp. *tritici* (*Pst*) and barley-adapted *P. striiformis* f. sp. *hordei* (*Psh*) are closely related but host-specialized *formae speciales* within the *P. striiformis* complex (Wellings 2011; Xia et al. 2018b), and both utilize *Berberis* spp. as alternate hosts for sexual reproduction (Jin et al. 2010; Zhao et al. 2016; Huang et al. 2019; Rodriguez-Algaba et al. 2021). Population genetic surveys in regions where wheat and barley are co-cultivated have reported patterns suggestive of occasional admixture/recombination between *Pst* and *Psh*, but direct experimental evidence for sexual compatibility between these host-adapted forms remains limited (Du et al. 2024).

In many basidiomycetes, mating compatibility is governed by a tetrapolar mating-type (MAT) system in which two genetically unlinked loci control compatibility: a homeodomain (HD) locus encoding transcription factors (*bW-HD1* and *bE-HD2*) that regulate post-mating development, and a pheromone-receptor (PR) locus encoding pheromone receptors (*Pra*) and pheromone peptide precursors (*mfa*) that mediate pre-mating recognition (Coelho et al. 2017; Cuomo et al. 2017; Luo et al. 2024). In a tetrapolar system, compatibility requires appropriate mating-type specificities at both independently segregating MAT loci, thereby restricting compatibility among related meiotic products while permitting outcrossing between isolates carrying compatible MAT specificities.

Multiallelic HD loci can be maintained by negative frequency-dependent selection, which favors rare mating-type alleles and preserves allelic diversity over extended evolutionary timescales (Nieuwenhuis et al. 2013; Coelho et al. 2017; Luo et al. 2024). Comparative genomics indicates that cereal rust fungi, including *Pst*, *P. graminis* f. sp. *tritici*, *P. triticina* and *P. coronata* f. sp. *avenae*, retain this MAT organization, with HD and PR loci located on separate chromosomes (Henningsen et al. 2024; Luo et al. 2024; Henningsen et al. 2026). However, whether HD alleles from wheat- and barley-adapted *P. striiformis* forms partition by *forma specialis*, and whether these forms retain sexual compatibility within a conserved tetrapolar mating system, remain unresolved.

Here, we used controlled reciprocal crosses between *Pst* and *Psh* on *B. vulgaris* to test whether host-associated differentiation has been accompanied by intrinsic reproductive isolation and corresponding differentiation in mating-type variation. We combined chromosome-scale parental assemblies and genome-wide sequence comparisons with MAT-locus characterization, progeny genotyping, and host phenotyping. We show that the tested wheat- and barley-adapted parental isolates retain sexual compatibility within a conserved tetrapolar MAT system despite genome-wide sequence differentiation and contrasting host-specialized infection phenotypes. The resulting progeny displayed recombinant host-compatibility profiles, including compatibility with subsets of both wheat and barley genotypes. Together, these results demonstrate incomplete intrinsic reproductive isolation and show that host-associated differentiation in *P. striiformis* can coexist with retained sexual compatibility and the generation of recombinant host phenotypes in the progeny.

## Results and Discussion

### Chromosome-scale, haplotype-resolved assemblies of the parental isolates

DK02d_12 (*Pst*) and NP85002 (*Psh*) were selected as a phenotypically contrasting parental pair. DK02d_12 showed compatibility on wheat but not barley, whereas NP85002 showed compatibility on barley but not wheat across the tested host panel, as described below (“Progeny display recombinant host-compatibility phenotypes, including dual-host compatibility”). Chromosome-scale, haplotype-resolved assemblies are available for both parents (NP85002, Tam et al. 2026; DK02d_12 generated here), enabling genome-wide comparisons and chromosome-level characterization of mating-type loci. Both assemblies showed high consensus accuracy and near-complete telomere recovery, with strong Hi-C support for haplotype separation (DK02d_12: QV 65.06, 68/72 chromosome ends, 99.00% within-haplotype Hi-C links; NP85002: QV 72.01, 69/72 chromosome ends, 97.54% within-haplotype links). BUSCO completeness was 86.10% and 90.10% for DK02d_12 and NP85002, respectively (Fig. 1a; Table S2; Tam et al. 2026).

**Figure 1.**
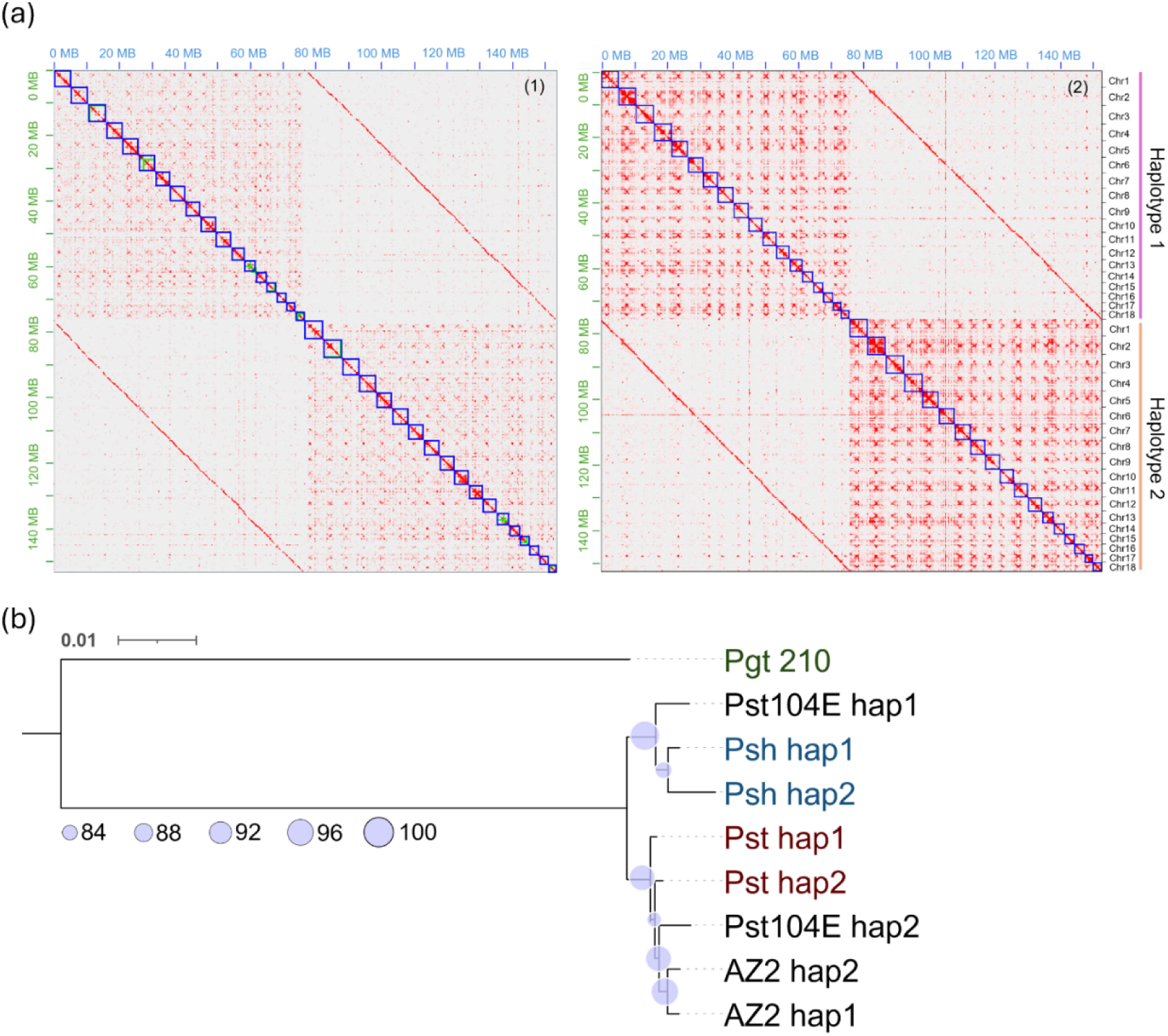
Chromosome-scale genome assemblies and genealogical relationships of parental isolates. **(a)** Hi-C contact maps of the haplotype-resolved parental assemblies visualized in Juicebox. Panel (1) shows *Pst* DK02d_12 and panel (2) shows *Psh* NP85002 (assembly from Tam et al. 2026). Each parental isolate is dikaryotic, containing two haploid nuclear genomes that were assembled separately. Each contact map displays one phased haplotype comprising 18 chromosomes, with strong diagonal interaction patterns and limited offdiagonal structure supporting chromosome-scale scaffolding. **(b)** Genome-wide genealogy inferred from single-copy orthologues using phased haplotypes. *Pst* DK02d_12 haplotypes (red) and *Psh* NP85002 haplotypes (blue) are placed relative to reference *Puccinia striiformis* isolates. Bootstrap support is indicated by node circle size. *Puccinia graminis* f. sp. *tritici* isolate Pgt_210 (green) serves as the outgroup.

### Genome-wide genealogy and sequence divergence show differentiation without sorting by *forma specialis*

To place the parental isolates in a broader genomic context, we inferred a genome-wide genealogy from single-copy orthologues using each phased haplotype. The resulting genealogy showed well-supported groupings within the *P. striiformis* complex that did not correspond to *forma specialis*: both *Psh* NP85002 haplotypes grouped with *Pst*104E hap1, whereas both *Pst* DK02d_12 haplotypes grouped with *Pst*104E hap2 and AZ2 (Fig. 1b). Whole-genome comparisons across all four pairwise combinations of the phased *Pst* and *Psh* haplotypes showed sequence differentiation, with one-to-one genome alignments yielding a nucleotide identity of 98.31–98.35% and 10.46–10.69 SNPs kb⁻¹ across aligned regions (Table S3). Within-isolate divergence was lower, with the two DK02d_12 haplotypes showing 99.58% identity and 2.42 SNPs kb⁻¹ and the two NP85002 haplotypes showing 99.05% identity and 5.88 SNPs kb⁻¹.

Previous comparative genomic analysis of an independent *Pst*–*Psh* isolate pair also found higher between-isolate than within-isolate SNP density, although absolute estimates are not directly comparable because different analytical approaches were used (Xia et al. 2018b). Together, these results establish clear sequence differentiation between the tested wheat- and barley-adapted parents that exceeds within-isolate haplotype divergence, while the genome-wide genealogy shows that haplotypes do not partition according to host specialization.

### Reciprocal crosses on *Berberis vulgaris* produce viable *Pst* × *Psh* progeny

Despite contrasting host-specialized infection phenotypes on wheat and barley, reciprocal crosses between *Pst* and *Psh* on the shared sexual host *B. vulgaris* produced viable aeciospores. This sexual compatibility is particularly notable given a previous phylogenomic estimate placing the divergence of wheat- and barley-adapted *P. striiformis* forms at approximately 8.12 million years ago (Xia et al. 2018a). A total of 18 aeciospore-derived progeny isolates were recovered and maintained through multiple asexual generations on wheat and/or barley (Table S4). These progeny represent first-generation (F1) dikaryotic hybrids formed by the combination of *Pst*- and *Psh*-derived haploid mating products at the pycnial stage. Although more progeny were recovered from *Pst* → *Psh* than from *Psh* → *Pst* crosses, the limited number of progeny precludes firm conclusions about directional asymmetry or potential cytoplasmic effects. Together, these results demonstrate that intrinsic reproductive isolation is incomplete between the tested parental isolates when mating occurs on a shared alternate host. However, the ability of the F1 progeny to complete a subsequent sexual cycle was not assessed, and their fertility and potential for genetic exchange in subsequent generations remain to be tested.

Sexual reproduction has previously been demonstrated independently within *Pst* and *Psh* on *Berberis* spp. (Jin et al. 2010; Huang et al. 2019; Du et al. 2022), while population-level surveys have reported patterns consistent with occasional admixture/recombination between wheat- and barley-associated *P. striiformis formae speciales* (Du et al. 2024). The controlled crosses reported here extend these observations by providing direct experimental evidence of sexual compatibility between wheat- and barley-adapted *P. striiformis* forms.

### HD and PR mating-type loci are unlinked, with HD allele lineages spanning *forma specialis* boundaries

Chromosome-scale parental assemblies revealed physically unlinked HD and PR MAT loci consistent with a tetrapolar mating system. In both parents, the HD locus localized to chromosome 2, whereas the pheromone-receptor (PR) locus (*STE3* and *mfa*) localized to chromosome 6 (Fig. 2a). This physical separation is consistent with the unlinked HD and PR loci that define tetrapolar mating in basidiomycetes and with MAT organization reported across cereal rust fungi (Cuomo et al. 2017; Holden et al. 2023; Henningsen et al. 2024; Luo et al. 2024).

**Figure 2.**
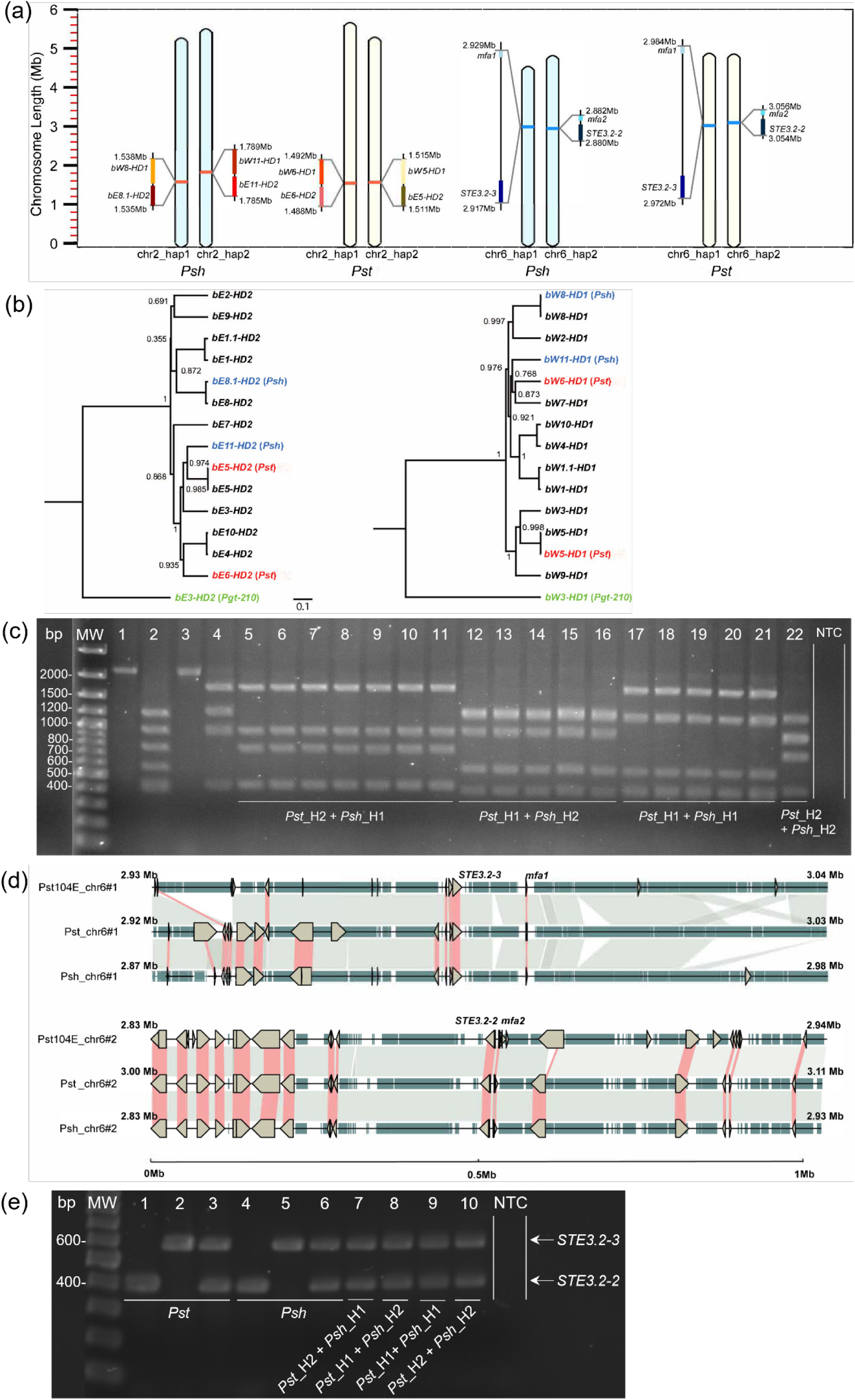
Sexual compatibility between host-adapted stripe rust forms is supported by biparental mating-type inheritance and unlinked mating-type (MAT) loci. **(a)** Chromosomal locations of mating-type loci in the parental chromosome-scale assemblies (haplotype 1 and haplotype 2). In both parents, the homeodomain (HD) locus (*bW-HD1/bE-HD2*) localized to chromosome 2, whereas the pheromone-receptor (PR) locus, containing *STE3* and *mfa*, localized to chromosome 6, consistent with physically unlinked MAT loci characteristic of tetrapolar mating. **(b)** Genealogical placement of HD mating-type alleles from *Pst* DK02d_12 (red) and *Psh* NP85002 (blue), shown separately for *bE-HD2* (left) and *bW-HD1* (right), relative to previously characterized *Puccinia striiformis* HD alleles (black). *Puccinia graminis* f. sp. *tritici* (green) is included as an outgroup. Parental alleles cluster within established HD allele lineages, showing that HD allele relationships do not partition according to *forma specialis*. Node support values are shown as posterior probabilities. Allele numbering follows previously characterized HD alleles from Holden et al. (2023), with decimal suffixes used here to distinguish closely related allele subvariants (e.g., *bW1.1* and *bE1.1*). **(c)** PCR–restriction digest genotyping of the HD mating-type locus in both parents and in sexual progeny. Lanes 1–2 show *Pst* DK02d_12 (undigested and digested), and lanes 3– 4 show *Psh* NP85002 (undigested and digested). The *Pst* parent carries Pst_H1 (1217, 576, 411 bp) and Pst_H2 (982, 775, 447 bp), whereas the *Psh* parent carries Psh_H1 (1729, 439 bp) and Psh_H2 (1249, 990 bp). In the *Pst* parental digest, the 411-bp and 447- bp fragments co-migrate, appearing as an intensified band. Progeny segregated into four genotype classes, representing all possible combinations of one *Pst*-derived and one *Psh*-derived HD haplotype following mating at the pycnial stage. These are shown in gel order: lanes 5–11 (Pst_H2 + Psh_H1), lanes 12–16 (Pst_H1 + Psh_H2), lanes 17–21 (Pst_H1 + Psh_H1), and lane 22 (Pst_H2 + Psh_H2), consistent with biparental inheritance and meiotic segregation at the HD locus (n = 18 progeny). MW, molecular weight marker (bp); NTC, no-template control. **(d)** The pheromone/receptor (PR) locus retains conserved organization across *Puccinia striiformis* haplotypes despite structural divergence in the surrounding transposable element (TE)-rich flanking regions. Synteny plots show the genomic regions surrounding the PR locus on chromosome 6 in *P. striiformis* f. sp. *tritici* (*Pst*) and *P. striiformis* f. sp. *hordei* (*Psh*) haplotypes. Brown arrows denote annotated genes, with arrow direction indicating transcriptional orientation. Green features indicate annotated TEs. Pink ribbons connect homologous syntenic blocks between haplotypes based on nucleotide alignments. The predicted pheromone-receptor genes (*STE3.2-2* and *STE3.2-3*) and adjacent pheromone precursor genes (*mfa1*, *mfa2*) are labelled. **(e)** PCR marker assay targeting the pheromone-receptor (PR) locus in parental isolates and representative sexual progeny. Locus-specific primers for *STE3.2-2* and *STE3.2-3* produced the expected 400 bp and 600 bp amplicons, respectively. In the parental isolates, *Pst* DK02d_12 was analysed by simplex PCR for each marker (lanes 1–2), followed by multiplex PCR (lane 3); *Psh* NP85002 was analysed similarly in lanes 4–6. Multiplex PCR was subsequently applied to four representative progeny (Psh/Pst_1, Pst/Psh_8, Psh/Pst_13, and Psh/Pst_18; lanes 7–10), each selected from one of the four HD segregation groups (Fig. 2c), resolving both *STE3*-associated markers in all cases and supporting the presence of both PR-associated markers in representative progeny classes. MW, molecular weight marker (bp); NTC, no-template control.

Genealogical analysis of HD alleles showed that HD mating-type variation did not partition according to *forma specialis*. The *Pst bW-HD1* and *bE-HD2* alleles grouped within previously characterized *Pst* HD allele lineages, while the *Psh* NP85002 alleles fell within the same broader lineages (Fig. 2b). Thus, HD allele relationships did not mirror the whole-genome sequence differentiation observed between the parental isolates. Given that multiallelic HD loci can be maintained by negative frequency-dependent selection (Nieuwenhuis et al. 2013; Coelho et al. 2017; Luo et al. 2024), this lack of sorting by *forma specialis* is consistent with long-term maintenance of HD allelic diversity despite host specialization.

### Biparental HD inheritance in all progeny is consistent with outcrossing and meiotic segregation

Restriction digest genotyping of the HD MAT locus supported biparental inheritance in all recovered progeny (Fig. 2c, Table S5, Fig. S1). The PCR–restriction digest assay generated distinct haplotype profiles for each dikaryotic parent. In each dikaryotic parent, H1 and H2 denote the two phased haploid nuclear genomes, each carrying a distinct HD allele combination: *Pst*_H1 = *bW5-HD1/bE5-HD2*; *Pst*_H2 = *bW6-HD1/bE6-HD2*; *Psh*_H1 = *bW11-HD1/bE11-HD2* and, *Psh*_H2 = *bW8-HD1/bE8.1-HD2*. The *Pst* parent (DK02d_12) carried two HD haplotypes, *Pst*_H1 (1217, 576, 411 bp) and *Pst*_H2 (982, 775, 447 bp). The 411-bp and 447-bp fragments co-migrated as a single intensified band due to their similar sizes. The *Psh* parent (NP85002) carried two alternative haplotypes, *Psh*_H1 (1729, 439 bp) and *Psh*_H2 (1249, 990 bp). All 18 progeny isolates carried restriction patterns consistent with inheritance of exactly one HD haplotype from each parent and segregated into four genotype classes representing all parental haplotype combinations (*Pst*_H2 + *Psh*_H1; *Pst*_H1 + *Psh*_H2; *Pst*_H1 + *Psh*_H1; *Pst*_H2 + *Psh*_H2) (Fig. 2c). Recovery of all four haplotype combinations is consistent with meiotic segregation of parental HD haplotypes and supports that the progeny derived from an outcross between the wheat- and barley-adapted parents. The observed frequencies of the four HD haplotype classes did not deviate significantly from the expectation of equal segregation under an exact multinomial goodness-of-fit test (*P* = 0.19).

Analysis of the PR locus provided additional support for conservation of loci associated with mating compatibility. Synteny comparisons across chromosome 6 haplotypes showed that the genomic regions flanking the PR locus were structurally divergent and enriched in TEs, whereas the *STE3* receptor genes and adjacent *mfa* genes retained conserved positional organization (Fig. 2d). Direct comparison of PR-associated coding sequences showed high nucleotide conservation across the aligned *STE3.2-2, STE3.2-3, mfa1* and *mfa2* sequences in *Pst* and *Psh* chromosome 6 haplotypes (Fig. S2). Together, the conserved positional organization and high nucleotide similarity of PR-associated genes indicate conservation of this component of the MAT system between the parental isolates.

PCR-based marker assays targeting the PR locus further supported the presence of both PR-associated markers in sexual progeny (Fig. 2e; Fig. S3). Locus-specific primers for *STE3.2-2* and *STE3.2-3* generated the expected 400 bp and 600 bp amplicons, respectively, and were validated in the parental isolates by simplex and multiplex PCR. Multiplex amplification of four representative progeny, each selected from one of the four HD segregation classes (Fig. 2c), resolved both *STE3*-associated markers in each case. Screening of the full progeny set showed consistent amplification of both markers across isolates (Fig. S3), confirming the presence of both PR-associated markers throughout the progeny population.

Together, the unlinked chromosomal positions of the HD and PR loci, segregation of parental HD haplotypes in sexual progeny, and conservation of PR-associated loci support a conserved MAT organization consistent with a tetrapolar mating system in the tested parental combination. Combined with successful sexual reproduction, these results show that host specialization has not been accompanied by complete loss of mating compatibility between the tested parental isolates.

### Progeny display recombinant host-interaction phenotypes, including dual-host compatibility

The two parental isolates retained strict host specialization during the uredinial stage, with wheat-adapted *Pst* infecting wheat but not barley and barley-adapted *Psh* showing the opposite pattern across the tested host panel (Fig. 3; Fig. S4; Table S6). Phenotyping across a diverse panel of wheat and barley genotypes showed that sexual progeny displayed recombinant host-interaction phenotypes and, in multiple cases, compatibility across both host backgrounds (Fig. 3a; Fig. S4). Two progeny could not be phenotyped because of insufficient spore production (Pst/Psh_6 and Psh/Pst_18). None of the progeny reproduced the complete phenotypic profile of either parent (Fig. 3a; Fig. S4; Table S7). On wheat, progeny were able to infect a subset of the wheat genotypes infected by the wheat-adapted *Pst* parent, while remaining incompatible on others. Conversely, despite the incompatibility of the wheat-adapted parent on barley, progeny were able to infect barley genotypes but showed variable infection responses across host lines. Sexual progeny isolate Pst/Psh_7 showed compatible infection on both host backgrounds, demonstrating dual-host compatibility in the tested panel (Fig. 3b). This broadened host compatibility remained constrained by host genotype, indicating that resistance gene-specific recognition continued to shape infection outcomes within each host background.

**Figure 3.**
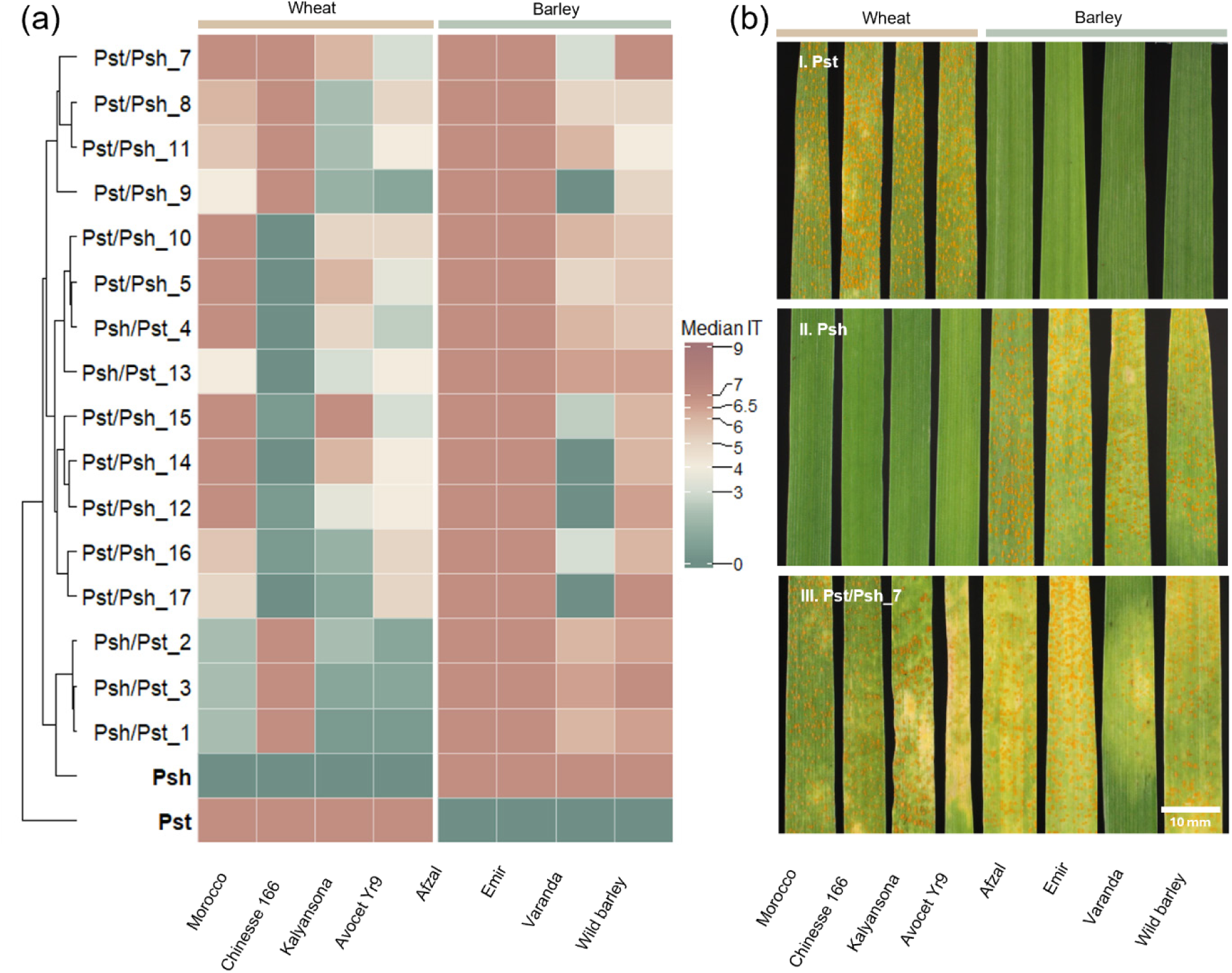
Sexual progeny from reciprocal crosses between wheat-adapted *Puccinia striiformis* f. sp. *tritici* (*Pst*; isolate DK02d_12) and barley-adapted *P. striiformis* f. sp. *hordei* (*Psh*; isolate NP85002) exhibit recombinant host-interaction phenotypes and dual-host compatibility on selected wheat and barley genotypes. **(a)** Heatmap of infection outcomes for the two parental isolates and selected sexual progeny across a subset of wheat and barley host genotypes chosen to highlight contrasting parental responses and recombinant host phenotypes in the progeny. Rows represent isolates and columns represent host genotypes. Cell colors indicate median infection type (IT) scores on the 0–9 scale. Individual IT scores ≥ 7 were classified as compatible and IT scores ≤ 6 as incompatible, with IT 4–6 representing intermediate reactions relative to strongly incompatible reactions (IT ≤ 3). For heatmap classification, median IT values ≥ 6.5 were considered compatible. The parental isolates retain strict host specialization on their respective cereal hosts, whereas the progeny display diverse recombinant phenotypes, including individual progeny compatible with genotypes from both wheat and barley. **(b)** Representative infection phenotypes on wheat and barley leaves illustrating parental host specialization and progeny dual-host compatibility. Wheat-adapted *Pst* shows compatible infection on wheat but an incompatible reaction on barley, whereas barley-adapted *Psh* shows the opposite pattern. The sexual progeny isolate Pst/Psh_7 shows compatible infection on both host backgrounds. Morocco and Afzal were used as susceptible wheat and barley controls, respectively. Images were taken 18 days post-inoculation. Scale bar = 10 mm.

Because avirulence profiles vary among isolates and reflect multiple host-interaction loci, recombinant phenotypes are expected to depend on the particular parental genetic combinations that mate on the sexual host (Ali et al. 2017; Du et al. 2024). In the progeny examined here, the combination of *Pst*- and *Psh*-derived nuclear genomes may bring together different host-interaction loci, including effector genes associated with avirulence, thereby contributing to the observed compatibility patterns across wheat and barley without producing generalized virulence. The host panel included differential lines carrying characterized resistance genes, so incompatible reactions in progeny should not be interpreted solely as generalized hybrid underperformance; they may also reflect segregating avirulence profiles interacting with host resistance backgrounds. At the same time, progeny frequently showed reduced virulence relative to parental isolates on their respective specialized hosts, consistent with earlier cereal rust crossing studies in which recombinant progeny often showed intermediate or attenuated pathogenicity (Fig. 3b; Fig. S4) (reviewed by Anikster 1984).

The observed progeny phenotypes suggest that host specialization may be shaped by variation in a subset of host-interaction determinants, including avirulence effectors recognized on defined resistance backgrounds (Schulze-Lefert and Panstruga 2011; Dracatos et al. 2018; Müller et al. 2021). Sexual hybridization can therefore combine distinct parental sets of host-interaction determinants within the same progeny, contributing to the host-compatibility profiles observed here. Future progeny resequencing and infection-stage expression profiling could link inherited genomic variation to the dual-host compatibility phenotypes observed here. Thus, the separation of sexual reproduction from primary-host infection may permit intrinsic reproductive isolation to remain incomplete while host-associated differentiation accumulates in *P. striiformis*.

## Materials and Methods

### Fungal Material and Parental Isolate Selection

Parental isolates DK02d_12 (*Pst* genetic group S8), collected from wheat in Denmark, and NP85002 (*Psh*), collected from barley in Nepal, were selected from the Global Rust Reference Center (GRRC, Denmark) collection based on contrasting host specificity (wheat-adapted *Pst* vs. barley-adapted *Psh*) (Ali et al. 2017; Thach et al. 2025). Urediniospores were propagated on susceptible cultivars Morocco (wheat) and Afzal (barley) under controlled greenhouse conditions (16 h photoperiod, 18°C/12°C day/night, 70-80% RH) and cryopreserved in liquid nitrogen for long-term storage.

### Teliospore Production and Sexual Cross Procedures

Teliospores were induced on adult plants (flag leaf fully emerged) of Morocco and Afzal grown in 15 cm pots containing Pindstrup substrate (Pindstrup Mosebrug A/S, Denmark). Urediniospores were suspended in Novec™ 7100 Engineered Fluid (3M) and applied to 1 cm² areas on flag leaves and flag-1 leaves using an airbrush spray gun. Inoculated plants were incubated at 100% relative humidity (RH) and 10-12°C in darkness for 24 h, then transferred to spore-proof greenhouse cabins and kept as described above. Leaves bearing telia were harvested approximately 8 weeks after inoculation at the onset of leaf senescence and stored in paper bags in desiccators at 5°C.

*Berberis vulgaris* (common barberry) plants were grown from seeds following stratification for two months at 5°C in darkness. Seedlings were maintained in greenhouse conditions (18-20°C, 16 h photoperiod) for 4 months to ensure adequate fresh foliage for infection. Teliospores were activated by washing leaf segments under running water followed by incubation in water at room temperature (20-25°C) in darkness for 2 days. Activated leaf segments were transferred to 2.5% water agar plates containing 50 mg l⁻¹ chloramphenicol and incubated at 13°C in darkness. After basidiospore formation was confirmed microscopically (typically 48 h), agar plates were inverted over barberry seedlings. Plants were maintained at 100% RH for 3 days at 13°C in darkness, then transferred to spore-proof greenhouse cabins and kept under the conditions described above. The two parental isolates were maintained in separate cabins throughout to prevent unintended cross-hybridization. Once pycnia matured (7-10 days after inoculation), reciprocal crosses were performed using a fine art brush (size 00) to transfer pycniospore-containing nectar droplets between plants infected with different parental isolates (*Pst* → *Psh* and *Psh* → *Pst*). Barberry leaves bearing mature aecia were harvested and positioned over seedlings of both parental hosts (Morocco and Afzal) to capture aeciospores during natural cup dehiscence. Inoculated seedlings were incubated in a dew chamber at 12°C in darkness for 48 h, with host plants replaced after 24 h to maximize aeciospore capture efficiency. Plants were then transferred to spore-proof greenhouse cabins under the conditions described above. Individual, spatially separated uredinia (putatively arising from single infection events) were excised from leaves, propagated separately on the host from which they were recovered (wheat or barley), and designated as distinct progeny isolates. Urediniospores from each progeny line were harvested, desiccated, and cryopreserved in liquid nitrogen. Dried leaf segments containing individual lesions were stored at -80°C for DNA extraction.

### DNA Extraction and Mating-Type Genotyping

Dried leaf segments containing individual uredinia from each progeny isolate and from both parents were ground with 2 mm steel beads in a Geno/Grinder® 2010 (SPEX SamplePrep, USA) at 1500 rpm for 3 min. Genomic DNA was extracted using the sbeadex™ mini plant kit (LGC Genomics) on a KingFisher™ automated system (Thermo Scientific) following Thach et al. (2025). High molecular weight DNA for long-read sequencing was extracted from ∼300 mg of fresh urediniospores following Schwessinger and Rathjen (2017), as previously described for the *Psh* isolate NP85002 (Tam et al. 2026). The same extraction workflow was applied here to the *Pst* parent DK02d_12.

### Long-read and Hi-C sequencing of parental isolates

Long-read and Hi-C datasets for the *Psh* parent (NP85002) and its corresponding haplotype-resolved assembly were published previously (Tam et al. 2026). Here, the *Pst* parent (DK02d_12) was sequenced and assembled using the same long-read and Hi-C strategy. In brief, the Oxford Nanopore Technologies (ONT) libraries were prepared using the Ligation Sequencing Kit v14 (SQK-LSK114) and sequenced on a PromethION platform using an R10.4.1 flow cell with Q20+ chemistry. Raw signals were basecalled with Dorado v0.7.2 using the super-accuracy model dna_r10.4.1_e8.2_400bps_sup@v5.0.0 to generate simplex reads. For Hi-C, fresh urediniospores (∼100 mg) were crosslinked in 1% formaldehyde for 20 min at room temperature with periodic mixing, quenched with 125 mM glycine, washed twice in 1× PBS, and pelleted following the Phase Genomics Proximo Hi-C (Fungal) guidelines. Pellets were frozen and shipped on dry ice to Phase Genomics (Seattle, WA, USA). Hi-C libraries were prepared using the Proximo Hi-C (Fungal) Kit (KT6040, protocol v4.0) with restriction enzymes DpnII, HinfI, MseI, and DdeI, and sequenced on an Illumina NovaSeq 6000 (2×150 bp).

### Haplotype-resolved genome assembly and scaffolding

The *Pst* (DK02d_12) haplotype-resolved assemblies were generated from ONT long reads with Hi-C–guided phasing using hifiasm (v0.25.0-r726) configured for ONT data and Hi-C read pairs (Cheng et al. 2021). ONT simplex reads were provided in ONT mode and Hi-C read pairs were supplied to enable haplotype phasing. Hifiasm was run in dual-scaffolding mode to improve contiguity while retaining haplotype separation by leveraging homologous regions across the alternative haplotype. Telomeric repeat handling was enabled using the canonical telomere motif (CCCTAA). This generated two phased assemblies for *Pst* DK02d_12 (haplo-type 1 and haplotype 2).

Assemblies were curated to remove non-nuclear sequences and low-confidence contigs using sequence-similarity screening and coverage-based filtering (Tam et al. 2025). Contigs were searched against the NCBI nt database using BLASTN (v2.15.0+) to identify potential contaminants and mitochondrial-derived sequences; contigs with strong similarity to mitochondrial DNA were excluded. ONT reads were then mapped back to each assembly with minimap2 (v2.28-r1209; -ax map-ont --secondary=no) and mean per-contig coverage was calculated with bamtocov (v2.7.0) (Li 2018; Birolo and Telatin 2022). Contigs shorter than 20 kb and/or supported by <10× mean coverage were removed prior to Hi-C scaffolding. Hi-C data were used to validate phasing consistency and to scaffold cleaned contigs into chromosome-scale sequences. Hi-C reads were processed with Juicer (v1.6) and scaffolding was performed with the 3D-DNA pipeline (v180922) (Durand et al. 2016b; Dudchenko et al. 2017). Draft scaffolds were manually inspected and curated using Juicebox Assembly Tools (v2.20.00) (Durand et al. 2016a). Telomeric completeness was evaluated by searching the first and last 50 bp of each scaffold for TTAGGG/CCCTAA repeats using FindTelomeres.py (https://github.com/Ja-naSperschneider/FindTelomeres). Assembly summary statistics were computed with seqkit v2.10.0 (Shen et al. 2016). Consensus quality and k-mer completeness were assessed with Merqury v1.3 using a k-mer database (k = 21) derived from ONT reads (Rhie et al. 2020). Gene-space completeness was evaluated with BUSCO v6.0.0 using the basidiomycota_odb12 dataset (Tegenfeldt et al. 2025). HiC-Pro v3.1.0 was used to generate contact matrices from MAPQ ≥ 20 Hi-C alignments and to quantify within-versus between-haplotype contact links to assess phasing consistency (Servant et al. 2015).

### Identification of mating-type genes in parental isolates

To identify MAT loci in the parental assemblies, sequences of *bW-HD1/bE-HD2* and *STE3* alleles from published *P. striiformis* datasets were used as queries in BLAST searches (BLASTN v2.15.0) against the parental genomes (Holden et al. 2023). Candidate loci were extracted and curated in Geneious Prime® 2026.0.2 by mapping reference coding sequences to assemblies and manually inspecting open reading frames (Luo et al. 2024). Pheromone receptor (*STE3/Pra*) and pheromone precursor (*mfa*) genes were extracted using the same approach as described previously (Luo et al. 2024). Diagnostic sequence differences between parental HD alleles were used to design a PCR–restriction digest assay for progeny genotyping. Chromosome ideograms were generated with Matplotlib v3.10.8 (Hunter 2007). To assess syntenic relationships between PR loci, protein sequences from the Pst104E reference isolate, *Pst* (DK02d_12), and *Psh* (NP85002) were compared using BLASTP v2.10.0 and matches with less than 70% identity were removed. Transposable element (TE) coordinates in the parental genomes were annotated using the REPETv4 snakemake pipeline as feature tracks for PR locus visualization (Bardet et al. 2025). Nucleotide sequences of chromosome 6 were compared using nucmer v3.1, retaining only alignments > 5 kb and > 90% sequence identity. Synteny plots of the 50 kb regions surrounding the PR loci were generated in R using gggenomes v1.2.2 (Hackl 2024). To compare PR-associated coding sequences between *Pst* DK02d_12 and *Psh* NP85002, the two *STE3* coding sequences*, STE3.2-2* and *STE3.2-3,* from the *Pst*104E reference isolate were mapped to both chromosome 6 haplotypes of *Pst* and *Psh* using the “Map to Reference” function in Geneious Prime® 2026.0.2. To assess conservation of the adjacent *mfa* sequences, chromosome 6 haplotypes were scanned in Geneious using the “Live Annotate & Predict” function with previously described wheat rust *mfa* sequences from Cuomo et al. (2017) as references. Sequence identity and conservation were then inspected across the corresponding *STE3* and *mfa* coding regions.

### Mating-type genotyping by HD restriction assay and *STE3* marker PCR

To verify progeny origin and assign parental contributions at the MAT locus, we genotyped the HD locus using a PCR–restriction digest assay designed from parental HD allele sequences. Degenerate primers were designed against conserved regions of HD genes shared between *Pst* and *Psh* using Geneious Prime® 2026.0.2 (Primer3 implementation): MAT-F 5′-CGAAAAGTCTACTAGSAAATGTC-3′ and MAT-R 5′-GGGTTCATATTGCATAWCAAAGGV-3′ (Eurofins, Europe). PCR was performed on an Applied Biosystems 2720 Thermal Cycler with an initial denaturation at 94°C for 2 min 30 s; 35 cycles of 94°C for 30 s, 57°C for 60 s, and 72°C for 30 s; followed by a final extension at 72°C for 5 min and a hold at 4°C. Amplicons were purified using the GenJET PCR Purification Kit (Thermo Scientific, #K0701) and digested with KpnI-HF and DraI restriction enzymes (New England Biolabs, UK) according to the manufacturer’s instructions. Digestion products were separated on 1.7% agarose gels (100 V, 45 min) and visualized by ethidium bromide staining. Deviation from the expected 1:1:1:1 segregation ratio of HD haplotype classes was assessed using an exact multinomial goodness-of-fit test implemented in R v4.5.2 (package *XNomial*, v1.0.4.1).

To further assess mating-type-associated markers, *STE3.2-2* and *STE3.2-3* loci were targeted using locus-specific primers designed from the parental genome assemblies in Geneious Prime® 2026.0.2. Primers were designed to specifically amplify each *STE3* locus while avoiding cross-amplification of the alternate *STE3* locus. Primers were selected based on absence of predicted off-target sites and distinct amplicon sizes. The selected primer pair for *STE3.2-2* was *STE3.2-2*_F (5′-AACCCACTCGTCTGGTAAACC-3′) and *STE3.2-2*_R (5′-GGCAC-CGTTACAATCCTTGAAG-3′), yielding an expected amplicon of 400 bp. The selected primer pair for *STE3.2-3* was *STE3.2-3*_F (5′-TTGACTCCTGATCGATGCGTC-3′) and *STE3.2-3*_R (5′-CTTTGTCAGTGGCACGATTCTG-3′), yielding an expected amplicon of 600 bp. Primer specificity was validated in silico using Geneious Prime® ’Test with Saved Primers’ function to confirm locus-specific amplification. *STE3.2-2* and *STE3.2-3* were amplified in multiplex PCR using the same cycling profile as described above, except with an annealing temperature of 60°C, and products were separated on 1.7% agarose gels (100 V, 45 min).

### Genome-wide orthology and HD allele genealogies

To place the parental isolates in a broader genealogical context and to relate genome-wide relationships to mating-type variation, we inferred phylogenies using (i) proteome-wide orthology and (ii) targeted HD allele sequences. For the genome-wide genealogy, orthology inference was performed with OrthoFinder v2.5.5 using predicted proteomes from phased haploid genomes of four *P. striiformis* isolates: *Pst* DK02d_12 (this study), *Pst*104E (Tam et al. 2025), AZ2 (Wang et al. 2024), and *Psh* NP85002 (Tam et al. 2026). *Pgt_*210 (Li et al. 2019) was included as an outgroup (Emms and Kelly 2015, 2019). For *Pst* DK02d_12, predicted gene models were generated by lifting over the *Pst*104E and AZ2 annotations onto each phased haplotype using Liftoff v1.6.3 with the -chrom option (Shumate and Salzberg, 2021; Tam et al. 2025). OrthoFinder was run using BLASTP similarity searches (BLAST+ v2.12.0) and the MSA-based species tree workflow (-M msa), which aligns single-copy orthologues with MAFFT and builds a concatenated supermatrix for species-tree inference. A maximum-likelihood tree and branch support were inferred with IQ-TREE (v3.0.1) using ModelFinder model selection, with branch support estimated using ultrafast bootstrap. The resulting tree was visualized in iTOL (Letunic and Bork 2024). Genome-wide sequence similarity between the phased parental assemblies was assessed using dnadiff from MUMmer v4.0.0rc1 (Kurtz et al. 2004; Marçais et al. 2018). All four pairwise combinations of the two *Pst* DK02d_12 and two *Psh* NP85002 haplotypes were compared using default settings. Nucleotide identity and aligned sequence length were obtained from the one-to-one genome alignment blocks reported by dnadiff. SNP density was calculated as the total number of SNPs divided by the one-to-one aligned sequence length and expressed as SNPs kb⁻¹.

For the HD allele genealogy, coding sequences of HD alleles from the parental isolates were aligned with published *Pst* HD alleles and with *bW3-HD1/bE3-HD2* from *Pgt* isolate 210 as an outgroup. Alignments were generated with MAFFT v7.490 (Katoh and Standley 2013) and trimmed with trimAl v1.2rev59 (Capella-Gutiérrez et al. 2009) to remove sites with gaps in >10% of sequences. Pairwise nucleotide differences among closely related HD coding sequences were additionally assessed from MAFFT alignments in Geneious Prime® 2026.0.2. Decimal suffixes (e.g., bE8.1) distinguish closely related HD allele subvariants, while closely related sequences without a subvariant designation retain the same allele name. Substitution model selection was performed with bModelTest v1.2.1, which supported TN93+G for both *bW-HD1* and *bE-HD2* alignments (Bouckaert and Drummond 2017). Inference was carried out in BEAST v2.7.8 (Bouckaert et al. 2019), summarized as maximum clade credibility trees using TreeAnnotator v2.7.6 (Drummond and Rambaut 2007), and visualized in FigTree v1.4.4 (https://tree.bio.ed.ac.uk/software/figtree/).

### Host range assessment and virulence phenotyping

Progeny isolates representing novel HD locus genotypes, together with both parental isolates, were inoculated on a host panel including (i) wheat (*Yr1*, *Yr2*, *Yr3*, *Yr6*, *Yr7*, *Yr8*, *Yr9*, *Yr15*, *Yr17*, *Yr24*, *Yr25*, *Yr27*, *Yr32*, *YrSp, YrAvS*) and barley (*rpsEm1*, *rpsEm2*, *rps2*, *RpsBa*, *Rps4*, *rpsHF*, *rpsTr1*, *rpsTr2*, *Rps3*, *rpsI5*, *rpsHi1*, *rpsHi2*, *Rps1.b*, *rpsAst*, *rpsVa1*, *rpsVa2*) differential lines carrying characterized resistance genes, (ii) susceptible controls (wheat cv. Morocco; barley cv. Afzal), and (iii) additional grass host relatives (*Aegilops tauschii*, *Triticum monococcum*, *T. dicoccum*, *Elymus repens*, and *Hordeum spontaneum*) (Chen and Line 2003; Thach et al. 2025) (Table S1). For each host line, 6–8 seeds were sown in peat-based substrate in 10 cm pots and maintained under controlled greenhouse conditions (18°C/12°C day/night, 70–80% RH, 16 h photoperiod). Seedlings were inoculated at the one-leaf stage, when the first leaf was fully expanded and the second leaf was half-emerged (approximately 10–12 days after sowing). Inoculum consisted of 20–30 mg urediniospores per isolate suspended in Novec™ 7100 (3M) and applied uniformly using an airbrush spray gun (Hovmøller et al. 2017; Thach et al. 2025). Inoculated plants were incubated at 100% relative humidity for 24 h at 12°C in darkness and then transferred to spore-proof greenhouse cabins under the conditions described above. Infection types (IT) were scored visually 18 days post-inoculation on the first and second leaves using a 0–9 scale (Hovmøller et al. 2017). IT ≤ 6 was considered incompatible and IT ≥ 7 compatible on the 0–9 scale. To capture quantitative differences among incompatible reactions, IT values of 4–6 were classified as intermediate relative to strongly incompatible reactions (IT ≤ 3). For each isolate × host combination, the distribution of IT scores across the first and second leaves was combined to calculate a median IT value. When the median fell between two adjacent classes, the midpoint value was retained (e.g., 6.5); median IT values ≥ 6.5 were considered compatible. Median IT values were visualized as heatmaps in R v4.5.2 using the ComplexHeatmap package v2.26.1 (Gu et al. 2016). Heatmaps were generated from isolate × host median IT matrices and isolates were hierarchically clustered to summarize similarity in host interaction phenotypes.

## Data availability

ONT long reads and Hi-C reads generated for *Pst* isolate DK02d_12 have been deposited in the NCBI Sequence Read Archive (SRA) under BioProject PRJNA1453228. The haplotype-resolved nuclear genome assemblies are publicly available in NCBI under accessions JBX-OVE000000000 (haplotype 1) and JBXOVF000000000 (haplotype 2). The assemblies and associated resources, including tree files, mating-type locus coding sequences, and coordinate files containing gene and transposable element features used as annotation tracks for PR locus synteny visualization, are available via Zenodo (DOI: 10.5281/zenodo.19565706). Study-specific scripts used for data visualization and figure generation are available on GitHub: https://github.com/julianralgaba/Pst-Psh-mating-type-host-range. Assembly and genome quality assessment workflows are available in the companion repository: https://github.com/ri-tatam/PshNP85002-genome-assembly/tree/v1.0.

## Acknowledgements

We acknowledge the contribution of the Plant Pathogen ‘Omics Initiative consortium to the generation of data used in this publication. This work was supported by computational resources provided by the Australian Government through the National Computational Infrastructure (NCI) under the ANU Merit Allocation Scheme. The authors acknowledge the provision of computing and data resources provided by the Australian BioCommons Leadership Share (ABLeS) program. This program is co-funded by Bioplatforms Australia, enabled by the Commonwealth Government National Collaborative Research Infrastructure Strategy (NCRIS), the National Computational Infrastructure and Pawsey Supercomputing Research Centre (Gustafsson, et al. 2023). We thank John Rathjen (ANU) for critical review of the draft manuscript, and Ellen Jørgensen and Jakob Sørensen (GRRC, Aarhus University) for assistance with isolate handling and plant multiplication, respectively.

## Author contributions

JRA and BS designed and conceptualized the study. JRA supervised the research. SM performed sexual crosses, prepared fungal spores for Hi-C library preparation, and carried out MAT genotyping. ZL identified mating-type genes in the parental isolates and performed PR locus synteny analyses and HD allele genealogies. RT assembled and scaffolded the *Pst* DK02d_12 genome. MM and AJ extracted high-molecular-weight DNA and performed ONT long-read sequencing. JRA designed the MAT primer pairs. SM and JRA conducted host range assessment and virulence phenotyping. ZL, RT and JRA carried out data visualization. SM and JRA wrote the original draft manuscript. All authors contributed to revision of the manuscript, reviewed the final version, and approved it for publication.

---

**Figure S1.** In silico restriction digest and restriction sites of homeodomain (HD) mating-type alleles.

**Figure S2.** Nucleotide-level comparison of PR-associated coding sequences across *Puccinia striiformis* f. sp. *tritici* DK02d_12 and *P. striiformis* f. sp*. hordei* NP85002 parental isolates.

**Figure S3.** PCR assessment of pheromone-receptor markers across all sexual progeny.

**Figure S4.** Heatmap showing infection outcomes of the two parental isolates, wheat-adapted *Puccinia striiformis* f. sp. *tritici* (*Pst*; isolate DK02d_12) and barley-adapted *Puccinia striiformis* f. sp. *hordei* (*Psh*; isolate NP85002), together with the sexual progeny.

**Table S1.** Differential lines and corresponding resistance genes used for host range assessment and virulence phenotyping.

**Table S2.** Assembly statistics of the genome assemblies for *Puccinia striiformis* f. sp. *Tritici* (*Pst*), isolate DK02d_12, and *P. striiformis* f. sp. *hordei* (*Psh*), isolate NP85002.

**Table S3.** Whole-genome sequence divergence between phased *Pst* DK02d_12 and *Psh* NP85002 haplotypes based on *dnadiff* comparisons.

**Table S4.** Progeny recovered from reciprocal crosses between *Puccinia striiformis* f. sp. *tritici* (*Pst*), isolate DK02d_12, and *P. striiformis* f. sp. *hordei* (*Psh*), isolate NP85002, and the host used for initial recovery/propagation.

**Table S5.** HD (homeodomain) locus haplotype assignments for parents and sexual progeny based on restriction fragment sizes.

**Table S6.** Virulence phenotyping of progeny and parental isolates (*Pst* and *Psh*).

**Table S7.** Median infection type (IT) values used for heatmap generation.

