## Supplementary Figures for "A conserved tetrapolar mating system permits sexual hybridization and recombinant host phenotypes in wheat- and barley-adapted *Puccinia striiformis*" (from another version of this preprint: version 1, 2026-05-05)

**Figure S1. In silico restriction digest and restriction sites of homeodomain (HD) mating-type alleles.**

**(a)** Virtual PCR products spanning the HD mating-type locus were generated in Geneious Prime from the chromosome-scale assemblies of the parental isolates (*Puccinia striiformis* f. sp. *tritici* (*Pst*) and *P. striiformis* f. sp. *hordei* (*Psh*)) and digested in silico with KpnI and DraI, producing haplotype-specific fragment patterns for all four parental haplotypes.

**(b)** Geneious Prime sequence views of the corresponding haplotype-specific amplicons show the positions of KpnI and DraI restriction sites within each sequence. Exact fragment sizes are provided in Table S4.

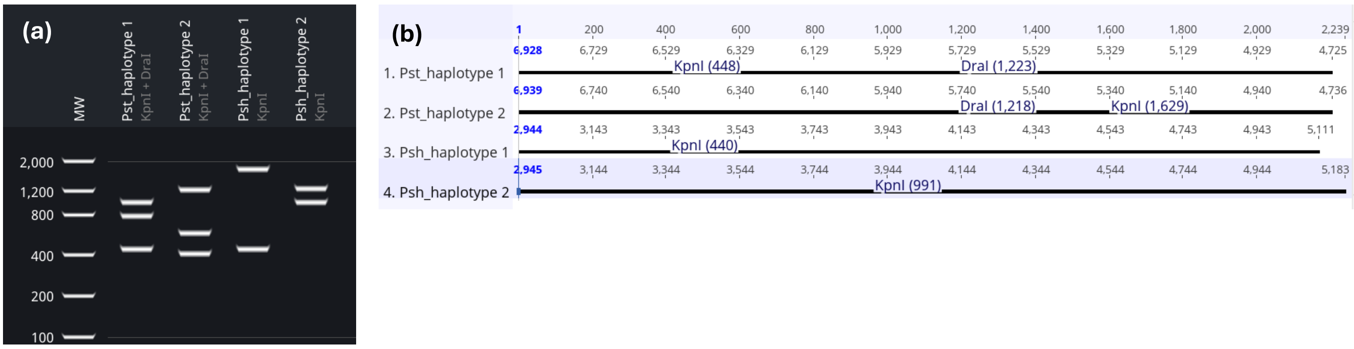

**Figure S2. Nucleotide-level comparison of PR-associated coding sequences across *Pst* DK02d_12 and *Psh* NP85002 parental isolates.**

**(a)** PR-associated *STE3* coding sequences were compared between the parental chromosome 6 haplotypes using Pst104E as a reference for mapping. *STE3.2-2* and *STE3.2-3* mapped to the expected PR loci on chr6_2 and chr6_1, respectively, in both parental isolates. Green identity tracks high indicate high nucleotide conservation across the aligned coding regions.

**(b)** Pheromone precursor genes, *mfa1* and *mfa2*, were identified on the corresponding chromosome 6 haplotypes using previously described wheat rust *mfa* sequences from Cuomo *et al.*, (2017) as references.

(a)

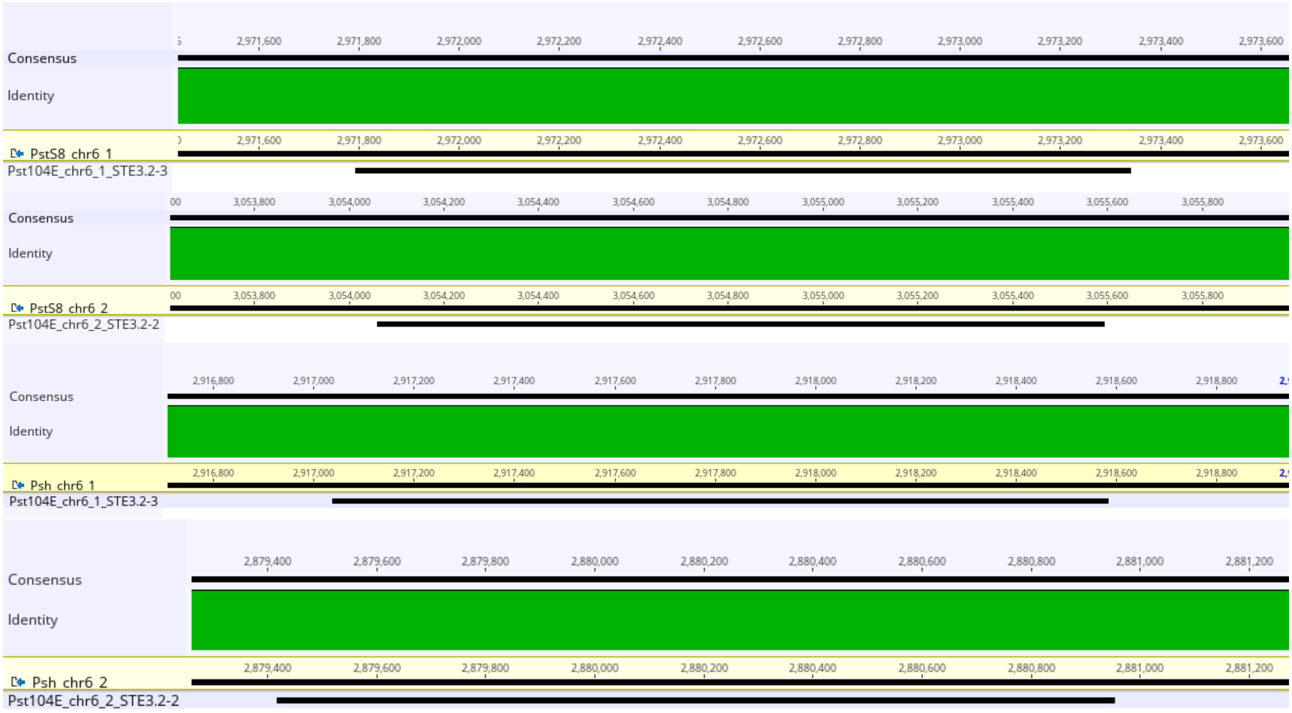


(b)

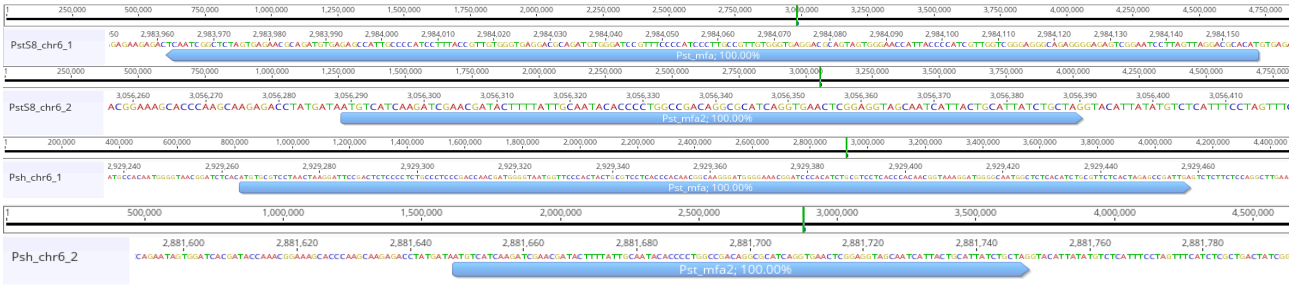

**Figure S3. PCR assessment of pheromone receptor markers across all sexual progeny.** Multiplex PCR targeting the PR locus was performed using locus-specific primers for *STE3.2-2* and *STE3.2-3*, yielding the expected 400-bp and 600-bp amplicons. In the parental isolates (*Pst* DK02d_12 and *Psh* NP85002), each marker was first validated by simplex PCR (lanes 1-2 and 4-5) and then co-amplified by multiplex PCR (lanes 3 and 6). Multiplex PCR was subsequently applied to the 18 progeny (lanes 7-24), resolving both *STE3*-associated markers across all progeny. MW, molecular weight marker (bp); NTC, no-template control.

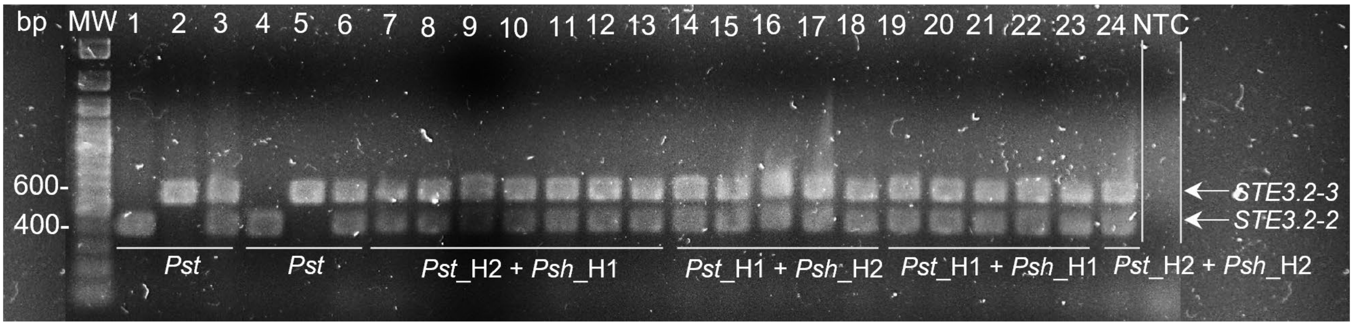

**Figure S4.** Heatmap showing infection outcomes of the two parental isolates, wheat-adapted *Puccinia striiformis* f. sp. *tritici* (*Pst*; isolate DK02d_12) and barley-adapted *Puccinia striiformis* f. sp. *hordei* (*Psh*; isolate NP85002), together with sexual progeny across the complete panel of wheat and barley host genotypes used in this study, including differential lines, near-isogenic lines, and wild relatives. Columns represent host genotypes and rows represent fungal isolates. Cell colors indicate median infection type (IT) scores on the 0–9 scale, where IT ≥ 7 indicates compatibility and IT ≤ 6 indicates non-compatibility. The parental isolates retained strict host specialization to their respective cereal hosts, whereas sexual progeny showed diverse recombinant phenotypes with compatibility on multiple wheat and barley genotypes, consistent with reshuffled host interaction phenotypes following sexual recombination.

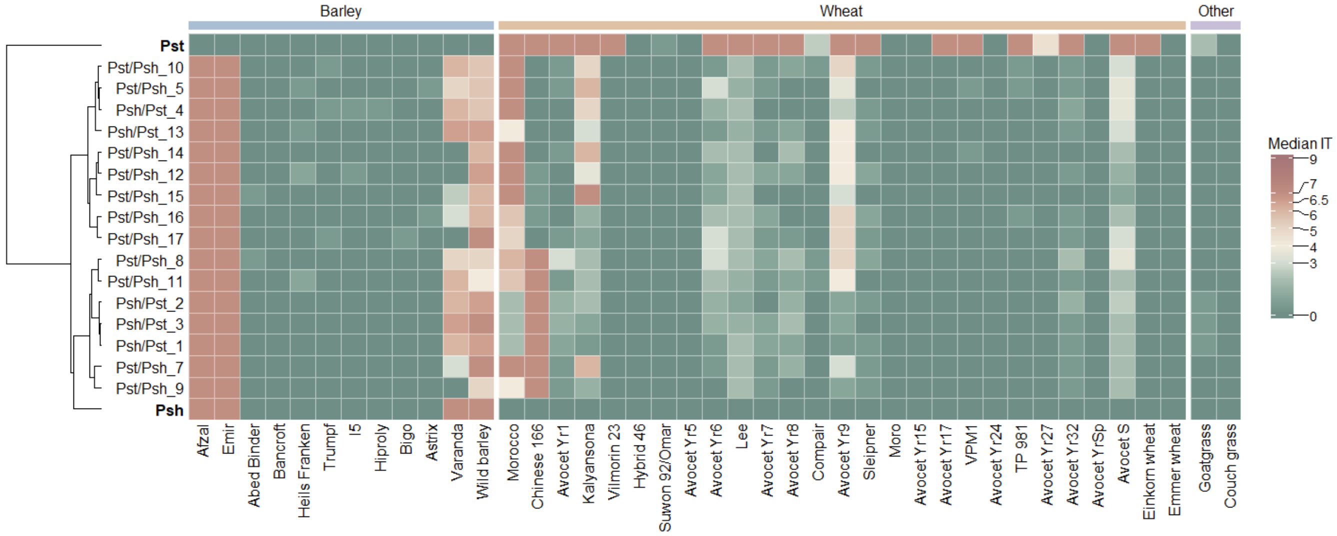
