## Supplementary Tables for "A conserved tetrapolar mating system permits sexual hybridization and recombinant host phenotypes in wheat- and barley-adapted *Puccinia striiformis*" (from another version of this preprint: version 1, 2026-05-05)

**Table S1.** Differential lines and corresponding resistance genes used for host range assessment and virulence phenotyping.

| No. | Differential line | Resistance gene(s)[a] | Host species |
|---|---|---|---|
| 1 | Afzal | none | *Hordeum vulgare* |
| 2 | Emir | *rpsEm1, rpsEm2* | *Hordeum vulgare* |
| 3 | Abed Binder | *rps2* | *Hordeum vulgare* |
| 4 | Bancroft | *RpsBa* | *Hordeum vulgare* |
| 5 | Heils Franken | *Rps4, rpsHF* | *Hordeum vulgare* |
| 6 | Trumpf | *rpsTr1, rpsTr2* | *Hordeum vulgare* |
| 7 | I5 | *Rps3, rpsI5* | *Hordeum vulgare* |
| 8 | Hiproly | *rpsHi1, rpsHi2* | *Hordeum vulgare* |
| 9 | Bigo | *Rps1.b* | *Hordeum vulgare* |
| 10 | Astrix | *Rps4, rpsAst* | *Hordeum vulgare* |
| 11 | Varanda | *rpsVa1, rpsVa2* | *Hordeum vulgare* |
| 12 | Wild barley | - | *Hordeum spontaneum* |
| 13 | Morocco | none | *Triticum aestivum* |
| 14 | Chinese 166 | *Yr1* | *Triticum aestivum* |
| 15 | Avocet Yr1 | *Yr1, Yr18, AvS* | *Triticum aestivum* |
| 16 | Kalyansona | *Yr2, +* | *Triticum aestivum* |
| 17 | Vilmorin 23 | *Yr3, Yr25, +* | *Triticum aestivum* |
| 18 | Hybrid 46 | *Yr4, +* | *Triticum aestivum* |
| 19 | Suwon 92/Omar | *Yr4, +* | *Triticum aestivum* |
| 20 | Avocet Yr5 | *Yr5, Yr18, AvS* | *Triticum aestivum* |
| 21 | Avocet Yr6 | *Yr6, AvS* | *Triticum aestivum* |
| 22 | Lee | *Yr7, +* | *Triticum aestivum* |
| 23 | Avocet Yr7 | *Yr7, AvS* | *Triticum aestivum* |
| 24 | Avocet Yr8 | *Yr8* | *Triticum aestivum* |
| 25 | Compair | *Yr8, +* | *Triticum aestivum* |
| 26 | Avocet Yr9 | *Yr9, AvS* | *Triticum aestivum* |
| 27 | Sleipner | *Yr9, +* | *Triticum aestivum* |
| 28 | Moro | *Yr10* | *Triticum aestivum* |
| 29 | Avocet Yr15 | *Yr15, AvS* | *Triticum aestivum* |
| 30 | Avocet Yr17 | *Yr17, AvS* | *Triticum aestivum* |
| 31 | VPM1 | *Yr17, +* | *Triticum aestivum* |
| 32 | Avocet Yr24 | *Yr24, AvS* | *Triticum aestivum* |
| 33 | TP 981 | *Yr25, +* | *Triticum aestivum* |
| 34 | Avocet Yr27 | *Yr27, AvS* | *Triticum aestivum* |
| 35 | Avocet Yr32 | *Yr32, AvS* | *Triticum aestivum* |
| 36 | Avocet YrSp | *YrSp, Yr18, AvS* | *Triticum aestivum* |
| 37 | Avocet S | *AvS* | *Triticum aestivum* |
| 38 | Einkorn wheat | - | *Triticum monococcum* |
| 39 | Emmer wheat | - | *Triticum dicoccon* |
| 40 | Goatgrass | - | *Aegilops tauschii* |
| 41 | Couch grass | - | *Elymus repens* |

[a]Gene designations for barley and wheat differentials are from Thach et al. (2025) and Chen & Line (2003). "+" indicates additional, uncharacterized resistance genes may be present in the differential background.

Thach T, Justesen AF, Rodriguez-Algaba J, Sørensen CK, Hovmøller MS 2025. Integrated Approach for Genotyping and Race Typing of Puccinia striiformis in Wheat. In: Brar GS, Holden S eds. Wheat Rusts and Resistance Breeding: Methods and Protocols. New York, NY: Springer US, 75-91.
Chen X, Line RF. 2003. Identification of genes for resistance to Puccinia striiformis f. sp. hordei in 18 barley genotypes. Euphytica 129(1): 127-146.

**Table S2.** Assembly statistics of the genome assemblies for *Puccinia striiformis* f. sp. *tritici* (*Pst),* isolate DK02d_12, and *Puccinia striiformis* f. sp. *hordei* (*Psh* ), isolate NP85002. Metrics are reported for the combined assembly and for each haplotype separately.

| Assembly statistics | Pst_hap1/2 | Pst_hap1 | Pst_hap2 | Psh_hap1/2[a] | Psh_hap1[a] | Psh_hap2[a] |
|---|---|---|---|---|---|---|
| Total size (bp) | 153,592,466 | 76,625,775 | 76,966,691 | 151,290,331 | 75,244,898 | 76,045,433 |
| Number of T2T chromosomes | 33 | 17 | 16 | 33 | 17 | 16 |
| Scaffold N50 (bp) | 4,684,083 | 4,705,463 | 4,619,999 | 4,580,210 | 4,519,295 | 4,580,210 |
| Longest scaffold (bp) | 5,749,073 | 5,749,073 | 5,373,267 | 5,598,305 | 5,377,700 | 5,598,305 |
| GC content (%) | 44.34 | 44.39 | 44.28 | 44.33 | 44.32 | 44.34 |
| Telomeres detected/expected | 68/72 | 35/36 | 33/36 | 69/72 | 35/36 | 34/36 |
| Number of internal gaps | 18 | 10 | 8 | 6 | 3 | 3 |

[a] Assembly stats from Tam et al., 2026.

**Table S3.** Progeny recovered from reciprocal crosses between *Puccinia striiformis* f. sp. *tritici* (*Pst*), isolate DK02d_12, and *Puccinia striiformis* f. sp. *hordei* (*Psh*), isolate NP85002, and the host used for initial recovery/propagation.

| Progeny ID | Cross direction[a] | Recovery host |
|---|---|---|
| Psh/Pst_1 | Psh → Pst | Afzal |
| Psh/Pst_2 | Psh → Pst | Afzal |
| Psh/Pst_3 | Psh → Pst | Afzal |
| Psh/Pst_4 | Psh → Pst | Afzal |
| Pst/Psh_5 | Pst → Psh | Afzal |
| Pst/Psh_6 | Pst → Psh | Afzal |
| Pst/Psh_7 | Pst → Psh | Morocco |
| Pst/Psh_8 | Pst → Psh | Afzal |
| Pst/Psh_9 | Pst → Psh | Afzal |
| Pst/Psh_10 | Pst → Psh | Afzal |
| Pst/Psh_11 | Pst → Psh | Afzal |
| Pst/Psh_12 | Pst → Psh | Morocco |
| Psh/Pst_13 | Psh → Pst | Afzal |
| Pst/Psh_14 | Pst → Psh | Morocco |
| Pst/Psh_15 | Pst → Psh | Morocco |
| Pst/Psh_16 | Pst → Psh | Afzal |
| Pst/Psh_17 | Pst → Psh | Afzal |
| Psh/Pst_18 | Psh → Pst | Morocco |

[a]Cross direction indicates the direction of fertilization during reciprocal crosses: pycniospores from the isolate at the arrow tail (donor) were transferred to pycnia produced by the isolate at the arrow head (recipient).

**Table S4**. HD (homeodomain) locus haplotype assignments for parents and sexual progeny based on restriction fragment sizes. HD haplotype names used for restriction digest genotyping correspond to the HD allele combinations shown in Fig. 2b: Pst_H1 = *bW5-HD1/bE5-HD2* ; Pst_H2 = *bW6-HD1/bE6-HD2* ; Psh_H1 = *bW11-HD1/bE11-HD2* ; Psh_H2 = *bW8-HD1/bE8-HD2* .

| Isolate | HD haplotype | Diagnostic fragment sizes (bp) |
|---|---|---|
| DK02d_12 (Pst) | Pst_H1 | 1217, 576, 411 |
| | Pst_H2 | 982, 775, 447 |
| NP85002 (Psh) | Psh_H1 | 1729, 439 |
| | Psh_H2 | 1249, 990 |

**Progeny haplotype combinations**

| Progeny ID | Fragment sizes (bp) | Assigned haplotype combination |
|---|---|---|
| Psh/Pst_1 | 1729,982,775,447,439 | Pst_H2 + Psh_H1 |
| Psh/Pst_2 | 1729,982,775,447,439 | |
| Psh/Pst_3 | 1729,982,775,447,439 | |
| Psh/Pst_4 | 1729,982,775,447,439 | |
| Pst/Psh_5 | 1729,982,775,447,439 | |
| Pst/Psh_6 | 1729,982,775,447,439 | |
| Pst/Psh_7 | 1729,982,775,447,439 | |
| Pst/Psh_8 | 1249,1217,990,576,411 | Pst_H1 + Psh_H2 |
| Pst/Psh_9 | 1249,1217,990,576,411 | |
| Pst/Psh_10 | 1249,1217,990,576,411 | |
| Pst/Psh_11 | 1249,1217,990,576,411 | |
| Pst/Psh_12 | 1249,1217,990,576,411 | |
| Psh/Pst_13 | 1729,1217,576,439,411 | Pst_H1 + Psh_H1 |
| Pst/Psh_14 | 1729,1217,576,439,411 | |
| Pst/Psh_15 | 1729,1217,576,439,411 | |
| Pst/Psh_16 | 1729,1217,576,439,411 | |
| Pst/Psh_17 | 1729,1217,576,439,411 | |
| Psh/Pst_18 | 1249,990,982,775,447 | Pst_H2 + Psh_H2 |

**Table S5.** Virulence phenotyping of progeny and parental isolates (*Pst* and *Psh*). Infection types (ITs) were scored visually 18 days post-inoculation on the first and second leaves using a 0–9 scale (Hovmøller et al., 2017). IT ≤ 6 was considered non-compatible and IT ≥ 7 compatible. To capture quantitative differences among non-compatible reactions, IT = 4–6 was classified as intermediate relative to strongly incompatible reactions (IT ≤ 3). For each isolate × host combination, IT scores from the first and second leaves were combined to calculate a median IT value. When the median fell between two adjacent classes, the midpoint value was retained (e.g. 6.5); median IT values ≥ 6.5 were classified as compatible.

| | | | | Infection Type | | | | | | | | | |
|---|---|---|---|---|---|---|---|---|---|---|---|---|---|
| **Isolate** | **Entry** | **Crop** | **Leaf** | **0** | **1** | **2** | **3** | **4** | **5** | **6** | **7** | **8** | **9** |
| Psh/Pst_1 | Afzal | Barley | 1 | | | | | | | | 6 | | |
| Psh/Pst_1 | Afzal | Barley | 2 | | | | | | | | 6 | | |
| Psh/Pst_1 | Emir | Barley | 1 | | | | | | | 3 | 3 | | |
| Psh/Pst_1 | Emir | Barley | 2 | | | | | | | | 6 | | |
| Psh/Pst_1 | Abed Binder | Barley | 1 | 8 | | | | | | | | | |
| Psh/Pst_1 | Abed Binder | Barley | 2 | 8 | | | | | | | | | |
| Psh/Pst_1 | Bancroft | Barley | 1 | 7 | | | | | | | | | |
| Psh/Pst_1 | Bancroft | Barley | 2 | 7 | | | | | | | | | |
| Psh/Pst_1 | Heils Franken | Barley | 1 | 7 | | | | | | | | | |
| Psh/Pst_1 | Heils Franken | Barley | 2 | 3 | 4 | | | | | | | | |
| Psh/Pst_1 | Trumpf | Barley | 1 | 7 | | | | | | | | | |
| Psh/Pst_1 | Trumpf | Barley | 2 | 7 | | | | | | | | | |
| Psh/Pst_1 | I5 | Barley | 1 | 7 | | | | | | | | | |
| Psh/Pst_1 | I5 | Barley | 2 | 7 | | | | | | | | | |
| Psh/Pst_1 | Hiproly | Barley | 1 | 8 | | | | | | | | | |
| Psh/Pst_1 | Hiproly | Barley | 2 | 8 | | | | | | | | | |
| Psh/Pst_1 | Bigo | Barley | 1 | 8 | | | | | | | | | |
| Psh/Pst_1 | Bigo | Barley | 2 | 8 | | | | | | | | | |
| Psh/Pst_1 | Astrix | Barley | 1 | 7 | | | | | | | | | |
| Psh/Pst_1 | Astrix | Barley | 2 | 7 | | | | | | | | | |
| Psh/Pst_1 | Varanda | Barley | 1 | | | | | | 2 | 4 | | | |
| Psh/Pst_1 | Varanda | Barley | 2 | | | | | | | 4 | 2 | | |
| Psh/Pst_1 | Hordeum spontaneum | Barley | 1 | | | | | | | 4 | 2 | | |
| Psh/Pst_1 | Hordeum spontaneum | Barley | 2 | | | | | | | 2 | 4 | | |
| Psh/Pst_1 | Aegilops tauschii | Other | 1 | 7 | | | | | | | | | |
| Psh/Pst_1 | Aegilops tauschii | Other | 2 | | 7 | | | | | | | | |
| Psh/Pst_1 | Elymus repens | Other | 1 | 7 | | | | | | | | | |
| Psh/Pst_1 | Elymus repens | Other | 2 | 7 | | | | | | | | | |
| Psh/Pst_1 | Morocco | Wheat | 1 | | 4 | 4 | | | | | | | |
| Psh/Pst_1 | Morocco | Wheat | 2 | | | 8 | | | | | | | |
| Psh/Pst_1 | Chinese 166 | Wheat | 1 | | | | | | | | 6 | | |
| Psh/Pst_1 | Chinese 166 | Wheat | 2 | | | | | | | | 6 | | |
| Psh/Pst_1 | Avocet Yr1 | Wheat | 1 | | 6 | | | | | | | | |
| Psh/Pst_1 | Avocet Yr1 | Wheat | 2 | | 6 | | | | | | | | |
| Psh/Pst_1 | Kalyansona | Wheat | 1 | 6 | | | | | | | | | |
| Psh/Pst_1 | Kalyansona | Wheat | 2 | | 3 | 3 | | | | | | | |
| Psh/Pst_1 | Vilmorin 23 | Wheat | 1 | 7 | | | | | | | | | |
| Psh/Pst_1 | Vilmorin 23 | Wheat | 2 | 7 | | | | | | | | | |

| | | | | | | | | | | | |
|---|---|---|---|---|---|---|---|---|---|---|---|
| Psh/Pst_1 | Hybrid 46 | Wheat | 1 | 8 | | | | | | | |
| Psh/Pst_1 | Hybrid 46 | Wheat | 2 | 8 | | | | | | | |
| Psh/Pst_1 | Suwon/Omar | Wheat | 1 | 8 | | | | | | | |
| Psh/Pst_1 | Suwon/Omar | Wheat | 2 | 8 | | | | | | | |
| Psh/Pst_1 | Avocet Yr5 | Wheat | 1 | 7 | | | | | | | |
| Psh/Pst_1 | Avocet Yr5 | Wheat | 2 | 7 | | | | | | | |
| Psh/Pst_1 | Avocet Yr6 | Wheat | 1 | 6 | | | | | | | |
| Psh/Pst_1 | Avocet Yr6 | Wheat | 2 | | 6 | | | | | | |
| Psh/Pst_1 | Lee | Wheat | 1 | | | 7 | | | | | |
| Psh/Pst_1 | Lee | Wheat | 2 | | | 7 | | | | | |
| Psh/Pst_1 | Avocet Yr7 | Wheat | 1 | | 6 | | | | | | |
| Psh/Pst_1 | Avocet Yr7 | Wheat | 2 | | 6 | | | | | | |
| Psh/Pst_1 | Avocet Yr8 | Wheat | 1 | | 8 | | | | | | |
| Psh/Pst_1 | Avocet Yr8 | Wheat | 2 | | 4 | 4 | | | | | |
| Psh/Pst_1 | Compair | Wheat | 1 | 8 | | | | | | | |
| Psh/Pst_1 | Compair | Wheat | 2 | 8 | | | | | | | |
| Psh/Pst_1 | Avocet Yr9 | Wheat | 1 | 7 | | | | | | | |
| Psh/Pst_1 | Avocet Yr9 | Wheat | 2 | | 7 | | | | | | |
| Psh/Pst_1 | Sleipner | Wheat | 1 | 7 | | | | | | | |
| Psh/Pst_1 | Sleipner | Wheat | 2 | | 7 | | | | | | |
| Psh/Pst_1 | Moro | Wheat | 1 | 8 | | | | | | | |
| Psh/Pst_1 | Moro | Wheat | 2 | 8 | | | | | | | |
| Psh/Pst_1 | Avocet Yr15 | Wheat | 1 | 8 | | | | | | | |
| Psh/Pst_1 | Avocet Yr15 | Wheat | 2 | 8 | | | | | | | |
| Psh/Pst_1 | VPM1 | Wheat | 1 | 6 | | | | | | | |
| Psh/Pst_1 | VPM1 | Wheat | 2 | 3 | 3 | | | | | | |
| Psh/Pst_1 | Avocet Yr17 | Wheat | 1 | 7 | | | | | | | |
| Psh/Pst_1 | Avocet Yr17 | Wheat | 2 | 7 | | | | | | | |
| Psh/Pst_1 | Avocet Yr24 | Wheat | 1 | 8 | | | | | | | |
| Psh/Pst_1 | Avocet Yr24 | Wheat | 2 | 8 | | | | | | | |
| Psh/Pst_1 | TP 981 | Wheat | 1 | 7 | | | | | | | |
| Psh/Pst_1 | TP 981 | Wheat | 2 | 7 | | | | | | | |
| Psh/Pst_1 | Avocet Yr27 | Wheat | 1 | 8 | | | | | | | |
| Psh/Pst_1 | Avocet Yr27 | Wheat | 2 | 8 | | | | | | | |
| Psh/Pst_1 | Avocet Yr32 | Wheat | 1 | 7 | | | | | | | |
| Psh/Pst_1 | Avocet Yr32 | Wheat | 2 | | 7 | | | | | | |
| Psh/Pst_1 | Avocet SP | Wheat | 1 | 8 | | | | | | | |
| Psh/Pst_1 | Avocet SP | Wheat | 2 | 8 | | | | | | | |
| Psh/Pst_1 | Avocet S | Wheat | 1 | | 4 | 3 | | | | | |
| Psh/Pst_1 | Avocet S | Wheat | 2 | | | 3 | 2 | 2 | | | |
| Psh/Pst_1 | Triticum monococcum | Wheat | 1 | 7 | | | | | | | |
| Psh/Pst_1 | Triticum monococcum | Wheat | 2 | 7 | | | | | | | |
| Psh/Pst_1 | Triticum dicoccon | Wheat | 1 | 7 | | | | | | | |
| Psh/Pst_1 | Triticum dicoccon | Wheat | 2 | 7 | | | | | | | |
| Psh/Pst_2 | Afzal | Barley | 1 | | | | | | | 1 | 6 |
| Psh/Pst_2 | Afzal | Barley | 2 | | | | | | | | 7 |
| Psh/Pst_2 | Emir | Barley | 1 | | | | | | | 2 | 4 |
| Psh/Pst_2 | Emir | Barley | 2 | | | | | | | 1 | 5 |
| Psh/Pst_2 | Abed Binder | Barley | 1 | 8 | | | | | | | |
| Psh/Pst_2 | Abed Binder | Barley | 2 | 8 | | | | | | | |

| | | | | | | | | | | | |
|---|---|---|---|---|---|---|---|---|---|---|---|
| Psh/Pst_2 | Bancroft | Barley | 1 | 8 | | | | | | | |
| Psh/Pst_2 | Bancroft | Barley | 2 | 8 | | | | | | | |
| Psh/Pst_2 | Heils Franken | Barley | 1 | 8 | | | | | | | |
| Psh/Pst_2 | Heils Franken | Barley | 2 | 8 | | | | | | | |
| Psh/Pst_2 | Trumpf | Barley | 1 | 6 | | | | | | | |
| Psh/Pst_2 | Trumpf | Barley | 2 | 6 | | | | | | | |
| Psh/Pst_2 | I5 | Barley | 1 | 7 | | | | | | | |
| Psh/Pst_2 | I5 | Barley | 2 | 7 | | | | | | | |
| Psh/Pst_2 | Hiproly | Barley | 1 | 8 | | | | | | | |
| Psh/Pst_2 | Hiproly | Barley | 2 | 8 | | | | | | | |
| Psh/Pst_2 | Bigo | Barley | 1 | 6 | | | | | | | |
| Psh/Pst_2 | Bigo | Barley | 2 | 3 | 3 | | | | | | |
| Psh/Pst_2 | Astrix | Barley | 1 | 7 | | | | | | | |
| Psh/Pst_2 | Astrix | Barley | 2 | 4 | 3 | | | | | | |
| Psh/Pst_2 | Varanda | Barley | 1 | | | | | | 3 | 2 | 1 |
| Psh/Pst_2 | Varanda | Barley | 2 | | | | | | | 2 | 4 |
| Psh/Pst_2 | Hordeum spontaneum | Barley | 1 | | | | | | | 3 | 3 |
| Psh/Pst_2 | Hordeum spontaneum | Barley | 2 | | | | | | | 3 | 3 |
| Psh/Pst_2 | Aegilops tauschii | Other | 1 | 8 | | | | | | | |
| Psh/Pst_2 | Aegilops tauschii | Other | 2 | | 8 | | | | | | |
| Psh/Pst_2 | Elymus repens | Other | 1 | 7 | | | | | | | |
| Psh/Pst_2 | Elymus repens | Other | 2 | 7 | | | | | | | |
| Psh/Pst_2 | Morocco | Wheat | 1 | | | 7 | | | | | |
| Psh/Pst_2 | Morocco | Wheat | 2 | | | 3 | 4 | | | | |
| Psh/Pst_2 | Chinese 166 | Wheat | 1 | | | | | | | | 7 |
| Psh/Pst_2 | Chinese 166 | Wheat | 2 | | | | | | | | 7 |
| Psh/Pst_2 | Avocet Yr1 | Wheat | 1 | | 6 | | | | | | |
| Psh/Pst_2 | Avocet Yr1 | Wheat | 2 | | | 6 | | | | | |
| Psh/Pst_2 | Kalyansona | Wheat | 1 | | 2 | 4 | | | | | |
| Psh/Pst_2 | Kalyansona | Wheat | 2 | | 3 | 3 | | | | | |
| Psh/Pst_2 | Vilmorin 23 | Wheat | 1 | 6 | | | | | | | |
| Psh/Pst_2 | Vilmorin 23 | Wheat | 2 | 6 | | | | | | | |
| Psh/Pst_2 | Hybrid 46 | Wheat | 1 | 8 | | | | | | | |
| Psh/Pst_2 | Hybrid 46 | Wheat | 2 | 8 | | | | | | | |
| Psh/Pst_2 | Suwon/Omar | Wheat | 1 | 8 | | | | | | | |
| Psh/Pst_2 | Suwon/Omar | Wheat | 2 | 8 | | | | | | | |
| Psh/Pst_2 | Avocet Yr5 | Wheat | 1 | 7 | | | | | | | |
| Psh/Pst_2 | Avocet Yr5 | Wheat | 2 | 7 | | | | | | | |
| Psh/Pst_2 | Avocet Yr6 | Wheat | 1 | | 7 | | | | | | |
| Psh/Pst_2 | Avocet Yr6 | Wheat | 2 | | | 7 | | | | | |
| Psh/Pst_2 | Lee | Wheat | 1 | | 7 | | | | | | |
| Psh/Pst_2 | Lee | Wheat | 2 | | 7 | | | | | | |
| Psh/Pst_2 | Avocet Yr7 | Wheat | 1 | 6 | | | | | | | |
| Psh/Pst_2 | Avocet Yr7 | Wheat | 2 | 6 | | | | | | | |
| Psh/Pst_2 | Avocet Yr8 | Wheat | 1 | | 7 | | | | | | |
| Psh/Pst_2 | Avocet Yr8 | Wheat | 2 | | | 7 | | | | | |
| Psh/Pst_2 | Compair | Wheat | 1 | 8 | | | | | | | |
| Psh/Pst_2 | Compair | Wheat | 2 | 8 | | | | | | | |
| Psh/Pst_2 | Avocet Yr9 | Wheat | 1 | 4 | 4 | | | | | | |
| Psh/Pst_2 | Avocet Yr9 | Wheat | 2 | | 4 | 4 | | | | | |

| | | | | | | | | | | | |
|---|---|---|---|---|---|---|---|---|---|---|---|
| Psh/Pst_2 | Sleipner | Wheat | 1 | 8 | | | | | | | |
| Psh/Pst_2 | Sleipner | Wheat | 2 | | 8 | | | | | | |
| Psh/Pst_2 | Moro | Wheat | 1 | 8 | | | | | | | |
| Psh/Pst_2 | Moro | Wheat | 2 | 8 | | | | | | | |
| Psh/Pst_2 | Avocet Yr15 | Wheat | 1 | 8 | | | | | | | |
| Psh/Pst_2 | Avocet Yr15 | Wheat | 2 | 8 | | | | | | | |
| Psh/Pst_2 | VPM1 | Wheat | 1 | 7 | | | | | | | |
| Psh/Pst_2 | VPM1 | Wheat | 2 | 7 | | | | | | | |
| Psh/Pst_2 | Avocet Yr17 | Wheat | 1 | 7 | | | | | | | |
| Psh/Pst_2 | Avocet Yr17 | Wheat | 2 | 7 | | | | | | | |
| Psh/Pst_2 | Avocet Yr24 | Wheat | 1 | 8 | | | | | | | |
| Psh/Pst_2 | Avocet Yr24 | Wheat | 2 | 8 | | | | | | | |
| Psh/Pst_2 | TP 981 | Wheat | 1 | 7 | | | | | | | |
| Psh/Pst_2 | TP 981 | Wheat | 2 | 7 | | | | | | | |
| Psh/Pst_2 | Avocet Yr27 | Wheat | 1 | 8 | | | | | | | |
| Psh/Pst_2 | Avocet Yr27 | Wheat | 2 | 8 | | | | | | | |
| Psh/Pst_2 | Avocet Yr32 | Wheat | 1 | | 7 | | | | | | |
| Psh/Pst_2 | Avocet Yr32 | Wheat | 2 | | | 7 | | | | | |
| Psh/Pst_2 | Avocet SP | Wheat | 1 | 8 | | | | | | | |
| Psh/Pst_2 | Avocet SP | Wheat | 2 | 8 | | | | | | | |
| Psh/Pst_2 | Avocet S | Wheat | 1 | | | 8 | | | | | |
| Psh/Pst_2 | Avocet S | Wheat | 2 | | | | 2 | 3 | 3 | | |
| Psh/Pst_2 | Triticum monococcum | Wheat | 1 | 7 | | | | | | | |
| Psh/Pst_2 | Triticum monococcum | Wheat | 2 | 7 | | | | | | | |
| Psh/Pst_2 | Triticum dicoccon | Wheat | 1 | 7 | | | | | | | |
| Psh/Pst_2 | Triticum dicoccon | Wheat | 2 | 7 | | | | | | | |
| Psh/Pst_3 | Afzal | Barley | 1 | | | | | | | 2 | 5 |
| Psh/Pst_3 | Afzal | Barley | 2 | | | | | | | | 7 |
| Psh/Pst_3 | Emir | Barley | 1 | | | | | | | 1 | 6 |
| Psh/Pst_3 | Emir | Barley | 2 | | | | | | | | 7 |
| Psh/Pst_3 | Abed Binder | Barley | 1 | 8 | | | | | | | |
| Psh/Pst_3 | Abed Binder | Barley | 2 | 4 | 4 | | | | | | |
| Psh/Pst_3 | Bancroft | Barley | 1 | 8 | | | | | | | |
| Psh/Pst_3 | Bancroft | Barley | 2 | 4 | 4 | | | | | | |
| Psh/Pst_3 | Heils Franken | Barley | 1 | 7 | | | | | | | |
| Psh/Pst_3 | Heils Franken | Barley | 2 | 4 | 3 | | | | | | |
| Psh/Pst_3 | Trumpf | Barley | 1 | 6 | | | | | | | |
| Psh/Pst_3 | Trumpf | Barley | 2 | 6 | | | | | | | |
| Psh/Pst_3 | I5 | Barley | 1 | 7 | | | | | | | |
| Psh/Pst_3 | I5 | Barley | 2 | 7 | | | | | | | |
| Psh/Pst_3 | Hiproly | Barley | 1 | 8 | | | | | | | |
| Psh/Pst_3 | Hiproly | Barley | 2 | 8 | | | | | | | |
| Psh/Pst_3 | Bigo | Barley | 1 | 7 | | | | | | | |
| Psh/Pst_3 | Bigo | Barley | 2 | 4 | 3 | | | | | | |
| Psh/Pst_3 | Astrix | Barley | 1 | 8 | | | | | | | |
| Psh/Pst_3 | Astrix | Barley | 2 | 8 | | | | | | | |
| Psh/Pst_3 | Varanda | Barley | 1 | | | | | | | 4 | 4 |
| Psh/Pst_3 | Varanda | Barley | 2 | | | | | | | 4 | 4 |
| Psh/Pst_3 | Hordeum spontaneum | Barley | 1 | | | | | | | 4 | 2 |
| Psh/Pst_3 | Hordeum spontaneum | Barley | 2 | | | | | | | 1 | 5 |

| | | | | | | | | | | | |
|---|---|---|---|---|---|---|---|---|---|---|---|
| Psh/Pst_3 | Aegilops tauschii | Other | 1 | 7 | | | | | | | |
| Psh/Pst_3 | Aegilops tauschii | Other | 2 | | 7 | | | | | | |
| Psh/Pst_3 | Elymus repens | Other | 1 | 7 | | | | | | | |
| Psh/Pst_3 | Elymus repens | Other | 2 | 7 | | | | | | | |
| Psh/Pst_3 | Morocco | Wheat | 1 | | 3 | 4 | | | | | |
| Psh/Pst_3 | Morocco | Wheat | 2 | | | 4 | 3 | | | | |
| Psh/Pst_3 | Chinese 166 | Wheat | 1 | | | | | | | | 8 |
| Psh/Pst_3 | Chinese 166 | Wheat | 2 | | | | | | | | 8 |
| Psh/Pst_3 | Avocet Yr1 | Wheat | 1 | | 6 | | | | | | |
| Psh/Pst_3 | Avocet Yr1 | Wheat | 2 | | | 6 | | | | | |
| Psh/Pst_3 | Kalyansona | Wheat | 1 | | 6 | | | | | | |
| Psh/Pst_3 | Kalyansona | Wheat | 2 | | 3 | 3 | | | | | |
| Psh/Pst_3 | Vilmorin 23 | Wheat | 1 | 8 | | | | | | | |
| Psh/Pst_3 | Vilmorin 23 | Wheat | 2 | 4 | 4 | | | | | | |
| Psh/Pst_3 | Hybrid 46 | Wheat | 1 | 8 | | | | | | | |
| Psh/Pst_3 | Hybrid 46 | Wheat | 2 | 8 | | | | | | | |
| Psh/Pst_3 | Suwon/Omar | Wheat | 1 | 7 | | | | | | | |
| Psh/Pst_3 | Suwon/Omar | Wheat | 2 | 7 | | | | | | | |
| Psh/Pst_3 | Avocet Yr5 | Wheat | 1 | 7 | | | | | | | |
| Psh/Pst_3 | Avocet Yr5 | Wheat | 2 | 7 | | | | | | | |
| Psh/Pst_3 | Avocet Yr6 | Wheat | 1 | | 8 | | | | | | |
| Psh/Pst_3 | Avocet Yr6 | Wheat | 2 | | | 8 | | | | | |
| Psh/Pst_3 | Lee | Wheat | 1 | | 4 | 4 | | | | | |
| Psh/Pst_3 | Lee | Wheat | 2 | | 4 | 4 | | | | | |
| Psh/Pst_3 | Avocet Yr7 | Wheat | 1 | | 6 | | | | | | |
| Psh/Pst_3 | Avocet Yr7 | Wheat | 2 | | 6 | | | | | | |
| Psh/Pst_3 | Avocet Yr8 | Wheat | 1 | | 4 | 4 | | | | | |
| Psh/Pst_3 | Avocet Yr8 | Wheat | 2 | | | 8 | | | | | |
| Psh/Pst_3 | Compair | Wheat | 1 | 8 | | | | | | | |
| Psh/Pst_3 | Compair | Wheat | 2 | 4 | 4 | | | | | | |
| Psh/Pst_3 | Avocet Yr9 | Wheat | 1 | | 8 | | | | | | |
| Psh/Pst_3 | Avocet Yr9 | Wheat | 2 | | 4 | 4 | | | | | |
| Psh/Pst_3 | Sleipner | Wheat | 1 | 8 | | | | | | | |
| Psh/Pst_3 | Sleipner | Wheat | 2 | 8 | | | | | | | |
| Psh/Pst_3 | Moro | Wheat | 1 | 7 | | | | | | | |
| Psh/Pst_3 | Moro | Wheat | 2 | 7 | | | | | | | |
| Psh/Pst_3 | Avocet Yr15 | Wheat | 1 | 8 | | | | | | | |
| Psh/Pst_3 | Avocet Yr15 | Wheat | 2 | 8 | | | | | | | |
| Psh/Pst_3 | VPM1 | Wheat | 1 | 7 | | | | | | | |
| Psh/Pst_3 | VPM1 | Wheat | 2 | 7 | | | | | | | |
| Psh/Pst_3 | Avocet Yr17 | Wheat | 1 | 6 | | | | | | | |
| Psh/Pst_3 | Avocet Yr17 | Wheat | 2 | 6 | | | | | | | |
| Psh/Pst_3 | Avocet Yr24 | Wheat | 1 | 7 | | | | | | | |
| Psh/Pst_3 | Avocet Yr24 | Wheat | 2 | 7 | | | | | | | |
| Psh/Pst_3 | TP 981 | Wheat | 1 | 6 | | | | | | | |
| Psh/Pst_3 | TP 981 | Wheat | 2 | 6 | | | | | | | |
| Psh/Pst_3 | Avocet Yr27 | Wheat | 1 | 8 | | | | | | | |
| Psh/Pst_3 | Avocet Yr27 | Wheat | 2 | 8 | | | | | | | |
| Psh/Pst_3 | Avocet Yr32 | Wheat | 1 | 8 | | | | | | | |
| Psh/Pst_3 | Avocet Yr32 | Wheat | 2 | | 8 | | | | | | |

| | | | | | | | | | | | |
|---|---|---|---|---|---|---|---|---|---|---|---|
| Psh/Pst_3 | Avocet SP | Wheat | 1 | 8 | | | | | | | |
| Psh/Pst_3 | Avocet SP | Wheat | 2 | 8 | | | | | | | |
| Psh/Pst_3 | Avocet S | Wheat | 1 | | | 7 | | | | | |
| Psh/Pst_3 | Avocet S | Wheat | 2 | | | 2 | 3 | 2 | | | |
| Psh/Pst_3 | Triticum monococcum | Wheat | 1 | 6 | | | | | | | |
| Psh/Pst_3 | Triticum monococcum | Wheat | 2 | 6 | | | | | | | |
| Psh/Pst_3 | Triticum dicoccon | Wheat | 1 | 7 | | | | | | | |
| Psh/Pst_3 | Triticum dicoccon | Wheat | 2 | 7 | | | | | | | |
| Psh/Pst_4 | Abed Binder | Barley | 1 | 7 | | | | | | | |
| Psh/Pst_4 | Abed Binder | Barley | 2 | 7 | | | | | | | |
| Psh/Pst_4 | Afzal | Barley | 1 | | | | | | | 2 | 4 |
| Psh/Pst_4 | Afzal | Barley | 2 | | | | | | | | 6 |
| Psh/Pst_4 | Astrix | Barley | 1 | 7 | | | | | | | |
| Psh/Pst_4 | Astrix | Barley | 2 | 7 | | | | | | | |
| Psh/Pst_4 | Bancroft | Barley | 1 | 8 | | | | | | | |
| Psh/Pst_4 | Bancroft | Barley | 2 | 4 | 4 | | | | | | |
| Psh/Pst_4 | Bigo | Barley | 1 | 7 | | | | | | | |
| Psh/Pst_4 | Bigo | Barley | 2 | 3 | 4 | | | | | | |
| Psh/Pst_4 | Emir | Barley | 1 | | | | | | | 1 | 5 |
| Psh/Pst_4 | Emir | Barley | 2 | | | | | | | | 6 |
| Psh/Pst_4 | Heils Franken | Barley | 1 | 7 | | | | | | | |
| Psh/Pst_4 | Heils Franken | Barley | 2 | 7 | | | | | | | |
| Psh/Pst_4 | Hiproly | Barley | 1 | 4 | 4 | | | | | | |
| Psh/Pst_4 | Hiproly | Barley | 2 | 4 | 4 | | | | | | |
| Psh/Pst_4 | Hordeum spontaneum | Barley | 1 | | | | | | 3 | 3 | |
| Psh/Pst_4 | Hordeum spontaneum | Barley | 2 | | | | | | 3 | 3 | |
| Psh/Pst_4 | I5 | Barley | 1 | 4 | 4 | | | | | | |
| Psh/Pst_4 | I5 | Barley | 2 | 4 | 4 | | | | | | |
| Psh/Pst_4 | Trumpf | Barley | 1 | 4 | 4 | | | | | | |
| Psh/Pst_4 | Trumpf | Barley | 2 | 4 | 4 | | | | | | |
| Psh/Pst_4 | Varanda | Barley | 1 | | | | | | | 4 | 2 |
| Psh/Pst_4 | Varanda | Barley | 2 | | | | | | | 4 | 2 |
| Psh/Pst_4 | Aegilops tauschii | Other | 1 | 8 | | | | | | | |
| Psh/Pst_4 | Aegilops tauschii | Other | 2 | 4 | 4 | | | | | | |
| Psh/Pst_4 | Elymus repens | Other | 1 | 7 | | | | | | | |
| Psh/Pst_4 | Elymus repens | Other | 2 | 7 | | | | | | | |
| Psh/Pst_4 | Avocet S | Wheat | 1 | | | 4 | 2 | | | | |
| Psh/Pst_4 | Avocet S | Wheat | 2 | | | | | 2 | 4 | | |
| Psh/Pst_4 | Avocet SP | Wheat | 1 | 8 | | | | | | | |
| Psh/Pst_4 | Avocet SP | Wheat | 2 | 8 | | | | | | | |
| Psh/Pst_4 | Avocet Yr1 | Wheat | 1 | 6 | | | | | | | |
| Psh/Pst_4 | Avocet Yr1 | Wheat | 2 | 6 | | | | | | | |
| Psh/Pst_4 | Avocet Yr15 | Wheat | 1 | 8 | | | | | | | |
| Psh/Pst_4 | Avocet Yr15 | Wheat | 2 | 8 | | | | | | | |
| Psh/Pst_4 | Avocet Yr17 | Wheat | 1 | 7 | | | | | | | |
| Psh/Pst_4 | Avocet Yr17 | Wheat | 2 | 7 | | | | | | | |
| Psh/Pst_4 | Avocet Yr24 | Wheat | 1 | 8 | | | | | | | |
| Psh/Pst_4 | Avocet Yr24 | Wheat | 2 | 8 | | | | | | | |
| Psh/Pst_4 | Avocet Yr27 | Wheat | 1 | 8 | | | | | | | |
| Psh/Pst_4 | Avocet Yr27 | Wheat | 2 | 8 | | | | | | | |

| | | | | | | | | | | | |
|---|---|---|---|---|---|---|---|---|---|---|---|
| Psh/Pst_4 | Avocet Yr32 | Wheat | 1 | | 7 | | | | | | |
| Psh/Pst_4 | Avocet Yr32 | Wheat | 2 | | 7 | | | | | | |
| Psh/Pst_4 | Avocet Yr5 | Wheat | 1 | 7 | | | | | | | |
| Psh/Pst_4 | Avocet Yr5 | Wheat | 2 | 7 | | | | | | | |
| Psh/Pst_4 | Avocet Yr6 | Wheat | 1 | | 6 | | | | | | |
| Psh/Pst_4 | Avocet Yr6 | Wheat | 2 | | | 3 | 3 | | | | |
| Psh/Pst_4 | Avocet Yr7 | Wheat | 1 | 6 | | | | | | | |
| Psh/Pst_4 | Avocet Yr7 | Wheat | 2 | 6 | | | | | | | |
| Psh/Pst_4 | Avocet Yr8 | Wheat | 1 | 7 | | | | | | | |
| Psh/Pst_4 | Avocet Yr8 | Wheat | 2 | 7 | | | | | | | |
| Psh/Pst_4 | Avocet Yr9 | Wheat | 1 | | | 3 | 3 | | | | |
| Psh/Pst_4 | Avocet Yr9 | Wheat | 2 | | | 3 | 2 | 1 | | | |
| Psh/Pst_4 | Chinese 166 | Wheat | 1 | 6 | | | | | | | |
| Psh/Pst_4 | Chinese 166 | Wheat | 2 | 6 | | | | | | | |
| Psh/Pst_4 | Compair | Wheat | 1 | 4 | 4 | | | | | | |
| Psh/Pst_4 | Compair | Wheat | 2 | 8 | | | | | | | |
| Psh/Pst_4 | Hybrid 46 | Wheat | 1 | 7 | | | | | | | |
| Psh/Pst_4 | Hybrid 46 | Wheat | 2 | 7 | | | | | | | |
| Psh/Pst_4 | Kalyansona | Wheat | 1 | | | | | 3 | 2 | 1 | |
| Psh/Pst_4 | Kalyansona | Wheat | 2 | | | | | | 2 | 4 | |
| Psh/Pst_4 | Lee | Wheat | 1 | | | 7 | | | | | |
| Psh/Pst_4 | Lee | Wheat | 2 | | | 7 | | | | | |
| Psh/Pst_4 | Moro | Wheat | 1 | 7 | | | | | | | |
| Psh/Pst_4 | Moro | Wheat | 2 | 7 | | | | | | | |
| Psh/Pst_4 | Morocco | Wheat | 1 | | | | | | | 3 | 4 |
| Psh/Pst_4 | Morocco | Wheat | 2 | | | | | | | | 7 |
| Psh/Pst_4 | Sleipner | Wheat | 1 | 7 | | | | | | | |
| Psh/Pst_4 | Sleipner | Wheat | 2 | | 7 | | | | | | |
| Psh/Pst_4 | Suwon/Omar | Wheat | 1 | 8 | | | | | | | |
| Psh/Pst_4 | Suwon/Omar | Wheat | 2 | 8 | | | | | | | |
| Psh/Pst_4 | TP 981 | Wheat | 1 | 7 | | | | | | | |
| Psh/Pst_4 | TP 981 | Wheat | 2 | 7 | | | | | | | |
| Psh/Pst_4 | Triticum dicoccon | Wheat | 1 | 7 | | | | | | | |
| Psh/Pst_4 | Triticum dicoccon | Wheat | 2 | 7 | | | | | | | |
| Psh/Pst_4 | Triticum monococcum | Wheat | 1 | 7 | | | | | | | |
| Psh/Pst_4 | Triticum monococcum | Wheat | 2 | 7 | | | | | | | |
| Psh/Pst_4 | Vilmorin 23 | Wheat | 1 | 6 | | | | | | | |
| Psh/Pst_4 | Vilmorin 23 | Wheat | 2 | 6 | | | | | | | |
| Psh/Pst_4 | VPM1 | Wheat | 1 | 7 | | | | | | | |
| Psh/Pst_4 | VPM1 | Wheat | 2 | 7 | | | | | | | |
| Psh | Afzal | Barley | 1 | | | | | | | | 7 |
| Psh | Afzal | Barley | 2 | | | | | | | | 7 |
| Psh | Emir | Barley | 1 | | | | | | | 1 | 6 |
| Psh | Emir | Barley | 2 | | | | | | | 1 | 6 |
| Psh | Abed Binder | Barley | 1 | 5 | 3 | | | | | | |
| Psh | Abed Binder | Barley | 2 | 8 | | | | | | | |
| Psh | Bancroft | Barley | 1 | 8 | | | | | | | |
| Psh | Bancroft | Barley | 2 | 8 | | | | | | | |
| Psh | Heils Franken | Barley | 1 | 4 | 4 | | | | | | |
| Psh | Heils Franken | Barley | 2 | 8 | | | | | | | |

| | | | | | | | | | | | |
|---|---|---|---|---|---|---|---|---|---|---|---|
| Psh | Trumpf | Barley | 1 | 8 | | | | | | | |
| Psh | Trumpf | Barley | 2 | 8 | | | | | | | |
| Psh | I5 | Barley | 1 | 5 | 2 | | | | | | |
| Psh | I5 | Barley | 2 | 7 | | | | | | | |
| Psh | Hiproly | Barley | 1 | 7 | | | | | | | |
| Psh | Hiproly | Barley | 2 | 7 | | | | | | | |
| Psh | Bigo | Barley | 1 | 7 | | | | | | | |
| Psh | Bigo | Barley | 2 | 7 | | | | | | | |
| Psh | Astrix | Barley | 1 | 6 | | | | | | | |
| Psh | Astrix | Barley | 2 | 6 | | | | | | | |
| Psh | Varanda | Barley | 1 | | | | | | | 1 | 5 |
| Psh | Varanda | Barley | 2 | | | | | | | | 6 |
| Psh | Hordeum spontaneum | Barley | 1 | | | | | | | 2 | 4 |
| Psh | Hordeum spontaneum | Barley | 2 | | | | | | | | 6 |
| Psh | Aegilops tauschii | Other | 1 | 8 | | | | | | | |
| Psh | Aegilops tauschii | Other | 2 | 8 | | | | | | | |
| Psh | Elymus repens | Other | 1 | 6 | | | | | | | |
| Psh | Elymus repens | Other | 2 | 6 | | | | | | | |
| Psh | Morocco | Wheat | 1 | 8 | | | | | | | |
| Psh | Morocco | Wheat | 2 | 8 | | | | | | | |
| Psh | Chinese 166 | Wheat | 1 | 8 | | | | | | | |
| Psh | Chinese 166 | Wheat | 2 | 8 | | | | | | | |
| Psh | Avocet Yr1 | Wheat | 1 | 7 | | | | | | | |
| Psh | Avocet Yr1 | Wheat | 2 | 7 | | | | | | | |
| Psh | Kalyansona | Wheat | 1 | 6 | | | | | | | |
| Psh | Kalyansona | Wheat | 2 | 6 | | | | | | | |
| Psh | Vilmorin 23 | Wheat | 1 | 8 | | | | | | | |
| Psh | Vilmorin 23 | Wheat | 2 | 8 | | | | | | | |
| Psh | Hybrid 46 | Wheat | 1 | 8 | | | | | | | |
| Psh | Hybrid 46 | Wheat | 2 | 8 | | | | | | | |
| Psh | Suwon/Omar | Wheat | 1 | 8 | | | | | | | |
| Psh | Suwon/Omar | Wheat | 2 | 8 | | | | | | | |
| Psh | Avocet Yr5 | Wheat | 1 | 7 | | | | | | | |
| Psh | Avocet Yr5 | Wheat | 2 | 7 | | | | | | | |
| Psh | Avocet Yr6 | Wheat | 1 | 8 | | | | | | | |
| Psh | Avocet Yr6 | Wheat | 2 | 8 | | | | | | | |
| Psh | Lee | Wheat | 1 | 8 | | | | | | | |
| Psh | Lee | Wheat | 2 | 8 | | | | | | | |
| Psh | Avocet Yr7 | Wheat | 1 | 7 | | | | | | | |
| Psh | Avocet Yr7 | Wheat | 2 | 7 | | | | | | | |
| Psh | Avocet Yr8 | Wheat | 1 | 7 | | | | | | | |
| Psh | Avocet Yr8 | Wheat | 2 | 7 | | | | | | | |
| Psh | Compair | Wheat | 1 | 8 | | | | | | | |
| Psh | Compair | Wheat | 2 | 8 | | | | | | | |
| Psh | Avocet Yr9 | Wheat | 1 | 8 | | | | | | | |
| Psh | Avocet Yr9 | Wheat | 2 | 8 | | | | | | | |
| Psh | Sleipner | Wheat | 1 | 7 | | | | | | | |
| Psh | Sleipner | Wheat | 2 | 7 | | | | | | | |
| Psh | Moro | Wheat | 1 | 8 | | | | | | | |
| Psh | Moro | Wheat | 2 | 8 | | | | | | | |

| | | | | | | |
|---|---|---|---|---|---|---|
| Psh | Avocet Yr15 | Wheat | 1 | 6 | | |
| Psh | Avocet Yr15 | Wheat | 2 | 6 | | |
| Psh | VPM1 | Wheat | 1 | 7 | | |
| Psh | VPM1 | Wheat | 2 | 7 | | |
| Psh | Avocet Yr17 | Wheat | 1 | 6 | | |
| Psh | Avocet Yr17 | Wheat | 2 | 6 | | |
| Psh | Avocet Yr24 | Wheat | 1 | 8 | | |
| Psh | Avocet Yr24 | Wheat | 2 | 8 | | |
| Psh | TP 981 | Wheat | 1 | 7 | | |
| Psh | TP 981 | Wheat | 2 | 7 | | |
| Psh | Avocet Yr27 | Wheat | 1 | 7 | | |
| Psh | Avocet Yr27 | Wheat | 2 | 7 | | |
| Psh | Avocet Yr32 | Wheat | 1 | 8 | | |
| Psh | Avocet Yr32 | Wheat | 2 | 8 | | |
| Psh | Avocet SP | Wheat | 1 | 6 | | |
| Psh | Avocet SP | Wheat | 2 | 6 | | |
| Psh | Avocet S | Wheat | 1 | 8 | | |
| Psh | Avocet S | Wheat | 2 | 8 | | |
| Psh | Triticum monococcum | Wheat | 1 | 8 | | |
| Psh | Triticum monococcum | Wheat | 2 | 8 | | |
| Psh | Triticum dicoccon | Wheat | 1 | 8 | | |
| Psh | Triticum dicoccon | Wheat | 2 | 8 | | |
| Pst | Afzal | Barley | 1 | 6 | | |
| Pst | Afzal | Barley | 2 | 6 | | |
| Pst | Emir | Barley | 1 | 8 | | |
| Pst | Emir | Barley | 2 | 8 | | |
| Pst | Abed Binder | Barley | 1 | 8 | | |
| Pst | Abed Binder | Barley | 2 | 8 | | |
| Pst | Bancroft | Barley | 1 | 7 | | |
| Pst | Bancroft | Barley | 2 | 7 | | |
| Pst | Heils Franken | Barley | 1 | 7 | | |
| Pst | Heils Franken | Barley | 2 | 7 | | |
| Pst | Trumpf | Barley | 1 | 7 | | |
| Pst | Trumpf | Barley | 2 | 7 | | |
| Pst | I5 | Barley | 1 | 8 | | |
| Pst | I5 | Barley | 2 | 8 | | |
| Pst | Hiproly | Barley | 1 | 8 | | |
| Pst | Hiproly | Barley | 2 | 8 | | |
| Pst | Bigo | Barley | 1 | 6 | | |
| Pst | Bigo | Barley | 2 | 6 | | |
| Pst | Astrix | Barley | 1 | 6 | | |
| Pst | Astrix | Barley | 2 | 6 | | |
| Pst | Varanda | Barley | 1 | 8 | | |
| Pst | Varanda | Barley | 2 | 8 | | |
| Pst | Hordeum spontaneum | Barley | 1 | 7 | | |
| Pst | Hordeum spontaneum | Barley | 2 | 7 | | |
| Pst | Aegilops tauschii | Other | 1 | | 3 | 4 |
| Pst | Aegilops tauschii | Other | 2 | | 2 | 5 |
| Pst | Elymus repens | Other | 1 | 6 | | |
| Pst | Elymus repens | Other | 2 | 6 | | |

| | | | | | | | | | | | | |
|---|---|---|---|---|---|---|---|---|---|---|---|---|
| Pst | Morocco | Wheat | 1 | | | | | | | | 8 | |
| Pst | Morocco | Wheat | 2 | | | | | | | | 8 | |
| Pst | Chinese 166 | Wheat | 1 | | | | | | | | 8 | |
| Pst | Chinese 166 | Wheat | 2 | | | | | | | | 4 | 4 |
| Pst | Avocet Yr1 | Wheat | 1 | | | | | | | | 6 | |
| Pst | Avocet Yr1 | Wheat | 2 | | | | | | | | 6 | |
| Pst | Kalyansona | Wheat | 1 | | | | | | | 2 | 5 | |
| Pst | Kalyansona | Wheat | 2 | | | | | | | | 7 | |
| Pst | Vilmorin 23 | Wheat | 1 | | | | | | | | 7 | |
| Pst | Vilmorin 23 | Wheat | 2 | | | | | | | | 7 | |
| Pst | Hybrid 46 | Wheat | 1 | 7 | | | | | | | | |
| Pst | Hybrid 46 | Wheat | 2 | 7 | | | | | | | | |
| Pst | Suwon/Omar | Wheat | 1 | 7 | | | | | | | | |
| Pst | Suwon/Omar | Wheat | 2 | | 7 | | | | | | | |
| Pst | Avocet Yr5 | Wheat | 1 | 6 | | | | | | | | |
| Pst | Avocet Yr5 | Wheat | 2 | 6 | | | | | | | | |
| Pst | Avocet Yr6 | Wheat | 1 | | | | | | | | 7 | |
| Pst | Avocet Yr6 | Wheat | 2 | | | | | | | | 7 | |
| Pst | Lee | Wheat | 1 | | | | | | | | 6 | |
| Pst | Lee | Wheat | 2 | | | | | | | | 6 | |
| Pst | Avocet Yr7 | Wheat | 1 | | | | | | | | 6 | |
| Pst | Avocet Yr7 | Wheat | 2 | | | | | | | | 6 | |
| Pst | Avocet Yr8 | Wheat | 1 | | | | | | | | 7 | |
| Pst | Avocet Yr8 | Wheat | 2 | | | | | | | | 7 | |
| Pst | Compair | Wheat | 1 | | | 4 | 3 | | | | | |
| Pst | Compair | Wheat | 2 | | | 3 | 4 | | | | | |
| Pst | Avocet Yr9 | Wheat | 1 | | | | | | | | 8 | |
| Pst | Avocet Yr9 | Wheat | 2 | | | | | | | | 8 | |
| Pst | Sleipner | Wheat | 1 | | | | | | | | 8 | |
| Pst | Sleipner | Wheat | 2 | | | | | | | | 8 | |
| Pst | Moro | Wheat | 1 | 8 | | | | | | | | |
| Pst | Moro | Wheat | 2 | 8 | | | | | | | | |
| Pst | Avocet Yr15 | Wheat | 1 | 6 | | | | | | | | |
| Pst | Avocet Yr15 | Wheat | 2 | 6 | | | | | | | | |
| Pst | VPM1 | Wheat | 1 | | | | | | | 2 | 4 | |
| Pst | VPM1 | Wheat | 2 | | | | | | | | 6 | |
| Pst | Avocet Yr17 | Wheat | 1 | | | | | | | | 6 | |
| Pst | Avocet Yr17 | Wheat | 2 | | | | | | | | 6 | |
| Pst | Avocet Yr24 | Wheat | 1 | 7 | | | | | | | | |
| Pst | Avocet Yr24 | Wheat | 2 | 7 | | | | | | | | |
| Pst | TP 981 | Wheat | 1 | | | | | | | | 6 | |
| Pst | TP 981 | Wheat | 2 | | | | | | | | 6 | |
| Pst | Avocet Yr27 | Wheat | 1 | | | | 3 | 3 | | | | |
| Pst | Avocet Yr27 | Wheat | 2 | | | | | | 3 | 3 | | |
| Pst | Avocet Yr32 | Wheat | 1 | | | | | | | | 8 | |
| Pst | Avocet Yr32 | Wheat | 2 | | | | | | | | 8 | |
| Pst | Avocet SP | Wheat | 1 | 7 | | | | | | | | |
| Pst | Avocet SP | Wheat | 2 | 7 | | | | | | | | |
| Pst | Avocet S | Wheat | 1 | | | | | | | | 8 | |
| Pst | Avocet S | Wheat | 2 | | | | | | | | 8 | |

| | | | | | | | | | | | |
|---|---|---|---|---|---|---|---|---|---|---|---|
| Pst | Triticum monococcum | Wheat | 1 | | | | | | | 2 | 4 |
| Pst | Triticum monococcum | Wheat | 2 | | | | | | | | 6 |
| Pst | Triticum dicoccon | Wheat | 1 | 7 | | | | | | | |
| Pst | Triticum dicoccon | Wheat | 2 | 7 | | | | | | | |
| Pst/Psh_8 | Abed Binder | Barley | 1 | 4 | 4 | | | | | | |
| Pst/Psh_8 | Abed Binder | Barley | 2 | 4 | 4 | | | | | | |
| Pst/Psh_8 | Afzal | Barley | 1 | | | | | | | 2 | 4 |
| Pst/Psh_8 | Afzal | Barley | 2 | | | | | | | | 6 |
| Pst/Psh_8 | Astrix | Barley | 1 | 7 | | | | | | | |
| Pst/Psh_8 | Astrix | Barley | 2 | 7 | | | | | | | |
| Pst/Psh_8 | Bancroft | Barley | 1 | 7 | | | | | | | |
| Pst/Psh_8 | Bancroft | Barley | 2 | 7 | | | | | | | |
| Pst/Psh_8 | Bigo | Barley | 1 | 8 | | | | | | | |
| Pst/Psh_8 | Bigo | Barley | 2 | 8 | | | | | | | |
| Pst/Psh_8 | Emir | Barley | 1 | | | | | | | 2 | 4 |
| Pst/Psh_8 | Emir | Barley | 2 | | | | | | | 1 | 5 |
| Pst/Psh_8 | Heils Franken | Barley | 1 | 7 | | | | | | | |
| Pst/Psh_8 | Heils Franken | Barley | 2 | 7 | | | | | | | |
| Pst/Psh_8 | Hiproly | Barley | 1 | 8 | | | | | | | |
| Pst/Psh_8 | Hiproly | Barley | 2 | 8 | | | | | | | |
| Pst/Psh_8 | Hordeum spontaneum | Barley | 1 | | | | | 3 | 3 | 1 | |
| Pst/Psh_8 | Hordeum spontaneum | Barley | 2 | | | | | 3 | 3 | 1 | |
| Pst/Psh_8 | I5 | Barley | 1 | 8 | | | | | | | |
| Pst/Psh_8 | I5 | Barley | 2 | 8 | | | | | | | |
| Pst/Psh_8 | Trumpf | Barley | 1 | 7 | | | | | | | |
| Pst/Psh_8 | Trumpf | Barley | 2 | 7 | | | | | | | |
| Pst/Psh_8 | Varanda | Barley | 1 | | | | | 2 | 2 | 2 | |
| Pst/Psh_8 | Varanda | Barley | 2 | | | | | | 4 | 2 | |
| Pst/Psh_8 | Aegilops tauschii | Other | 1 | 8 | | | | | | | |
| Pst/Psh_8 | Aegilops tauschii | Other | 2 | 8 | | | | | | | |
| Pst/Psh_8 | Elymus repens | Other | 1 | 7 | | | | | | | |
| Pst/Psh_8 | Elymus repens | Other | 2 | 7 | | | | | | | |
| Pst/Psh_8 | Avocet S | Wheat | 1 | | | 3 | 3 | | | | |
| Pst/Psh_8 | Avocet S | Wheat | 2 | | | | | 3 | 3 | | |
| Pst/Psh_8 | Avocet SP | Wheat | 1 | 8 | | | | | | | |
| Pst/Psh_8 | Avocet SP | Wheat | 2 | 8 | | | | | | | |
| Pst/Psh_8 | Avocet Yr1 | Wheat | 1 | | | 3 | 3 | | | | |
| Pst/Psh_8 | Avocet Yr1 | Wheat | 2 | | | 2 | 2 | 2 | | | |
| Pst/Psh_8 | Avocet Yr15 | Wheat | 1 | 7 | | | | | | | |
| Pst/Psh_8 | Avocet Yr15 | Wheat | 2 | 7 | | | | | | | |
| Pst/Psh_8 | Avocet Yr17 | Wheat | 1 | 6 | | | | | | | |
| Pst/Psh_8 | Avocet Yr17 | Wheat | 2 | 6 | | | | | | | |
| Pst/Psh_8 | Avocet Yr24 | Wheat | 1 | 8 | | | | | | | |
| Pst/Psh_8 | Avocet Yr24 | Wheat | 2 | 8 | | | | | | | |
| Pst/Psh_8 | Avocet Yr27 | Wheat | 1 | 7 | | | | | | | |
| Pst/Psh_8 | Avocet Yr27 | Wheat | 2 | 7 | | | | | | | |
| Pst/Psh_8 | Avocet Yr32 | Wheat | 1 | | 4 | 4 | | | | | |
| Pst/Psh_8 | Avocet Yr32 | Wheat | 2 | | | 2 | 2 | 3 | | | |
| Pst/Psh_8 | Avocet Yr5 | Wheat | 1 | 7 | | | | | | | |
| Pst/Psh_8 | Avocet Yr5 | Wheat | 2 | 7 | | | | | | | |

| | | | | | | | | | | | |
|---|---|---|---|---|---|---|---|---|---|---|---|
| Pst/Psh_8 | Avocet Yr6 | Wheat | 1 | | | 3 | 4 | | | | |
| Pst/Psh_8 | Avocet Yr6 | Wheat | 2 | | | | 3 | 2 | 2 | | |
| Pst/Psh_8 | Avocet Yr7 | Wheat | 1 | | 6 | | | | | | |
| Pst/Psh_8 | Avocet Yr7 | Wheat | 2 | | 6 | | | | | | |
| Pst/Psh_8 | Avocet Yr8 | Wheat | 1 | | | 7 | | | | | |
| Pst/Psh_8 | Avocet Yr8 | Wheat | 2 | | | 4 | 3 | | | | |
| Pst/Psh_8 | Avocet Yr9 | Wheat | 1 | | | | 2 | 2 | 2 | | |
| Pst/Psh_8 | Avocet Yr9 | Wheat | 2 | | | | | | 3 | 3 | |
| Pst/Psh_8 | Chinese 166 | Wheat | 1 | | | | | | | 3 | 4 |
| Pst/Psh_8 | Chinese 166 | Wheat | 2 | | | | | | | | 7 |
| Pst/Psh_8 | Compair | Wheat | 1 | 7 | | | | | | | |
| Pst/Psh_8 | Compair | Wheat | 2 | | 7 | | | | | | |
| Pst/Psh_8 | Hybrid 46 | Wheat | 1 | 7 | | | | | | | |
| Pst/Psh_8 | Hybrid 46 | Wheat | 2 | 7 | | | | | | | |
| Pst/Psh_8 | Kalyansona | Wheat | 1 | | | 6 | | | | | |
| Pst/Psh_8 | Kalyansona | Wheat | 2 | | | 3 | 3 | | | | |
| Pst/Psh_8 | Lee | Wheat | 1 | | 3 | 4 | | | | | |
| Pst/Psh_8 | Lee | Wheat | 2 | | | 7 | | | | | |
| Pst/Psh_8 | Moro | Wheat | 1 | 7 | | | | | | | |
| Pst/Psh_8 | Moro | Wheat | 2 | 7 | | | | | | | |
| Pst/Psh_8 | Morocco | Wheat | 1 | | | | | | 4 | 4 | |
| Pst/Psh_8 | Morocco | Wheat | 2 | | | | | | | 2 | 6 |
| Pst/Psh_8 | Sleipner | Wheat | 1 | 3 | 4 | | | | | | |
| Pst/Psh_8 | Sleipner | Wheat | 2 | | 7 | | | | | | |
| Pst/Psh_8 | Suwon/Omar | Wheat | 1 | 7 | | | | | | | |
| Pst/Psh_8 | Suwon/Omar | Wheat | 2 | 7 | | | | | | | |
| Pst/Psh_8 | TP 981 | Wheat | 1 | 7 | | | | | | | |
| Pst/Psh_8 | TP 981 | Wheat | 2 | 7 | | | | | | | |
| Pst/Psh_8 | Triticum dicoccon | Wheat | 1 | 7 | | | | | | | |
| Pst/Psh_8 | Triticum dicoccon | Wheat | 2 | 7 | | | | | | | |
| Pst/Psh_8 | Triticum monococcum | Wheat | 1 | 8 | | | | | | | |
| Pst/Psh_8 | Triticum monococcum | Wheat | 2 | 8 | | | | | | | |
| Pst/Psh_8 | Vilmorin 23 | Wheat | 1 | 7 | | | | | | | |
| Pst/Psh_8 | Vilmorin 23 | Wheat | 2 | 7 | | | | | | | |
| Pst/Psh_8 | VPM1 | Wheat | 1 | 7 | | | | | | | |
| Pst/Psh_8 | VPM1 | Wheat | 2 | 7 | | | | | | | |
| Pst/Psh_5 | Abed Binder | Barley | 1 | 6 | | | | | | | |
| Pst/Psh_5 | Abed Binder | Barley | 2 | 6 | | | | | | | |
| Pst/Psh_5 | Afzal | Barley | 1 | | | | | | | 1 | 5 |
| Pst/Psh_5 | Afzal | Barley | 2 | | | | | | | | 6 |
| Pst/Psh_5 | Astrix | Barley | 1 | 4 | 4 | | | | | | |
| Pst/Psh_5 | Astrix | Barley | 2 | 4 | 4 | | | | | | |
| Pst/Psh_5 | Bancroft | Barley | 1 | 7 | | | | | | | |
| Pst/Psh_5 | Bancroft | Barley | 2 | 3 | 4 | | | | | | |
| Pst/Psh_5 | Bigo | Barley | 1 | 6 | | | | | | | |
| Pst/Psh_5 | Bigo | Barley | 2 | 6 | | | | | | | |
| Pst/Psh_5 | Emir | Barley | 1 | | | | | | | 2 | 4 |
| Pst/Psh_5 | Emir | Barley | 2 | | | | | | | 1 | 5 |
| Pst/Psh_5 | Heils Franken | Barley | 1 | 4 | 4 | | | | | | |
| Pst/Psh_5 | Heils Franken | Barley | 2 | 4 | 4 | | | | | | |

| | | | | | | | | | | | |
|---|---|---|---|---|---|---|---|---|---|---|---|
| Pst/Psh_5 | Hiproly | Barley | 1 | 7 | | | | | | | |
| Pst/Psh_5 | Hiproly | Barley | 2 | 7 | | | | | | | |
| Pst/Psh_5 | Hordeum spontaneum | Barley | 1 | | | | | | 3 | 3 | |
| Pst/Psh_5 | Hordeum spontaneum | Barley | 2 | | | | | | 3 | 3 | |
| Pst/Psh_5 | I5 | Barley | 1 | 7 | | | | | | | |
| Pst/Psh_5 | I5 | Barley | 2 | 7 | | | | | | | |
| Pst/Psh_5 | Trumpf | Barley | 1 | 7 | | | | | | | |
| Pst/Psh_5 | Trumpf | Barley | 2 | 7 | | | | | | | |
| Pst/Psh_5 | Varanda | Barley | 1 | | | | | | 3 | 3 | |
| Pst/Psh_5 | Varanda | Barley | 2 | | | | | | 4 | 2 | |
| Pst/Psh_5 | Aegilops tauschii | Other | 1 | 7 | | | | | | | |
| Pst/Psh_5 | Aegilops tauschii | Other | 2 | 7 | | | | | | | |
| Pst/Psh_5 | Elymus repens | Other | 1 | 7 | | | | | | | |
| Pst/Psh_5 | Elymus repens | Other | 2 | 7 | | | | | | | |
| Pst/Psh_5 | Avocet S | Wheat | 1 | | | 5 | 2 | | | | |
| Pst/Psh_5 | Avocet S | Wheat | 2 | | | | | 4 | 3 | | |
| Pst/Psh_5 | Avocet SP | Wheat | 1 | 7 | | | | | | | |
| Pst/Psh_5 | Avocet SP | Wheat | 2 | 7 | | | | | | | |
| Pst/Psh_5 | Avocet Yr1 | Wheat | 1 | 4 | 4 | | | | | | |
| Pst/Psh_5 | Avocet Yr1 | Wheat | 2 | 4 | 4 | | | | | | |
| Pst/Psh_5 | Avocet Yr15 | Wheat | 1 | 7 | | | | | | | |
| Pst/Psh_5 | Avocet Yr15 | Wheat | 2 | 7 | | | | | | | |
| Pst/Psh_5 | Avocet Yr17 | Wheat | 1 | 6 | | | | | | | |
| Pst/Psh_5 | Avocet Yr17 | Wheat | 2 | 6 | | | | | | | |
| Pst/Psh_5 | Avocet Yr24 | Wheat | 1 | 7 | | | | | | | |
| Pst/Psh_5 | Avocet Yr24 | Wheat | 2 | 7 | | | | | | | |
| Pst/Psh_5 | Avocet Yr27 | Wheat | 1 | 7 | | | | | | | |
| Pst/Psh_5 | Avocet Yr27 | Wheat | 2 | 7 | | | | | | | |
| Pst/Psh_5 | Avocet Yr32 | Wheat | 1 | 7 | | | | | | | |
| Pst/Psh_5 | Avocet Yr32 | Wheat | 2 | | 7 | | | | | | |
| Pst/Psh_5 | Avocet Yr5 | Wheat | 1 | 8 | | | | | | | |
| Pst/Psh_5 | Avocet Yr5 | Wheat | 2 | 8 | | | | | | | |
| Pst/Psh_5 | Avocet Yr6 | Wheat | 1 | | | 4 | 2 | | | | |
| Pst/Psh_5 | Avocet Yr6 | Wheat | 2 | | | | 2 | 4 | | | |
| Pst/Psh_5 | Avocet Yr7 | Wheat | 1 | 6 | | | | | | | |
| Pst/Psh_5 | Avocet Yr7 | Wheat | 2 | | 6 | | | | | | |
| Pst/Psh_5 | Avocet Yr8 | Wheat | 1 | 7 | | | | | | | |
| Pst/Psh_5 | Avocet Yr8 | Wheat | 2 | 7 | | | | | | | |
| Pst/Psh_5 | Avocet Yr9 | Wheat | 1 | | | 3 | 3 | | | | |
| Pst/Psh_5 | Avocet Yr9 | Wheat | 2 | | | | | 3 | 3 | | |
| Pst/Psh_5 | Chinese 166 | Wheat | 1 | 8 | | | | | | | |
| Pst/Psh_5 | Chinese 166 | Wheat | 2 | 4 | 4 | | | | | | |
| Pst/Psh_5 | Compair | Wheat | 1 | 7 | | | | | | | |
| Pst/Psh_5 | Compair | Wheat | 2 | 7 | | | | | | | |
| Pst/Psh_5 | Hybrid 46 | Wheat | 1 | 8 | | | | | | | |
| Pst/Psh_5 | Hybrid 46 | Wheat | 2 | 8 | | | | | | | |
| Pst/Psh_5 | Kalyansona | Wheat | 1 | | | | | | 3 | 3 | |
| Pst/Psh_5 | Kalyansona | Wheat | 2 | | | | | | | 4 | 2 |
| Pst/Psh_5 | Lee | Wheat | 1 | | 6 | | | | | | |
| Pst/Psh_5 | Lee | Wheat | 2 | | | 6 | | | | | |

| | | | | | | | | | | | |
|---|---|---|---|---|---|---|---|---|---|---|---|
| Pst/Psh_5 | Moro | Wheat | 1 | 7 | | | | | | | |
| Pst/Psh_5 | Moro | Wheat | 2 | 7 | | | | | | | |
| Pst/Psh_5 | Morocco | Wheat | 1 | | | | | | | 5 | 2 |
| Pst/Psh_5 | Morocco | Wheat | 2 | | | | | | | | 7 |
| Pst/Psh_5 | Sleipner | Wheat | 1 | 8 | | | | | | | |
| Pst/Psh_5 | Sleipner | Wheat | 2 | 8 | | | | | | | |
| Pst/Psh_5 | Suwon/Omar | Wheat | 1 | 7 | | | | | | | |
| Pst/Psh_5 | Suwon/Omar | Wheat | 2 | 7 | | | | | | | |
| Pst/Psh_5 | TP 981 | Wheat | 1 | 3 | 3 | | | | | | |
| Pst/Psh_5 | TP 981 | Wheat | 2 | 3 | 3 | | | | | | |
| Pst/Psh_5 | Triticum dicoccon | Wheat | 1 | 8 | | | | | | | |
| Pst/Psh_5 | Triticum dicoccon | Wheat | 2 | 8 | | | | | | | |
| Pst/Psh_5 | Triticum monococcum | Wheat | 1 | 7 | | | | | | | |
| Pst/Psh_5 | Triticum monococcum | Wheat | 2 | 7 | | | | | | | |
| Pst/Psh_5 | Vilmorin 23 | Wheat | 1 | 7 | | | | | | | |
| Pst/Psh_5 | Vilmorin 23 | Wheat | 2 | 7 | | | | | | | |
| Pst/Psh_5 | VPM1 | Wheat | 1 | 4 | 4 | | | | | | |
| Pst/Psh_5 | VPM1 | Wheat | 2 | 4 | 4 | | | | | | |
| Pst/Psh_9 | Abed Binder | Barley | 1 | 7 | | | | | | | |
| Pst/Psh_9 | Abed Binder | Barley | 2 | 7 | | | | | | | |
| Pst/Psh_9 | Afzal | Barley | 1 | | | | | | | 2 | 5 |
| Pst/Psh_9 | Afzal | Barley | 2 | | | | | | | | 7 |
| Pst/Psh_9 | Astrix | Barley | 1 | 7 | | | | | | | |
| Pst/Psh_9 | Astrix | Barley | 2 | 7 | | | | | | | |
| Pst/Psh_9 | Bancroft | Barley | 1 | 8 | | | | | | | |
| Pst/Psh_9 | Bancroft | Barley | 2 | 8 | | | | | | | |
| Pst/Psh_9 | Bigo | Barley | 1 | 6 | | | | | | | |
| Pst/Psh_9 | Bigo | Barley | 2 | 6 | | | | | | | |
| Pst/Psh_9 | Emir | Barley | 1 | | | | | | | 2 | 5 |
| Pst/Psh_9 | Emir | Barley | 2 | | | | | | | 1 | 6 |
| Pst/Psh_9 | Heils Franken | Barley | 1 | 8 | | | | | | | |
| Pst/Psh_9 | Heils Franken | Barley | 2 | 4 | 4 | | | | | | |
| Pst/Psh_9 | Hiproly | Barley | 1 | 8 | | | | | | | |
| Pst/Psh_9 | Hiproly | Barley | 2 | 8 | | | | | | | |
| Pst/Psh_9 | Hordeum spontaneum | Barley | 1 | | | | | 3 | 3 | 1 | |
| Pst/Psh_9 | Hordeum spontaneum | Barley | 2 | | | | | 3 | 3 | 1 | |
| Pst/Psh_9 | I5 | Barley | 1 | 8 | | | | | | | |
| Pst/Psh_9 | I5 | Barley | 2 | 8 | | | | | | | |
| Pst/Psh_9 | Trumpf | Barley | 1 | 6 | | | | | | | |
| Pst/Psh_9 | Trumpf | Barley | 2 | 6 | | | | | | | |
| Pst/Psh_9 | Varanda | Barley | 1 | 7 | | | | | | | |
| Pst/Psh_9 | Varanda | Barley | 2 | 7 | | | | | | | |
| Pst/Psh_9 | Aegilops tauschii | Other | 1 | 8 | | | | | | | |
| Pst/Psh_9 | Aegilops tauschii | Other | 2 | 8 | | | | | | | |
| Pst/Psh_9 | Elymus repens | Other | 1 | 6 | | | | | | | |
| Pst/Psh_9 | Elymus repens | Other | 2 | 6 | | | | | | | |
| Pst/Psh_9 | Avocet S | Wheat | 1 | | 3 | 4 | | | | | |
| Pst/Psh_9 | Avocet S | Wheat | 2 | | | 4 | 3 | | | | |
| Pst/Psh_9 | Avocet SP | Wheat | 1 | 8 | | | | | | | |
| Pst/Psh_9 | Avocet SP | Wheat | 2 | 8 | | | | | | | |

| | | | | | | | | | | | |
|---|---|---|---|---|---|---|---|---|---|---|---|
| Pst/Psh_9 | Avocet Yr1 | Wheat | 1 | 6 | | | | | | | |
| Pst/Psh_9 | Avocet Yr1 | Wheat | 2 | | 6 | | | | | | |
| Pst/Psh_9 | Avocet Yr15 | Wheat | 1 | 8 | | | | | | | |
| Pst/Psh_9 | Avocet Yr15 | Wheat | 2 | 8 | | | | | | | |
| Pst/Psh_9 | Avocet Yr17 | Wheat | 1 | 7 | | | | | | | |
| Pst/Psh_9 | Avocet Yr17 | Wheat | 2 | 7 | | | | | | | |
| Pst/Psh_9 | Avocet Yr24 | Wheat | 1 | 8 | | | | | | | |
| Pst/Psh_9 | Avocet Yr24 | Wheat | 2 | 8 | | | | | | | |
| Pst/Psh_9 | Avocet Yr27 | Wheat | 1 | 7 | | | | | | | |
| Pst/Psh_9 | Avocet Yr27 | Wheat | 2 | 7 | | | | | | | |
| Pst/Psh_9 | Avocet Yr32 | Wheat | 1 | 7 | | | | | | | |
| Pst/Psh_9 | Avocet Yr32 | Wheat | 2 | | 7 | | | | | | |
| Pst/Psh_9 | Avocet Yr5 | Wheat | 1 | 7 | | | | | | | |
| Pst/Psh_9 | Avocet Yr5 | Wheat | 2 | 7 | | | | | | | |
| Pst/Psh_9 | Avocet Yr6 | Wheat | 1 | 8 | | | | | | | |
| Pst/Psh_9 | Avocet Yr6 | Wheat | 2 | 4 | 4 | | | | | | |
| Pst/Psh_9 | Avocet Yr7 | Wheat | 1 | 7 | | | | | | | |
| Pst/Psh_9 | Avocet Yr7 | Wheat | 2 | | 7 | | | | | | |
| Pst/Psh_9 | Avocet Yr8 | Wheat | 1 | 7 | | | | | | | |
| Pst/Psh_9 | Avocet Yr8 | Wheat | 2 | 7 | | | | | | | |
| Pst/Psh_9 | Avocet Yr9 | Wheat | 1 | 4 | 4 | | | | | | |
| Pst/Psh_9 | Avocet Yr9 | Wheat | 2 | | 4 | 4 | | | | | |
| Pst/Psh_9 | Chinese 166 | Wheat | 1 | | | | | | | | 6 |
| Pst/Psh_9 | Chinese 166 | Wheat | 2 | | | | | | | | 6 |
| Pst/Psh_9 | Compair | Wheat | 1 | 8 | | | | | | | |
| Pst/Psh_9 | Compair | Wheat | 2 | 8 | | | | | | | |
| Pst/Psh_9 | Hybrid 46 | Wheat | 1 | 8 | | | | | | | |
| Pst/Psh_9 | Hybrid 46 | Wheat | 2 | 8 | | | | | | | |
| Pst/Psh_9 | Kalyansona | Wheat | 1 | | 3 | 3 | | | | | |
| Pst/Psh_9 | Kalyansona | Wheat | 2 | | 3 | 3 | | | | | |
| Pst/Psh_9 | Lee | Wheat | 1 | | 4 | 4 | | | | | |
| Pst/Psh_9 | Lee | Wheat | 2 | | | 8 | | | | | |
| Pst/Psh_9 | Moro | Wheat | 1 | 7 | | | | | | | |
| Pst/Psh_9 | Moro | Wheat | 2 | 7 | | | | | | | |
| Pst/Psh_9 | Morocco | Wheat | 1 | | | | 3 | 3 | | | |
| Pst/Psh_9 | Morocco | Wheat | 2 | | | | | 2 | 2 | 2 | |
| Pst/Psh_9 | Sleipner | Wheat | 1 | 7 | | | | | | | |
| Pst/Psh_9 | Sleipner | Wheat | 2 | | 7 | | | | | | |
| Pst/Psh_9 | Suwon/Omar | Wheat | 1 | 8 | | | | | | | |
| Pst/Psh_9 | Suwon/Omar | Wheat | 2 | 8 | | | | | | | |
| Pst/Psh_9 | TP 981 | Wheat | 1 | 8 | | | | | | | |
| Pst/Psh_9 | TP 981 | Wheat | 2 | 8 | | | | | | | |
| Pst/Psh_9 | Triticum dicoccon | Wheat | 1 | 8 | | | | | | | |
| Pst/Psh_9 | Triticum dicoccon | Wheat | 2 | 8 | | | | | | | |
| Pst/Psh_9 | Triticum monococcum | Wheat | 1 | 6 | | | | | | | |
| Pst/Psh_9 | Triticum monococcum | Wheat | 2 | 6 | | | | | | | |
| Pst/Psh_9 | Vilmorin 23 | Wheat | 1 | 8 | | | | | | | |
| Pst/Psh_9 | Vilmorin 23 | Wheat | 2 | 8 | | | | | | | |
| Pst/Psh_9 | VPM1 | Wheat | 1 | 8 | | | | | | | |
| Pst/Psh_9 | VPM1 | Wheat | 2 | 8 | | | | | | | |

| | | | | | | | | | | | |
|---|---|---|---|---|---|---|---|---|---|---|---|
| Psh/Pst_13 | Abed Binder | Barley | 1 | 7 | | | | | | | |
| Psh/Pst_13 | Abed Binder | Barley | 2 | 7 | | | | | | | |
| Psh/Pst_13 | Afzal | Barley | 1 | | | | | | | 2 | 4 |
| Psh/Pst_13 | Afzal | Barley | 2 | | | | | | | 1 | 5 |
| Psh/Pst_13 | Astrix | Barley | 1 | 7 | | | | | | | |
| Psh/Pst_13 | Astrix | Barley | 2 | 3 | 4 | | | | | | |
| Psh/Pst_13 | Bancroft | Barley | 1 | 7 | | | | | | | |
| Psh/Pst_13 | Bancroft | Barley | 2 | 7 | | | | | | | |
| Psh/Pst_13 | Bigo | Barley | 1 | 8 | | | | | | | |
| Psh/Pst_13 | Bigo | Barley | 2 | 4 | 4 | | | | | | |
| Psh/Pst_13 | Emir | Barley | 1 | | | | | | | 2 | 4 |
| Psh/Pst_13 | Emir | Barley | 2 | | | | | | | | 6 |
| Psh/Pst_13 | Heils Franken | Barley | 1 | 4 | 4 | | | | | | |
| Psh/Pst_13 | Heils Franken | Barley | 2 | 4 | 4 | | | | | | |
| Psh/Pst_13 | Hiproly | Barley | 1 | 7 | | | | | | | |
| Psh/Pst_13 | Hiproly | Barley | 2 | 7 | | | | | | | |
| Psh/Pst_13 | Hordeum spontaneum | Barley | 1 | | | | | | 2 | 2 | 4 |
| Psh/Pst_13 | Hordeum spontaneum | Barley | 2 | | | | | | 2 | 2 | 4 |
| Psh/Pst_13 | I5 | Barley | 1 | 7 | | | | | | | |
| Psh/Pst_13 | I5 | Barley | 2 | 7 | | | | | | | |
| Psh/Pst_13 | Trumpf | Barley | 1 | 8 | | | | | | | |
| Psh/Pst_13 | Trumpf | Barley | 2 | 8 | | | | | | | |
| Psh/Pst_13 | Varanda | Barley | 1 | | | | | | | 3 | 3 |
| Psh/Pst_13 | Varanda | Barley | 2 | | | | | | | 3 | 3 |
| Psh/Pst_13 | Aegilops tauschii | Other | 1 | 8 | | | | | | | |
| Psh/Pst_13 | Aegilops tauschii | Other | 2 | 8 | | | | | | | |
| Psh/Pst_13 | Elymus repens | Other | 1 | 6 | | | | | | | |
| Psh/Pst_13 | Elymus repens | Other | 2 | 6 | | | | | | | |
| Psh/Pst_13 | Avocet S | Wheat | 1 | | | 4 | 2 | | | | |
| Psh/Pst_13 | Avocet S | Wheat | 2 | | | | 2 | 2 | 2 | | |
| Psh/Pst_13 | Avocet SP | Wheat | 1 | 8 | | | | | | | |
| Psh/Pst_13 | Avocet SP | Wheat | 2 | 8 | | | | | | | |
| Psh/Pst_13 | Avocet Yr1 | Wheat | 1 | 6 | | | | | | | |
| Psh/Pst_13 | Avocet Yr1 | Wheat | 2 | 6 | | | | | | | |
| Psh/Pst_13 | Avocet Yr15 | Wheat | 1 | 7 | | | | | | | |
| Psh/Pst_13 | Avocet Yr15 | Wheat | 2 | 7 | | | | | | | |
| Psh/Pst_13 | Avocet Yr17 | Wheat | 1 | 7 | | | | | | | |
| Psh/Pst_13 | Avocet Yr17 | Wheat | 2 | 7 | | | | | | | |
| Psh/Pst_13 | Avocet Yr24 | Wheat | 1 | 7 | | | | | | | |
| Psh/Pst_13 | Avocet Yr24 | Wheat | 2 | 7 | | | | | | | |
| Psh/Pst_13 | Avocet Yr27 | Wheat | 1 | 6 | | | | | | | |
| Psh/Pst_13 | Avocet Yr27 | Wheat | 2 | 6 | | | | | | | |
| Psh/Pst_13 | Avocet Yr32 | Wheat | 1 | 8 | | | | | | | |
| Psh/Pst_13 | Avocet Yr32 | Wheat | 2 | | 8 | | | | | | |
| Psh/Pst_13 | Avocet Yr5 | Wheat | 1 | 7 | | | | | | | |
| Psh/Pst_13 | Avocet Yr5 | Wheat | 2 | 7 | | | | | | | |
| Psh/Pst_13 | Avocet Yr6 | Wheat | 1 | 7 | | | | | | | |
| Psh/Pst_13 | Avocet Yr6 | Wheat | 2 | | 7 | | | | | | |
| Psh/Pst_13 | Avocet Yr7 | Wheat | 1 | 6 | | | | | | | |
| Psh/Pst_13 | Avocet Yr7 | Wheat | 2 | | 6 | | | | | | |

| | | | | | | | | | | | |
|---|---|---|---|---|---|---|---|---|---|---|---|
| Psh/Pst_13 | Avocet Yr8 | Wheat | 1 | | 7 | | | | | | |
| Psh/Pst_13 | Avocet Yr8 | Wheat | 2 | | 7 | | | | | | |
| Psh/Pst_13 | Avocet Yr9 | Wheat | 1 | | | | 3 | 2 | 1 | | |
| Psh/Pst_13 | Avocet Yr9 | Wheat | 2 | | | | 1 | 2 | 2 | 1 | |
| Psh/Pst_13 | Chinese 166 | Wheat | 1 | 6 | | | | | | | |
| Psh/Pst_13 | Chinese 166 | Wheat | 2 | 6 | | | | | | | |
| Psh/Pst_13 | Compair | Wheat | 1 | 8 | | | | | | | |
| Psh/Pst_13 | Compair | Wheat | 2 | 8 | | | | | | | |
| Psh/Pst_13 | Hybrid 46 | Wheat | 1 | 7 | | | | | | | |
| Psh/Pst_13 | Hybrid 46 | Wheat | 2 | 7 | | | | | | | |
| Psh/Pst_13 | Kalyansona | Wheat | 1 | | | 2 | 2 | 2 | | | |
| Psh/Pst_13 | Kalyansona | Wheat | 2 | | | | 3 | 3 | | | |
| Psh/Pst_13 | Lee | Wheat | 1 | | 4 | 4 | | | | | |
| Psh/Pst_13 | Lee | Wheat | 2 | | 4 | 4 | | | | | |
| Psh/Pst_13 | Moro | Wheat | 1 | 8 | | | | | | | |
| Psh/Pst_13 | Moro | Wheat | 2 | 8 | | | | | | | |
| Psh/Pst_13 | Morocco | Wheat | 1 | | | | 2 | 3 | 1 | | |
| Psh/Pst_13 | Morocco | Wheat | 2 | | | | 2 | 2 | 2 | | |
| Psh/Pst_13 | Sleipner | Wheat | 1 | 7 | | | | | | | |
| Psh/Pst_13 | Sleipner | Wheat | 2 | | 7 | | | | | | |
| Psh/Pst_13 | Suwon/Omar | Wheat | 1 | 7 | | | | | | | |
| Psh/Pst_13 | Suwon/Omar | Wheat | 2 | 7 | | | | | | | |
| Psh/Pst_13 | TP 981 | Wheat | 1 | 7 | | | | | | | |
| Psh/Pst_13 | TP 981 | Wheat | 2 | 7 | | | | | | | |
| Psh/Pst_13 | Triticum dicoccon | Wheat | 1 | 8 | | | | | | | |
| Psh/Pst_13 | Triticum dicoccon | Wheat | 2 | 8 | | | | | | | |
| Psh/Pst_13 | Triticum monococcum | Wheat | 1 | 7 | | | | | | | |
| Psh/Pst_13 | Triticum monococcum | Wheat | 2 | 7 | | | | | | | |
| Psh/Pst_13 | Vilmorin 23 | Wheat | 1 | 6 | | | | | | | |
| Psh/Pst_13 | Vilmorin 23 | Wheat | 2 | 6 | | | | | | | |
| Psh/Pst_13 | VPM1 | Wheat | 1 | 7 | | | | | | | |
| Psh/Pst_13 | VPM1 | Wheat | 2 | 7 | | | | | | | |
| Pst/Psh_7 | Abed Binder | Barley | 1 | 7 | | | | | | | |
| Pst/Psh_7 | Abed Binder | Barley | 2 | 4 | 3 | | | | | | |
| Pst/Psh_7 | Afzal | Barley | 1 | | | | | | | 1 | 6 |
| Pst/Psh_7 | Afzal | Barley | 2 | | | | | | | | 7 |
| Pst/Psh_7 | Astrix | Barley | 1 | 7 | | | | | | | |
| Pst/Psh_7 | Astrix | Barley | 2 | 7 | | | | | | | |
| Pst/Psh_7 | Bancroft | Barley | 1 | 8 | | | | | | | |
| Pst/Psh_7 | Bancroft | Barley | 2 | 8 | | | | | | | |
| Pst/Psh_7 | Bigo | Barley | 1 | 8 | | | | | | | |
| Pst/Psh_7 | Bigo | Barley | 2 | 8 | | | | | | | |
| Pst/Psh_7 | Emir | Barley | 1 | | | | | | | 2 | 5 |
| Pst/Psh_7 | Emir | Barley | 2 | | | | | | | 1 | 6 |
| Pst/Psh_7 | Heils Franken | Barley | 1 | 8 | | | | | | | |
| Pst/Psh_7 | Heils Franken | Barley | 2 | 8 | | | | | | | |
| Pst/Psh_7 | Hiproly | Barley | 1 | 8 | | | | | | | |
| Pst/Psh_7 | Hiproly | Barley | 2 | 8 | | | | | | | |
| Pst/Psh_7 | Hordeum spontaneum | Barley | 1 | | | | | | | 2 | 5 |
| Pst/Psh_7 | Hordeum spontaneum | Barley | 2 | | | | | | | 2 | 5 |

| | | | | | | | | | | | |
|---|---|---|---|---|---|---|---|---|---|---|---|
| Pst/Psh_7 | I5 | Barley | 1 | 7 | | | | | | | |
| Pst/Psh_7 | I5 | Barley | 2 | 7 | | | | | | | |
| Pst/Psh_7 | Trumpf | Barley | 1 | 7 | | | | | | | |
| Pst/Psh_7 | Trumpf | Barley | 2 | 7 | | | | | | | |
| Pst/Psh_7 | Varanda | Barley | 1 | | | 2 | 2 | 2 | | | |
| Pst/Psh_7 | Varanda | Barley | 2 | | | 2 | 2 | 2 | | | |
| Pst/Psh_7 | Aegilops tauschii | Other | 1 | 7 | | | | | | | |
| Pst/Psh_7 | Aegilops tauschii | Other | 2 | 7 | | | | | | | |
| Pst/Psh_7 | Elymus repens | Other | 1 | 6 | | | | | | | |
| Pst/Psh_7 | Elymus repens | Other | 2 | 6 | | | | | | | |
| Pst/Psh_7 | Avocet S | Wheat | 1 | | 3 | 3 | | | | | |
| Pst/Psh_7 | Avocet S | Wheat | 2 | | | 1 | 2 | 1 | 2 | | |
| Pst/Psh_7 | Avocet SP | Wheat | 1 | 7 | | | | | | | |
| Pst/Psh_7 | Avocet SP | Wheat | 2 | 7 | | | | | | | |
| Pst/Psh_7 | Avocet Yr1 | Wheat | 1 | 6 | | | | | | | |
| Pst/Psh_7 | Avocet Yr1 | Wheat | 2 | | 6 | | | | | | |
| Pst/Psh_7 | Avocet Yr15 | Wheat | 1 | 8 | | | | | | | |
| Pst/Psh_7 | Avocet Yr15 | Wheat | 2 | 8 | | | | | | | |
| Pst/Psh_7 | Avocet Yr17 | Wheat | 1 | 7 | | | | | | | |
| Pst/Psh_7 | Avocet Yr17 | Wheat | 2 | 7 | | | | | | | |
| Pst/Psh_7 | Avocet Yr24 | Wheat | 1 | 8 | | | | | | | |
| Pst/Psh_7 | Avocet Yr24 | Wheat | 2 | 8 | | | | | | | |
| Pst/Psh_7 | Avocet Yr27 | Wheat | 1 | 7 | | | | | | | |
| Pst/Psh_7 | Avocet Yr27 | Wheat | 2 | 7 | | | | | | | |
| Pst/Psh_7 | Avocet Yr32 | Wheat | 1 | 4 | 4 | | | | | | |
| Pst/Psh_7 | Avocet Yr32 | Wheat | 2 | 4 | 4 | | | | | | |
| Pst/Psh_7 | Avocet Yr5 | Wheat | 1 | 8 | | | | | | | |
| Pst/Psh_7 | Avocet Yr5 | Wheat | 2 | 8 | | | | | | | |
| Pst/Psh_7 | Avocet Yr6 | Wheat | 1 | 7 | | | | | | | |
| Pst/Psh_7 | Avocet Yr6 | Wheat | 2 | | 7 | | | | | | |
| Pst/Psh_7 | Avocet Yr7 | Wheat | 1 | 6 | | | | | | | |
| Pst/Psh_7 | Avocet Yr7 | Wheat | 2 | | 6 | | | | | | |
| Pst/Psh_7 | Avocet Yr8 | Wheat | 1 | | 4 | 4 | | | | | |
| Pst/Psh_7 | Avocet Yr8 | Wheat | 2 | | 4 | 4 | | | | | |
| Pst/Psh_7 | Avocet Yr9 | Wheat | 1 | | 2 | 2 | 2 | 1 | | | |
| Pst/Psh_7 | Avocet Yr9 | Wheat | 2 | | | 2 | 3 | 2 | | | |
| Pst/Psh_7 | Chinese 166 | Wheat | 1 | | | | | | | | 8 |
| Pst/Psh_7 | Chinese 166 | Wheat | 2 | | | | | | | | 8 |
| Pst/Psh_7 | Compair | Wheat | 1 | 8 | | | | | | | |
| Pst/Psh_7 | Compair | Wheat | 2 | 8 | | | | | | | |
| Pst/Psh_7 | Hybrid 46 | Wheat | 1 | 8 | | | | | | | |
| Pst/Psh_7 | Hybrid 46 | Wheat | 2 | 8 | | | | | | | |
| Pst/Psh_7 | Kalyansona | Wheat | 1 | | | | | | | 4 | 2 |
| Pst/Psh_7 | Kalyansona | Wheat | 2 | | | | | | | 3 | 3 |
| Pst/Psh_7 | Lee | Wheat | 1 | | | 7 | | | | | |
| Pst/Psh_7 | Lee | Wheat | 2 | | | 7 | | | | | |
| Pst/Psh_7 | Moro | Wheat | 1 | 7 | | | | | | | |
| Pst/Psh_7 | Moro | Wheat | 2 | 7 | | | | | | | |
| Pst/Psh_7 | Morocco | Wheat | 1 | | | | | | | | 7 |
| Pst/Psh_7 | Morocco | Wheat | 2 | | | | | | | | 7 |

| | | | | | | | | | | | |
|---|---|---|---|---|---|---|---|---|---|---|---|
| Pst/Psh_7 | Sleipner | Wheat | 1 | 8 | | | | | | | |
| Pst/Psh_7 | Sleipner | Wheat | 2 | | 8 | | | | | | |
| Pst/Psh_7 | Suwon/Omar | Wheat | 1 | 8 | | | | | | | |
| Pst/Psh_7 | Suwon/Omar | Wheat | 2 | 8 | | | | | | | |
| Pst/Psh_7 | TP 981 | Wheat | 1 | 6 | | | | | | | |
| Pst/Psh_7 | TP 981 | Wheat | 2 | 6 | | | | | | | |
| Pst/Psh_7 | Triticum dicoccon | Wheat | 1 | 7 | | | | | | | |
| Pst/Psh_7 | Triticum dicoccon | Wheat | 2 | 7 | | | | | | | |
| Pst/Psh_7 | Triticum monococcum | Wheat | 1 | 7 | | | | | | | |
| Pst/Psh_7 | Triticum monococcum | Wheat | 2 | 7 | | | | | | | |
| Pst/Psh_7 | Vilmorin 23 | Wheat | 1 | 8 | | | | | | | |
| Pst/Psh_7 | Vilmorin 23 | Wheat | 2 | 8 | | | | | | | |
| Pst/Psh_7 | VPM1 | Wheat | 1 | 8 | | | | | | | |
| Pst/Psh_7 | VPM1 | Wheat | 2 | 8 | | | | | | | |
| Pst/Psh_15 | Abed Binder | Barley | 1 | 4 | 4 | | | | | | |
| Pst/Psh_15 | Abed Binder | Barley | 2 | 4 | 4 | | | | | | |
| Pst/Psh_15 | Afzal | Barley | 1 | | | | | | | | 6 |
| Pst/Psh_15 | Afzal | Barley | 2 | | | | | | | | 6 |
| Pst/Psh_15 | Astrix | Barley | 1 | 7 | | | | | | | |
| Pst/Psh_15 | Astrix | Barley | 2 | 7 | | | | | | | |
| Pst/Psh_15 | Bancroft | Barley | 1 | 8 | | | | | | | |
| Pst/Psh_15 | Bancroft | Barley | 2 | 4 | 4 | | | | | | |
| Pst/Psh_15 | Bigo | Barley | 1 | 7 | | | | | | | |
| Pst/Psh_15 | Bigo | Barley | 2 | 7 | | | | | | | |
| Pst/Psh_15 | Emir | Barley | 1 | | | | | | | 1 | 6 |
| Pst/Psh_15 | Emir | Barley | 2 | | | | | | | 1 | 6 |
| Pst/Psh_15 | Heils Franken | Barley | 1 | 7 | | | | | | | |
| Pst/Psh_15 | Heils Franken | Barley | 2 | 7 | | | | | | | |
| Pst/Psh_15 | Hiproly | Barley | 1 | 7 | | | | | | | |
| Pst/Psh_15 | Hiproly | Barley | 2 | 7 | | | | | | | |
| Pst/Psh_15 | Hordeum spontaneum | Barley | 1 | | | | | | 3 | 3 | 1 |
| Pst/Psh_15 | Hordeum spontaneum | Barley | 2 | | | | | | 3 | 3 | 1 |
| Pst/Psh_15 | I5 | Barley | 1 | 8 | | | | | | | |
| Pst/Psh_15 | I5 | Barley | 2 | 8 | | | | | | | |
| Pst/Psh_15 | Trumpf | Barley | 1 | 8 | | | | | | | |
| Pst/Psh_15 | Trumpf | Barley | 2 | 8 | | | | | | | |
| Pst/Psh_15 | Varanda | Barley | 1 | | | 6 | | | | | |
| Pst/Psh_15 | Varanda | Barley | 2 | | | | 6 | | | | |
| Pst/Psh_15 | Aegilops tauschii | Other | 1 | 8 | | | | | | | |
| Pst/Psh_15 | Aegilops tauschii | Other | 2 | 8 | | | | | | | |
| Pst/Psh_15 | Elymus repens | Other | 1 | 6 | | | | | | | |
| Pst/Psh_15 | Elymus repens | Other | 2 | 6 | | | | | | | |
| Pst/Psh_15 | Avocet S | Wheat | 1 | | 3 | 4 | | | | | |
| Pst/Psh_15 | Avocet S | Wheat | 2 | | 5 | 2 | | | | | |
| Pst/Psh_15 | Avocet SP | Wheat | 1 | 8 | | | | | | | |
| Pst/Psh_15 | Avocet SP | Wheat | 2 | 8 | | | | | | | |
| Pst/Psh_15 | Avocet Yr1 | Wheat | 1 | 6 | | | | | | | |
| Pst/Psh_15 | Avocet Yr1 | Wheat | 2 | 6 | | | | | | | |
| Pst/Psh_15 | Avocet Yr15 | Wheat | 1 | 7 | | | | | | | |
| Pst/Psh_15 | Avocet Yr15 | Wheat | 2 | 7 | | | | | | | |

| | | | | | | | | | | | |
|---|---|---|---|---|---|---|---|---|---|---|---|
| Pst/Psh_15 | Avocet Yr17 | Wheat | 1 | 7 | | | | | | | |
| Pst/Psh_15 | Avocet Yr17 | Wheat | 2 | 7 | | | | | | | |
| Pst/Psh_15 | Avocet Yr24 | Wheat | 1 | 7 | | | | | | | |
| Pst/Psh_15 | Avocet Yr24 | Wheat | 2 | 7 | | | | | | | |
| Pst/Psh_15 | Avocet Yr27 | Wheat | 1 | 7 | | | | | | | |
| Pst/Psh_15 | Avocet Yr27 | Wheat | 2 | 7 | | | | | | | |
| Pst/Psh_15 | Avocet Yr32 | Wheat | 1 | 6 | | | | | | | |
| Pst/Psh_15 | Avocet Yr32 | Wheat | 2 | 6 | | | | | | | |
| Pst/Psh_15 | Avocet Yr5 | Wheat | 1 | 7 | | | | | | | |
| Pst/Psh_15 | Avocet Yr5 | Wheat | 2 | 7 | | | | | | | |
| Pst/Psh_15 | Avocet Yr6 | Wheat | 1 | 2 | 4 | 2 | | | | | |
| Pst/Psh_15 | Avocet Yr6 | Wheat | 2 | | 4 | 4 | | | | | |
| Pst/Psh_15 | Avocet Yr7 | Wheat | 1 | 6 | | | | | | | |
| Pst/Psh_15 | Avocet Yr7 | Wheat | 2 | 3 | 3 | | | | | | |
| Pst/Psh_15 | Avocet Yr8 | Wheat | 1 | 6 | 2 | | | | | | |
| Pst/Psh_15 | Avocet Yr8 | Wheat | 2 | 8 | | | | | | | |
| Pst/Psh_15 | Avocet Yr9 | Wheat | 1 | | | 3 | 2 | 1 | | | |
| Pst/Psh_15 | Avocet Yr9 | Wheat | 2 | | | 2 | 2 | 2 | | | |
| Pst/Psh_15 | Chinese 166 | Wheat | 1 | 4 | 3 | 1 | | | | | |
| Pst/Psh_15 | Chinese 166 | Wheat | 2 | 4 | 4 | | | | | | |
| Pst/Psh_15 | Compair | Wheat | 1 | 8 | | | | | | | |
| Pst/Psh_15 | Compair | Wheat | 2 | 8 | | | | | | | |
| Pst/Psh_15 | Hybrid 46 | Wheat | 1 | 7 | | | | | | | |
| Pst/Psh_15 | Hybrid 46 | Wheat | 2 | 7 | | | | | | | |
| Pst/Psh_15 | Kalyansona | Wheat | 1 | | | | | | | 3 | 3 |
| Pst/Psh_15 | Kalyansona | Wheat | 2 | | | | | | | 1 | 5 |
| Pst/Psh_15 | Lee | Wheat | 1 | | 3 | 4 | | | | | |
| Pst/Psh_15 | Lee | Wheat | 2 | | | 7 | | | | | |
| Pst/Psh_15 | Moro | Wheat | 1 | 8 | | | | | | | |
| Pst/Psh_15 | Moro | Wheat | 2 | 8 | | | | | | | |
| Pst/Psh_15 | Morocco | Wheat | 1 | | | | | | | | 6 |
| Pst/Psh_15 | Morocco | Wheat | 2 | | | | | | | | 6 |
| Pst/Psh_15 | Sleipner | Wheat | 1 | 8 | | | | | | | |
| Pst/Psh_15 | Sleipner | Wheat | 2 | 8 | | | | | | | |
| Pst/Psh_15 | Suwon/Omar | Wheat | 1 | 8 | | | | | | | |
| Pst/Psh_15 | Suwon/Omar | Wheat | 2 | 8 | | | | | | | |
| Pst/Psh_15 | TP 981 | Wheat | 1 | 7 | | | | | | | |
| Pst/Psh_15 | TP 981 | Wheat | 2 | 7 | | | | | | | |
| Pst/Psh_15 | Triticum dicoccon | Wheat | 1 | 7 | | | | | | | |
| Pst/Psh_15 | Triticum dicoccon | Wheat | 2 | 7 | | | | | | | |
| Pst/Psh_15 | Triticum monococcum | Wheat | 1 | 8 | | | | | | | |
| Pst/Psh_15 | Triticum monococcum | Wheat | 2 | 8 | | | | | | | |
| Pst/Psh_15 | Vilmorin 23 | Wheat | 1 | 7 | | | | | | | |
| Pst/Psh_15 | Vilmorin 23 | Wheat | 2 | 7 | | | | | | | |
| Pst/Psh_15 | VPM1 | Wheat | 1 | 6 | | | | | | | |
| Pst/Psh_15 | VPM1 | Wheat | 2 | 6 | | | | | | | |
| Pst/Psh_12 | Abed Binder | Barley | 1 | 7 | | | | | | | |
| Pst/Psh_12 | Abed Binder | Barley | 2 | 7 | | | | | | | |
| Pst/Psh_12 | Afzal | Barley | 1 | | | | | | | 2 | 4 |
| Pst/Psh_12 | Afzal | Barley | 2 | | | | | | | 1 | 5 |

| | | | | | | | | | | | |
|---|---|---|---|---|---|---|---|---|---|---|---|
| Pst/Psh_12 | Astrix | Barley | 1 | 7 | | | | | | | |
| Pst/Psh_12 | Astrix | Barley | 2 | 7 | | | | | | | |
| Pst/Psh_12 | Bancroft | Barley | 1 | 7 | | | | | | | |
| Pst/Psh_12 | Bancroft | Barley | 2 | 7 | | | | | | | |
| Pst/Psh_12 | Bigo | Barley | 1 | 7 | | | | | | | |
| Pst/Psh_12 | Bigo | Barley | 2 | 7 | | | | | | | |
| Pst/Psh_12 | Emir | Barley | 1 | | | | | | | 2 | 5 |
| Pst/Psh_12 | Emir | Barley | 2 | | | | | | | | 7 |
| Pst/Psh_12 | Heils Franken | Barley | 1 | 4 | 4 | | | | | | |
| Pst/Psh_12 | Heils Franken | Barley | 2 | | 8 | | | | | | |
| Pst/Psh_12 | Hiproly | Barley | 1 | 7 | | | | | | | |
| Pst/Psh_12 | Hiproly | Barley | 2 | 7 | | | | | | | |
| Pst/Psh_12 | Hordeum spontaneum | Barley | 1 | | | | | | | 3 | 3 |
| Pst/Psh_12 | Hordeum spontaneum | Barley | 2 | | | | | | | 3 | 3 |
| Pst/Psh_12 | I5 | Barley | 1 | 4 | 4 | | | | | | |
| Pst/Psh_12 | I5 | Barley | 2 | 4 | 4 | | | | | | |
| Pst/Psh_12 | Trumpf | Barley | 1 | 6 | | | | | | | |
| Pst/Psh_12 | Trumpf | Barley | 2 | 6 | | | | | | | |
| Pst/Psh_12 | Varanda | Barley | 1 | 7 | | | | | | | |
| Pst/Psh_12 | Varanda | Barley | 2 | 7 | | | | | | | |
| Pst/Psh_12 | Aegilops tauschii | Other | 1 | 8 | | | | | | | |
| Pst/Psh_12 | Aegilops tauschii | Other | 2 | 4 | 4 | | | | | | |
| Pst/Psh_12 | Elymus repens | Other | 1 | 6 | | | | | | | |
| Pst/Psh_12 | Elymus repens | Other | 2 | 6 | | | | | | | |
| Pst/Psh_12 | Avocet S | Wheat | 1 | | 3 | 3 | | | | | |
| Pst/Psh_12 | Avocet S | Wheat | 2 | | 3 | 2 | 1 | | | | |
| Pst/Psh_12 | Avocet SP | Wheat | 1 | 7 | | | | | | | |
| Pst/Psh_12 | Avocet SP | Wheat | 2 | 7 | | | | | | | |
| Pst/Psh_12 | Avocet Yr1 | Wheat | 1 | 7 | | | | | | | |
| Pst/Psh_12 | Avocet Yr1 | Wheat | 2 | 7 | | | | | | | |
| Pst/Psh_12 | Avocet Yr15 | Wheat | 1 | 7 | | | | | | | |
| Pst/Psh_12 | Avocet Yr15 | Wheat | 2 | 7 | | | | | | | |
| Pst/Psh_12 | Avocet Yr17 | Wheat | 1 | 6 | | | | | | | |
| Pst/Psh_12 | Avocet Yr17 | Wheat | 2 | 6 | | | | | | | |
| Pst/Psh_12 | Avocet Yr24 | Wheat | 1 | 6 | | | | | | | |
| Pst/Psh_12 | Avocet Yr24 | Wheat | 2 | 6 | | | | | | | |
| Pst/Psh_12 | Avocet Yr27 | Wheat | 1 | 8 | | | | | | | |
| Pst/Psh_12 | Avocet Yr27 | Wheat | 2 | 8 | | | | | | | |
| Pst/Psh_12 | Avocet Yr32 | Wheat | 1 | 7 | | | | | | | |
| Pst/Psh_12 | Avocet Yr32 | Wheat | 2 | | 7 | | | | | | |
| Pst/Psh_12 | Avocet Yr5 | Wheat | 1 | 8 | | | | | | | |
| Pst/Psh_12 | Avocet Yr5 | Wheat | 2 | 8 | | | | | | | |
| Pst/Psh_12 | Avocet Yr6 | Wheat | 1 | | 7 | | | | | | |
| Pst/Psh_12 | Avocet Yr6 | Wheat | 2 | | 4 | 2 | 1 | | | | |
| Pst/Psh_12 | Avocet Yr7 | Wheat | 1 | | 6 | | | | | | |
| Pst/Psh_12 | Avocet Yr7 | Wheat | 2 | | 6 | | | | | | |
| Pst/Psh_12 | Avocet Yr8 | Wheat | 1 | | 8 | | | | | | |
| Pst/Psh_12 | Avocet Yr8 | Wheat | 2 | | 8 | | | | | | |
| Pst/Psh_12 | Avocet Yr9 | Wheat | 1 | | | | 2 | 2 | 2 | | |
| Pst/Psh_12 | Avocet Yr9 | Wheat | 2 | | | 1 | 2 | 1 | 1 | 1 | |

| | | | | | | | | | | | |
|---|---|---|---|---|---|---|---|---|---|---|---|
| Pst/Psh_12 | Chinese 166 | Wheat | 1 | 8 | | | | | | | |
| Pst/Psh_12 | Chinese 166 | Wheat | 2 | | 8 | | | | | | |
| Pst/Psh_12 | Compair | Wheat | 1 | 8 | | | | | | | |
| Pst/Psh_12 | Compair | Wheat | 2 | 8 | | | | | | | |
| Pst/Psh_12 | Hybrid 46 | Wheat | 1 | 7 | | | | | | | |
| Pst/Psh_12 | Hybrid 46 | Wheat | 2 | 7 | | | | | | | |
| Pst/Psh_12 | Kalyansona | Wheat | 1 | | | 3 | 3 | | | | |
| Pst/Psh_12 | Kalyansona | Wheat | 2 | | | | | 2 | 1 | 3 | |
| Pst/Psh_12 | Lee | Wheat | 1 | | | 8 | | | | | |
| Pst/Psh_12 | Lee | Wheat | 2 | | | 8 | | | | | |
| Pst/Psh_12 | Moro | Wheat | 1 | 7 | | | | | | | |
| Pst/Psh_12 | Moro | Wheat | 2 | 7 | | | | | | | |
| Pst/Psh_12 | Morocco | Wheat | 1 | | | | | | | | 8 |
| Pst/Psh_12 | Morocco | Wheat | 2 | | | | | | | | 8 |
| Pst/Psh_12 | Sleipner | Wheat | 1 | | 7 | | | | | | |
| Pst/Psh_12 | Sleipner | Wheat | 2 | | 7 | | | | | | |
| Pst/Psh_12 | Suwon/Omar | Wheat | 1 | 7 | | | | | | | |
| Pst/Psh_12 | Suwon/Omar | Wheat | 2 | 7 | | | | | | | |
| Pst/Psh_12 | TP 981 | Wheat | 1 | 6 | | | | | | | |
| Pst/Psh_12 | TP 981 | Wheat | 2 | 6 | | | | | | | |
| Pst/Psh_12 | Triticum dicoccon | Wheat | 1 | 8 | | | | | | | |
| Pst/Psh_12 | Triticum dicoccon | Wheat | 2 | 8 | | | | | | | |
| Pst/Psh_12 | Triticum monococcum | Wheat | 1 | 7 | | | | | | | |
| Pst/Psh_12 | Triticum monococcum | Wheat | 2 | 7 | | | | | | | |
| Pst/Psh_12 | Vilmorin 23 | Wheat | 1 | 7 | | | | | | | |
| Pst/Psh_12 | Vilmorin 23 | Wheat | 2 | 7 | | | | | | | |
| Pst/Psh_12 | VPM1 | Wheat | 1 | 8 | | | | | | | |
| Pst/Psh_12 | VPM1 | Wheat | 2 | 8 | | | | | | | |
| Pst/Psh_14 | Abed Binder | Barley | 1 | 7 | | | | | | | |
| Pst/Psh_14 | Abed Binder | Barley | 2 | 3 | 4 | | | | | | |
| Pst/Psh_14 | Afzal | Barley | 1 | | | | | | | 2 | 5 |
| Pst/Psh_14 | Afzal | Barley | 2 | | | | | | | | 7 |
| Pst/Psh_14 | Astrix | Barley | 1 | 7 | | | | | | | |
| Pst/Psh_14 | Astrix | Barley | 2 | 7 | | | | | | | |
| Pst/Psh_14 | Bancroft | Barley | 1 | 8 | | | | | | | |
| Pst/Psh_14 | Bancroft | Barley | 2 | 8 | | | | | | | |
| Pst/Psh_14 | Bigo | Barley | 1 | 7 | | | | | | | |
| Pst/Psh_14 | Bigo | Barley | 2 | 7 | | | | | | | |
| Pst/Psh_14 | Emir | Barley | 1 | | | | | | | 2 | 4 |
| Pst/Psh_14 | Emir | Barley | 2 | | | | | | | | 6 |
| Pst/Psh_14 | Heils Franken | Barley | 1 | 7 | | | | | | | |
| Pst/Psh_14 | Heils Franken | Barley | 2 | 7 | | | | | | | |
| Pst/Psh_14 | Hiproly | Barley | 1 | 7 | | | | | | | |
| Pst/Psh_14 | Hiproly | Barley | 2 | 7 | | | | | | | |
| Pst/Psh_14 | Hordeum spontaneum | Barley | 1 | | | | | | 2 | 3 | 1 |
| Pst/Psh_14 | Hordeum spontaneum | Barley | 2 | | | | | | | 5 | 2 |
| Pst/Psh_14 | I5 | Barley | 1 | 8 | | | | | | | |
| Pst/Psh_14 | I5 | Barley | 2 | 4 | 4 | | | | | | |
| Pst/Psh_14 | Trumpf | Barley | 1 | 7 | | | | | | | |
| Pst/Psh_14 | Trumpf | Barley | 2 | 7 | | | | | | | |

| | | | | | | | | | | | |
|---|---|---|---|---|---|---|---|---|---|---|---|
| Pst/Psh_14 | Varanda | Barley | 1 | 7 | | | | | | | |
| Pst/Psh_14 | Varanda | Barley | 2 | 7 | | | | | | | |
| Pst/Psh_14 | Aegilops tauschii | Other | 1 | 8 | | | | | | | |
| Pst/Psh_14 | Aegilops tauschii | Other | 2 | 8 | | | | | | | |
| Pst/Psh_14 | Elymus repens | Other | 1 | 6 | | | | | | | |
| Pst/Psh_14 | Elymus repens | Other | 2 | 6 | | | | | | | |
| Pst/Psh_14 | Avocet S | Wheat | 1 | | 3 | 4 | | | | | |
| Pst/Psh_14 | Avocet S | Wheat | 2 | | | 3 | 4 | | | | |
| Pst/Psh_14 | Avocet SP | Wheat | 1 | 8 | | | | | | | |
| Pst/Psh_14 | Avocet SP | Wheat | 2 | 8 | | | | | | | |
| Pst/Psh_14 | Avocet Yr1 | Wheat | 1 | 3 | 3 | | | | | | |
| Pst/Psh_14 | Avocet Yr1 | Wheat | 2 | 3 | 3 | | | | | | |
| Pst/Psh_14 | Avocet Yr15 | Wheat | 1 | 7 | | | | | | | |
| Pst/Psh_14 | Avocet Yr15 | Wheat | 2 | 7 | | | | | | | |
| Pst/Psh_14 | Avocet Yr17 | Wheat | 1 | 7 | | | | | | | |
| Pst/Psh_14 | Avocet Yr17 | Wheat | 2 | 7 | | | | | | | |
| Pst/Psh_14 | Avocet Yr24 | Wheat | 1 | 8 | | | | | | | |
| Pst/Psh_14 | Avocet Yr24 | Wheat | 2 | 8 | | | | | | | |
| Pst/Psh_14 | Avocet Yr27 | Wheat | 1 | 7 | | | | | | | |
| Pst/Psh_14 | Avocet Yr27 | Wheat | 2 | 7 | | | | | | | |
| Pst/Psh_14 | Avocet Yr32 | Wheat | 1 | 7 | | | | | | | |
| Pst/Psh_14 | Avocet Yr32 | Wheat | 2 | 7 | | | | | | | |
| Pst/Psh_14 | Avocet Yr5 | Wheat | 1 | 7 | | | | | | | |
| Pst/Psh_14 | Avocet Yr5 | Wheat | 2 | 7 | | | | | | | |
| Pst/Psh_14 | Avocet Yr6 | Wheat | 1 | | 4 | 4 | | | | | |
| Pst/Psh_14 | Avocet Yr6 | Wheat | 2 | | | 2 | 2 | 2 | 2 | | |
| Pst/Psh_14 | Avocet Yr7 | Wheat | 1 | 6 | | | | | | | |
| Pst/Psh_14 | Avocet Yr7 | Wheat | 2 | 6 | | | | | | | |
| Pst/Psh_14 | Avocet Yr8 | Wheat | 1 | | | 7 | | | | | |
| Pst/Psh_14 | Avocet Yr8 | Wheat | 2 | | | 3 | 2 | 2 | | | |
| Pst/Psh_14 | Avocet Yr9 | Wheat | 1 | | | | 3 | 1 | 1 | 2 | |
| Pst/Psh_14 | Avocet Yr9 | Wheat | 2 | | | 1 | 1 | 2 | 3 | | |
| Pst/Psh_14 | Chinese 166 | Wheat | 1 | 8 | | | | | | | |
| Pst/Psh_14 | Chinese 166 | Wheat | 2 | 4 | 4 | | | | | | |
| Pst/Psh_14 | Compair | Wheat | 1 | 8 | | | | | | | |
| Pst/Psh_14 | Compair | Wheat | 2 | 8 | | | | | | | |
| Pst/Psh_14 | Hybrid 46 | Wheat | 1 | 7 | | | | | | | |
| Pst/Psh_14 | Hybrid 46 | Wheat | 2 | 7 | | | | | | | |
| Pst/Psh_14 | Kalyansona | Wheat | 1 | | | | | | 2 | 4 | |
| Pst/Psh_14 | Kalyansona | Wheat | 2 | | | | | | | 5 | 1 |
| Pst/Psh_14 | Lee | Wheat | 1 | | | 8 | | | | | |
| Pst/Psh_14 | Lee | Wheat | 2 | | | 8 | | | | | |
| Pst/Psh_14 | Moro | Wheat | 1 | 7 | | | | | | | |
| Pst/Psh_14 | Moro | Wheat | 2 | 7 | | | | | | | |
| Pst/Psh_14 | Morocco | Wheat | 1 | | | | | | | | 7 |
| Pst/Psh_14 | Morocco | Wheat | 2 | | | | | | | | 7 |
| Pst/Psh_14 | Sleipner | Wheat | 1 | 8 | | | | | | | |
| Pst/Psh_14 | Sleipner | Wheat | 2 | 8 | | | | | | | |
| Pst/Psh_14 | Suwon/Omar | Wheat | 1 | 7 | | | | | | | |
| Pst/Psh_14 | Suwon/Omar | Wheat | 2 | 7 | | | | | | | |

| | | | | | | | | | | | |
|---|---|---|---|---|---|---|---|---|---|---|---|
| Pst/Psh_14 | TP 981 | Wheat | 1 | 7 | | | | | | | |
| Pst/Psh_14 | TP 981 | Wheat | 2 | 7 | | | | | | | |
| Pst/Psh_14 | Triticum dicoccon | Wheat | 1 | 7 | | | | | | | |
| Pst/Psh_14 | Triticum dicoccon | Wheat | 2 | 7 | | | | | | | |
| Pst/Psh_14 | Triticum monococcum | Wheat | 1 | 7 | | | | | | | |
| Pst/Psh_14 | Triticum monococcum | Wheat | 2 | 7 | | | | | | | |
| Pst/Psh_14 | Vilmorin 23 | Wheat | 1 | 7 | | | | | | | |
| Pst/Psh_14 | Vilmorin 23 | Wheat | 2 | 7 | | | | | | | |
| Pst/Psh_14 | VPM1 | Wheat | 1 | 4 | 4 | | | | | | |
| Pst/Psh_14 | VPM1 | Wheat | 2 | 4 | 4 | | | | | | |
| Pst/Psh_16 | Abed Binder | Barley | 1 | 7 | | | | | | | |
| Pst/Psh_16 | Abed Binder | Barley | 2 | 7 | | | | | | | |
| Pst/Psh_16 | Afzal | Barley | 1 | | | | | | | 2 | 5 |
| Pst/Psh_16 | Afzal | Barley | 2 | | | | | | | | 7 |
| Pst/Psh_16 | Astrix | Barley | 1 | 4 | 4 | | | | | | |
| Pst/Psh_16 | Astrix | Barley | 2 | 4 | 4 | | | | | | |
| Pst/Psh_16 | Bancroft | Barley | 1 | 4 | 4 | | | | | | |
| Pst/Psh_16 | Bancroft | Barley | 2 | 8 | | | | | | | |
| Pst/Psh_16 | Bigo | Barley | 1 | 7 | | | | | | | |
| Pst/Psh_16 | Bigo | Barley | 2 | 7 | | | | | | | |
| Pst/Psh_16 | Emir | Barley | 1 | | | | | | | 2 | 5 |
| Pst/Psh_16 | Emir | Barley | 2 | | | | | | | | 7 |
| Pst/Psh_16 | Heils Franken | Barley | 1 | 8 | | | | | | | |
| Pst/Psh_16 | Heils Franken | Barley | 2 | 8 | | | | | | | |
| Pst/Psh_16 | Hiproly | Barley | 1 | 8 | | | | | | | |
| Pst/Psh_16 | Hiproly | Barley | 2 | 8 | | | | | | | |
| Pst/Psh_16 | Hordeum spontaneum | Barley | 1 | | | | | | 3 | 3 | |
| Pst/Psh_16 | Hordeum spontaneum | Barley | 2 | | | | | | | 3 | 3 |
| Pst/Psh_16 | I5 | Barley | 1 | 7 | | | | | | | |
| Pst/Psh_16 | I5 | Barley | 2 | 7 | | | | | | | |
| Pst/Psh_16 | Trumpf | Barley | 1 | 7 | | | | | | | |
| Pst/Psh_16 | Trumpf | Barley | 2 | 7 | | | | | | | |
| Pst/Psh_16 | Varanda | Barley | 1 | | | | 3 | 3 | | | |
| Pst/Psh_16 | Varanda | Barley | 2 | | | 2 | 2 | 2 | | | |
| Pst/Psh_16 | Aegilops tauschii | Other | 1 | 7 | | | | | | | |
| Pst/Psh_16 | Aegilops tauschii | Other | 2 | 7 | | | | | | | |
| Pst/Psh_16 | Elymus repens | Other | 1 | 7 | | | | | | | |
| Pst/Psh_16 | Elymus repens | Other | 2 | 7 | | | | | | | |
| Pst/Psh_16 | Avocet S | Wheat | 1 | | | 5 | 2 | | | | |
| Pst/Psh_16 | Avocet S | Wheat | 2 | | | 3 | 2 | 2 | | | |
| Pst/Psh_16 | Avocet SP | Wheat | 1 | 7 | | | | | | | |
| Pst/Psh_16 | Avocet SP | Wheat | 2 | 7 | | | | | | | |
| Pst/Psh_16 | Avocet Yr1 | Wheat | 1 | 6 | | | | | | | |
| Pst/Psh_16 | Avocet Yr1 | Wheat | 2 | 6 | | | | | | | |
| Pst/Psh_16 | Avocet Yr15 | Wheat | 1 | 8 | | | | | | | |
| Pst/Psh_16 | Avocet Yr15 | Wheat | 2 | 8 | | | | | | | |
| Pst/Psh_16 | Avocet Yr17 | Wheat | 1 | 6 | | | | | | | |
| Pst/Psh_16 | Avocet Yr17 | Wheat | 2 | 6 | | | | | | | |
| Pst/Psh_16 | Avocet Yr24 | Wheat | 1 | 7 | | | | | | | |
| Pst/Psh_16 | Avocet Yr24 | Wheat | 2 | 7 | | | | | | | |

| | | | | | | | | | | | |
|---|---|---|---|---|---|---|---|---|---|---|---|
| Pst/Psh_16 | Avocet Yr27 | Wheat | 1 | 8 | | | | | | | |
| Pst/Psh_16 | Avocet Yr27 | Wheat | 2 | 8 | | | | | | | |
| Pst/Psh_16 | Avocet Yr32 | Wheat | 1 | 7 | | | | | | | |
| Pst/Psh_16 | Avocet Yr32 | Wheat | 2 | | 7 | | | | | | |
| Pst/Psh_16 | Avocet Yr5 | Wheat | 1 | 8 | | | | | | | |
| Pst/Psh_16 | Avocet Yr5 | Wheat | 2 | 8 | | | | | | | |
| Pst/Psh_16 | Avocet Yr6 | Wheat | 1 | | | 8 | | | | | |
| Pst/Psh_16 | Avocet Yr6 | Wheat | 2 | | | 2 | 2 | 4 | | | |
| Pst/Psh_16 | Avocet Yr7 | Wheat | 1 | | 3 | | | | | | |
| Pst/Psh_16 | Avocet Yr7 | Wheat | 2 | | 3 | | | | | | |
| Pst/Psh_16 | Avocet Yr8 | Wheat | 1 | 8 | | | | | | | |
| Pst/Psh_16 | Avocet Yr8 | Wheat | 2 | 4 | 4 | | | | | | |
| Pst/Psh_16 | Avocet Yr9 | Wheat | 1 | | | | 4 | 2 | 2 | | |
| Pst/Psh_16 | Avocet Yr9 | Wheat | 2 | | | | | | 5 | 3 | |
| Pst/Psh_16 | Chinese 166 | Wheat | 1 | 8 | | | | | | | |
| Pst/Psh_16 | Chinese 166 | Wheat | 2 | | 8 | | | | | | |
| Pst/Psh_16 | Compair | Wheat | 1 | 8 | | | | | | | |
| Pst/Psh_16 | Compair | Wheat | 2 | 8 | | | | | | | |
| Pst/Psh_16 | Hybrid 46 | Wheat | 1 | 7 | | | | | | | |
| Pst/Psh_16 | Hybrid 46 | Wheat | 2 | 7 | | | | | | | |
| Pst/Psh_16 | Kalyansona | Wheat | 1 | | 6 | | | | | | |
| Pst/Psh_16 | Kalyansona | Wheat | 2 | | | 6 | | | | | |
| Pst/Psh_16 | Lee | Wheat | 1 | | | 7 | | | | | |
| Pst/Psh_16 | Lee | Wheat | 2 | | | 7 | | | | | |
| Pst/Psh_16 | Moro | Wheat | 1 | 7 | | | | | | | |
| Pst/Psh_16 | Moro | Wheat | 2 | 7 | | | | | | | |
| Pst/Psh_16 | Morocco | Wheat | 1 | | | | | 2 | 3 | 2 | |
| Pst/Psh_16 | Morocco | Wheat | 2 | | | | | | 2 | 3 | 2 |
| Pst/Psh_16 | Sleipner | Wheat | 1 | | 8 | | | | | | |
| Pst/Psh_16 | Sleipner | Wheat | 2 | | 8 | | | | | | |
| Pst/Psh_16 | Suwon/Omar | Wheat | 1 | 7 | | | | | | | |
| Pst/Psh_16 | Suwon/Omar | Wheat | 2 | 7 | | | | | | | |
| Pst/Psh_16 | TP 981 | Wheat | 1 | 4 | 4 | | | | | | |
| Pst/Psh_16 | TP 981 | Wheat | 2 | 4 | 4 | | | | | | |
| Pst/Psh_16 | Triticum dicoccon | Wheat | 1 | 7 | | | | | | | |
| Pst/Psh_16 | Triticum dicoccon | Wheat | 2 | 7 | | | | | | | |
| Pst/Psh_16 | Triticum monococcum | Wheat | 1 | 6 | | | | | | | |
| Pst/Psh_16 | Triticum monococcum | Wheat | 2 | 6 | | | | | | | |
| Pst/Psh_16 | Vilmorin 23 | Wheat | 1 | 7 | | | | | | | |
| Pst/Psh_16 | Vilmorin 23 | Wheat | 2 | 7 | | | | | | | |
| Pst/Psh_16 | VPM1 | Wheat | 1 | 7 | | | | | | | |
| Pst/Psh_16 | VPM1 | Wheat | 2 | 7 | | | | | | | |
| Pst/Psh_10 | Abed Binder | Barley | 1 | 8 | | | | | | | |
| Pst/Psh_10 | Abed Binder | Barley | 2 | 4 | 4 | | | | | | |
| Pst/Psh_10 | Afzal | Barley | 1 | | | | | | | 2 | 5 |
| Pst/Psh_10 | Afzal | Barley | 2 | | | | | | | | 7 |
| Pst/Psh_10 | Astrix | Barley | 1 | 8 | | | | | | | |
| Pst/Psh_10 | Astrix | Barley | 2 | 8 | | | | | | | |
| Pst/Psh_10 | Bancroft | Barley | 1 | 7 | | | | | | | |
| Pst/Psh_10 | Bancroft | Barley | 2 | 7 | | | | | | | |

| | | | | | | | | | | | |
|---|---|---|---|---|---|---|---|---|---|---|---|
| Pst/Psh_10 | Bigo | Barley | 1 | 7 | | | | | | | |
| Pst/Psh_10 | Bigo | Barley | 2 | 7 | | | | | | | |
| Pst/Psh_10 | Emir | Barley | 1 | | | | | | | 1 | 5 |
| Pst/Psh_10 | Emir | Barley | 2 | | | | | | | 1 | 5 |
| Pst/Psh_10 | Heils Franken | Barley | 1 | 7 | | | | | | | |
| Pst/Psh_10 | Heils Franken | Barley | 2 | 7 | | | | | | | |
| Pst/Psh_10 | Hiproly | Barley | 1 | 7 | | | | | | | |
| Pst/Psh_10 | Hiproly | Barley | 2 | 7 | | | | | | | |
| Pst/Psh_10 | Hordeum spontaneum | Barley | 1 | | | | | | 4 | 4 | |
| Pst/Psh_10 | Hordeum spontaneum | Barley | 2 | | | | | | 4 | 4 | |
| Pst/Psh_10 | I5 | Barley | 1 | 4 | 4 | | | | | | |
| Pst/Psh_10 | I5 | Barley | 2 | 4 | | | | | | | |
| Pst/Psh_10 | Trumpf | Barley | 1 | 4 | 4 | | | | | | |
| Pst/Psh_10 | Trumpf | Barley | 2 | 4 | 4 | | | | | | |
| Pst/Psh_10 | Varanda | Barley | 1 | | | | | | 3 | 3 | |
| Pst/Psh_10 | Varanda | Barley | 2 | | | | | | 2 | 4 | |
| Pst/Psh_10 | Aegilops tauschii | Other | 1 | 7 | | | | | | | |
| Pst/Psh_10 | Aegilops tauschii | Other | 2 | 7 | | | | | | | |
| Pst/Psh_10 | Elymus repens | Other | 1 | 7 | | | | | | | |
| Pst/Psh_10 | Elymus repens | Other | 2 | 7 | | | | | | | |
| Pst/Psh_10 | Avocet S | Wheat | 1 | | | 5 | 2 | | | | |
| Pst/Psh_10 | Avocet S | Wheat | 2 | | | | 2 | 2 | 3 | | |
| Pst/Psh_10 | Avocet SP | Wheat | 1 | 7 | | | | | | | |
| Pst/Psh_10 | Avocet SP | Wheat | 2 | 7 | | | | | | | |
| Pst/Psh_10 | Avocet Yr1 | Wheat | 1 | 3 | 3 | | | | | | |
| Pst/Psh_10 | Avocet Yr1 | Wheat | 2 | 3 | 3 | | | | | | |
| Pst/Psh_10 | Avocet Yr15 | Wheat | 1 | 7 | | | | | | | |
| Pst/Psh_10 | Avocet Yr15 | Wheat | 2 | 7 | | | | | | | |
| Pst/Psh_10 | Avocet Yr17 | Wheat | 1 | 6 | | | | | | | |
| Pst/Psh_10 | Avocet Yr17 | Wheat | 2 | 6 | | | | | | | |
| Pst/Psh_10 | Avocet Yr24 | Wheat | 1 | 6 | | | | | | | |
| Pst/Psh_10 | Avocet Yr24 | Wheat | 2 | 6 | | | | | | | |
| Pst/Psh_10 | Avocet Yr27 | Wheat | 1 | 7 | | | | | | | |
| Pst/Psh_10 | Avocet Yr27 | Wheat | 2 | 7 | | | | | | | |
| Pst/Psh_10 | Avocet Yr32 | Wheat | 1 | 6 | | | | | | | |
| Pst/Psh_10 | Avocet Yr32 | Wheat | 2 | | 6 | | | | | | |
| Pst/Psh_10 | Avocet Yr5 | Wheat | 1 | 8 | | | | | | | |
| Pst/Psh_10 | Avocet Yr5 | Wheat | 2 | 8 | | | | | | | |
| Pst/Psh_10 | Avocet Yr6 | Wheat | 1 | 8 | | | | | | | |
| Pst/Psh_10 | Avocet Yr6 | Wheat | 2 | | 4 | 4 | | | | | |
| Pst/Psh_10 | Avocet Yr7 | Wheat | 1 | 4 | | | | | | | |
| Pst/Psh_10 | Avocet Yr7 | Wheat | 2 | | 4 | | | | | | |
| Pst/Psh_10 | Avocet Yr8 | Wheat | 1 | 4 | 4 | | | | | | |
| Pst/Psh_10 | Avocet Yr8 | Wheat | 2 | | 8 | | | | | | |
| Pst/Psh_10 | Avocet Yr9 | Wheat | 1 | | | | 1 | 1 | 2 | 2 | |
| Pst/Psh_10 | Avocet Yr9 | Wheat | 2 | | | | | 1 | 2 | 3 | |
| Pst/Psh_10 | Chinese 166 | Wheat | 1 | 8 | | | | | | | |
| Pst/Psh_10 | Chinese 166 | Wheat | 2 | 4 | 4 | | | | | | |
| Pst/Psh_10 | Compair | Wheat | 1 | 4 | 4 | | | | | | |
| Pst/Psh_10 | Compair | Wheat | 2 | 4 | 4 | | | | | | |

| | | | | | | | | | | | |
|---|---|---|---|---|---|---|---|---|---|---|---|
| Pst/Psh_10 | Hybrid 46 | Wheat | 1 | 8 | | | | | | | |
| Pst/Psh_10 | Hybrid 46 | Wheat | 2 | 8 | | | | | | | |
| Pst/Psh_10 | Kalyansona | Wheat | 1 | | | | | 2 | 2 | 2 | |
| Pst/Psh_10 | Kalyansona | Wheat | 2 | | | | | | 3 | 2 | 1 |
| Pst/Psh_10 | Lee | Wheat | 1 | | | 7 | | | | | |
| Pst/Psh_10 | Lee | Wheat | 2 | | | 7 | | | | | |
| Pst/Psh_10 | Moro | Wheat | 1 | 8 | | | | | | | |
| Pst/Psh_10 | Moro | Wheat | 2 | 8 | | | | | | | |
| Pst/Psh_10 | Morocco | Wheat | 1 | | | | | | | 2 | 5 |
| Pst/Psh_10 | Morocco | Wheat | 2 | | | | | | | | 7 |
| Pst/Psh_10 | Sleipner | Wheat | 1 | 8 | | | | | | | |
| Pst/Psh_10 | Sleipner | Wheat | 2 | | 8 | | | | | | |
| Pst/Psh_10 | Suwon/Omar | Wheat | 1 | 7 | | | | | | | |
| Pst/Psh_10 | Suwon/Omar | Wheat | 2 | 7 | | | | | | | |
| Pst/Psh_10 | TP 981 | Wheat | 1 | 4 | 4 | | | | | | |
| Pst/Psh_10 | TP 981 | Wheat | 2 | 4 | 4 | | | | | | |
| Pst/Psh_10 | Triticum dicoccon | Wheat | 1 | 8 | | | | | | | |
| Pst/Psh_10 | Triticum dicoccon | Wheat | 2 | 8 | | | | | | | |
| Pst/Psh_10 | Triticum monococcum | Wheat | 1 | 8 | | | | | | | |
| Pst/Psh_10 | Triticum monococcum | Wheat | 2 | 8 | | | | | | | |
| Pst/Psh_10 | Vilmorin 23 | Wheat | 1 | 8 | | | | | | | |
| Pst/Psh_10 | Vilmorin 23 | Wheat | 2 | 8 | | | | | | | |
| Pst/Psh_10 | VPM1 | Wheat | 1 | 8 | | | | | | | |
| Pst/Psh_10 | VPM1 | Wheat | 2 | 8 | | | | | | | |
| Pst/Psh_17 | Abed Binder | Barley | 1 | 7 | | | | | | | |
| Pst/Psh_17 | Abed Binder | Barley | 2 | 7 | | | | | | | |
| Pst/Psh_17 | Afzal | Barley | 1 | | | | | | | 2 | 4 |
| Pst/Psh_17 | Afzal | Barley | 2 | | | | | | | | 6 |
| Pst/Psh_17 | Astrix | Barley | 1 | 7 | | | | | | | |
| Pst/Psh_17 | Astrix | Barley | 2 | 7 | | | | | | | |
| Pst/Psh_17 | Bancroft | Barley | 1 | 7 | | | | | | | |
| Pst/Psh_17 | Bancroft | Barley | 2 | 7 | | | | | | | |
| Pst/Psh_17 | Bigo | Barley | 1 | 4 | 4 | | | | | | |
| Pst/Psh_17 | Bigo | Barley | 2 | 4 | 4 | | | | | | |
| Pst/Psh_17 | Emir | Barley | 1 | | | | | | | 3 | 3 |
| Pst/Psh_17 | Emir | Barley | 2 | | | | | | | | 6 |
| Pst/Psh_17 | Heils Franken | Barley | 1 | 8 | | | | | | | |
| Pst/Psh_17 | Heils Franken | Barley | 2 | 4 | 4 | | | | | | |
| Pst/Psh_17 | Hiproly | Barley | 1 | 8 | | | | | | | |
| Pst/Psh_17 | Hiproly | Barley | 2 | 8 | | | | | | | |
| Pst/Psh_17 | Hordeum spontaneum | Barley | 1 | | | | | | | 3 | 3 |
| Pst/Psh_17 | Hordeum spontaneum | Barley | 2 | | | | | | | 1 | 5 |
| Pst/Psh_17 | I5 | Barley | 1 | 7 | | | | | | | |
| Pst/Psh_17 | I5 | Barley | 2 | 7 | | | | | | | |
| Pst/Psh_17 | Trumpf | Barley | 1 | 4 | 4 | | | | | | |
| Pst/Psh_17 | Trumpf | Barley | 2 | 4 | 4 | | | | | | |
| Pst/Psh_17 | Varanda | Barley | 1 | 6 | | | | | | | |
| Pst/Psh_17 | Varanda | Barley | 2 | 6 | | | | | | | |
| Pst/Psh_17 | Aegilops tauschii | Other | 1 | 8 | | | | | | | |
| Pst/Psh_17 | Aegilops tauschii | Other | 2 | 8 | | | | | | | |

| | | | | | | | | | | | |
|---|---|---|---|---|---|---|---|---|---|---|---|
| Pst/Psh_17 | Elymus repens | Other | 1 | 6 | | | | | | | |
| Pst/Psh_17 | Elymus repens | Other | 2 | 6 | | | | | | | |
| Pst/Psh_17 | Avocet S | Wheat | 1 | | | 6 | 1 | 1 | | | |
| Pst/Psh_17 | Avocet S | Wheat | 2 | | | | 2 | 2 | 4 | | |
| Pst/Psh_17 | Avocet SP | Wheat | 1 | 8 | | | | | | | |
| Pst/Psh_17 | Avocet SP | Wheat | 2 | 8 | | | | | | | |
| Pst/Psh_17 | Avocet Yr1 | Wheat | 1 | 6 | | | | | | | |
| Pst/Psh_17 | Avocet Yr1 | Wheat | 2 | | 6 | | | | | | |
| Pst/Psh_17 | Avocet Yr15 | Wheat | 1 | 6 | | | | | | | |
| Pst/Psh_17 | Avocet Yr15 | Wheat | 2 | 6 | | | | | | | |
| Pst/Psh_17 | Avocet Yr17 | Wheat | 1 | 6 | | | | | | | |
| Pst/Psh_17 | Avocet Yr17 | Wheat | 2 | 6 | | | | | | | |
| Pst/Psh_17 | Avocet Yr24 | Wheat | 1 | 8 | | | | | | | |
| Pst/Psh_17 | Avocet Yr24 | Wheat | 2 | 8 | | | | | | | |
| Pst/Psh_17 | Avocet Yr27 | Wheat | 1 | 7 | | | | | | | |
| Pst/Psh_17 | Avocet Yr27 | Wheat | 2 | 7 | | | | | | | |
| Pst/Psh_17 | Avocet Yr32 | Wheat | 1 | 4 | 4 | | | | | | |
| Pst/Psh_17 | Avocet Yr32 | Wheat | 2 | 4 | 4 | | | | | | |
| Pst/Psh_17 | Avocet Yr5 | Wheat | 1 | 7 | | | | | | | |
| Pst/Psh_17 | Avocet Yr5 | Wheat | 2 | 7 | | | | | | | |
| Pst/Psh_17 | Avocet Yr6 | Wheat | 1 | | | 4 | 3 | | | | |
| Pst/Psh_17 | Avocet Yr6 | Wheat | 2 | | | | 2 | 2 | 3 | | |
| Pst/Psh_17 | Avocet Yr7 | Wheat | 1 | 6 | | | | | | | |
| Pst/Psh_17 | Avocet Yr7 | Wheat | 2 | | 6 | | | | | | |
| Pst/Psh_17 | Avocet Yr8 | Wheat | 1 | 8 | | | | | | | |
| Pst/Psh_17 | Avocet Yr8 | Wheat | 2 | | 8 | | | | | | |
| Pst/Psh_17 | Avocet Yr9 | Wheat | 1 | | | | 3 | 2 | 3 | | |
| Pst/Psh_17 | Avocet Yr9 | Wheat | 2 | | | | 1 | 1 | 3 | 3 | |
| Pst/Psh_17 | Chinese 166 | Wheat | 1 | 8 | | | | | | | |
| Pst/Psh_17 | Chinese 166 | Wheat | 2 | 4 | 4 | | | | | | |
| Pst/Psh_17 | Compair | Wheat | 1 | 8 | | | | | | | |
| Pst/Psh_17 | Compair | Wheat | 2 | 8 | | | | | | | |
| Pst/Psh_17 | Hybrid 46 | Wheat | 1 | 8 | | | | | | | |
| Pst/Psh_17 | Hybrid 46 | Wheat | 2 | 3 | 5 | | | | | | |
| Pst/Psh_17 | Kalyansona | Wheat | 1 | | 6 | | | | | | |
| Pst/Psh_17 | Kalyansona | Wheat | 2 | | 3 | 3 | | | | | |
| Pst/Psh_17 | Lee | Wheat | 1 | | | 7 | | | | | |
| Pst/Psh_17 | Lee | Wheat | 2 | | | 7 | | | | | |
| Pst/Psh_17 | Moro | Wheat | 1 | 7 | | | | | | | |
| Pst/Psh_17 | Moro | Wheat | 2 | 7 | | | | | | | |
| Pst/Psh_17 | Morocco | Wheat | 1 | | | | 2 | 2 | 2 | | |
| Pst/Psh_17 | Morocco | Wheat | 2 | | | | | | 2 | 3 | 1 |
| Pst/Psh_17 | Sleipner | Wheat | 1 | 8 | | | | | | | |
| Pst/Psh_17 | Sleipner | Wheat | 2 | | 8 | | | | | | |
| Pst/Psh_17 | Suwon/Omar | Wheat | 1 | 7 | | | | | | | |
| Pst/Psh_17 | Suwon/Omar | Wheat | 2 | 7 | | | | | | | |
| Pst/Psh_17 | TP 981 | Wheat | 1 | 7 | | | | | | | |
| Pst/Psh_17 | TP 981 | Wheat | 2 | 7 | | | | | | | |
| Pst/Psh_17 | Triticum dicoccon | Wheat | 1 | 8 | | | | | | | |
| Pst/Psh_17 | Triticum dicoccon | Wheat | 2 | 8 | | | | | | | |

| | | | | | | | | | | | |
|---|---|---|---|---|---|---|---|---|---|---|---|
| Pst/Psh_17 | Triticum monococcum | Wheat | 1 | 7 | | | | | | | |
| Pst/Psh_17 | Triticum monococcum | Wheat | 2 | 7 | | | | | | | |
| Pst/Psh_17 | Vilmorin 23 | Wheat | 1 | 6 | | | | | | | |
| Pst/Psh_17 | Vilmorin 23 | Wheat | 2 | 6 | | | | | | | |
| Pst/Psh_17 | VPM1 | Wheat | 1 | 6 | | | | | | | |
| Pst/Psh_17 | VPM1 | Wheat | 2 | 6 | | | | | | | |
| Pst/Psh_11 | Abed Binder | Barley | 1 | 7 | | | | | | | |
| Pst/Psh_11 | Abed Binder | Barley | 2 | 7 | | | | | | | |
| Pst/Psh_11 | Afzal | Barley | 1 | | | | | | | 2 | 5 |
| Pst/Psh_11 | Afzal | Barley | 2 | | | | | | | | 7 |
| Pst/Psh_11 | Astrix | Barley | 1 | 8 | | | | | | | |
| Pst/Psh_11 | Astrix | Barley | 2 | 8 | | | | | | | |
| Pst/Psh_11 | Bancroft | Barley | 1 | 6 | | | | | | | |
| Pst/Psh_11 | Bancroft | Barley | 2 | 6 | | | | | | | |
| Pst/Psh_11 | Bigo | Barley | 1 | 7 | | | | | | | |
| Pst/Psh_11 | Bigo | Barley | 2 | 7 | | | | | | | |
| Pst/Psh_11 | Emir | Barley | 1 | | | | | | | 2 | 4 |
| Pst/Psh_11 | Emir | Barley | 2 | | | | | | | 1 | 5 |
| Pst/Psh_11 | Heils Franken | Barley | 1 | 4 | 3 | | | | | | |
| Pst/Psh_11 | Heils Franken | Barley | 2 | | 7 | | | | | | |
| Pst/Psh_11 | Hiproly | Barley | 1 | 8 | | | | | | | |
| Pst/Psh_11 | Hiproly | Barley | 2 | 8 | | | | | | | |
| Pst/Psh_11 | Hordeum spontaneum | Barley | 1 | | | | 2 | 2 | 2 | | |
| Pst/Psh_11 | Hordeum spontaneum | Barley | 2 | | | | | 3 | 3 | | |
| Pst/Psh_11 | I5 | Barley | 1 | 8 | | | | | | | |
| Pst/Psh_11 | I5 | Barley | 2 | 8 | | | | | | | |
| Pst/Psh_11 | Trumpf | Barley | 1 | 6 | | | | | | | |
| Pst/Psh_11 | Trumpf | Barley | 2 | 6 | | | | | | | |
| Pst/Psh_11 | Varanda | Barley | 1 | | | | | | 3 | 3 | |
| Pst/Psh_11 | Varanda | Barley | 2 | | | | | | | 6 | |
| Pst/Psh_11 | Aegilops tauschii | Other | 1 | 7 | | | | | | | |
| Pst/Psh_11 | Aegilops tauschii | Other | 2 | 7 | | | | | | | |
| Pst/Psh_11 | Elymus repens | Other | 1 | 6 | | | | | | | |
| Pst/Psh_11 | Elymus repens | Other | 2 | 6 | | | | | | | |
| Pst/Psh_11 | Avocet S | Wheat | 1 | | 4 | 4 | | | | | |
| Pst/Psh_11 | Avocet S | Wheat | 2 | | | 4 | 2 | 2 | | | |
| Pst/Psh_11 | Avocet SP | Wheat | 1 | 8 | | | | | | | |
| Pst/Psh_11 | Avocet SP | Wheat | 2 | 8 | | | | | | | |
| Pst/Psh_11 | Avocet Yr1 | Wheat | 1 | 7 | | | | | | | |
| Pst/Psh_11 | Avocet Yr1 | Wheat | 2 | | 7 | | | | | | |
| Pst/Psh_11 | Avocet Yr15 | Wheat | 1 | 7 | | | | | | | |
| Pst/Psh_11 | Avocet Yr15 | Wheat | 2 | 7 | | | | | | | |
| Pst/Psh_11 | Avocet Yr17 | Wheat | 1 | 6 | | | | | | | |
| Pst/Psh_11 | Avocet Yr17 | Wheat | 2 | 6 | | | | | | | |
| Pst/Psh_11 | Avocet Yr24 | Wheat | 1 | 7 | | | | | | | |
| Pst/Psh_11 | Avocet Yr24 | Wheat | 2 | 7 | | | | | | | |
| Pst/Psh_11 | Avocet Yr27 | Wheat | 1 | 7 | | | | | | | |
| Pst/Psh_11 | Avocet Yr27 | Wheat | 2 | 7 | | | | | | | |
| Pst/Psh_11 | Avocet Yr32 | Wheat | 1 | 8 | | | | | | | |
| Pst/Psh_11 | Avocet Yr32 | Wheat | 2 | | 8 | | | | | | |

| | | | | | | | | | | | |
|---|---|---|---|---|---|---|---|---|---|---|---|
| Pst/Psh_11 | Avocet Yr5 | Wheat | 1 | 8 | | | | | | | |
| Pst/Psh_11 | Avocet Yr5 | Wheat | 2 | 8 | | | | | | | |
| Pst/Psh_11 | Avocet Yr6 | Wheat | 1 | | 4 | 3 | | | | | |
| Pst/Psh_11 | Avocet Yr6 | Wheat | 2 | | | 3 | 2 | 2 | | | |
| Pst/Psh_11 | Avocet Yr7 | Wheat | 1 | 6 | | | | | | | |
| Pst/Psh_11 | Avocet Yr7 | Wheat | 2 | | 6 | | | | | | |
| Pst/Psh_11 | Avocet Yr8 | Wheat | 1 | 4 | 4 | | | | | | |
| Pst/Psh_11 | Avocet Yr8 | Wheat | 2 | | 4 | 4 | | | | | |
| Pst/Psh_11 | Avocet Yr9 | Wheat | 1 | | | 2 | 3 | 2 | 1 | | |
| Pst/Psh_11 | Avocet Yr9 | Wheat | 2 | | | | 2 | 2 | 2 | 2 | |
| Pst/Psh_11 | Chinese 166 | Wheat | 1 | | | | | | | | 7 |
| Pst/Psh_11 | Chinese 166 | Wheat | 2 | | | | | | | | 7 |
| Pst/Psh_11 | Compair | Wheat | 1 | 3 | 4 | | | | | | |
| Pst/Psh_11 | Compair | Wheat | 2 | 4 | 3 | | | | | | |
| Pst/Psh_11 | Hybrid 46 | Wheat | 1 | 7 | | | | | | | |
| Pst/Psh_11 | Hybrid 46 | Wheat | 2 | 7 | | | | | | | |
| Pst/Psh_11 | Kalyansona | Wheat | 1 | | | 3 | 3 | | | | |
| Pst/Psh_11 | Kalyansona | Wheat | 2 | | | 6 | | | | | |
| Pst/Psh_11 | Lee | Wheat | 1 | | 3 | 3 | | | | | |
| Pst/Psh_11 | Lee | Wheat | 2 | | 3 | 3 | | | | | |
| Pst/Psh_11 | Moro | Wheat | 1 | 8 | | | | | | | |
| Pst/Psh_11 | Moro | Wheat | 2 | 8 | | | | | | | |
| Pst/Psh_11 | Morocco | Wheat | 1 | | | | | | 3 | 3 | |
| Pst/Psh_11 | Morocco | Wheat | 2 | | | | | | 3 | 3 | |
| Pst/Psh_11 | Sleipner | Wheat | 1 | 7 | | | | | | | |
| Pst/Psh_11 | Sleipner | Wheat | 2 | 4 | 3 | | | | | | |
| Pst/Psh_11 | Suwon/Omar | Wheat | 1 | 8 | | | | | | | |
| Pst/Psh_11 | Suwon/Omar | Wheat | 2 | 8 | | | | | | | |
| Pst/Psh_11 | TP 981 | Wheat | 1 | 8 | | | | | | | |
| Pst/Psh_11 | TP 981 | Wheat | 2 | 8 | | | | | | | |
| Pst/Psh_11 | Triticum dicoccon | Wheat | 1 | 7 | | | | | | | |
| Pst/Psh_11 | Triticum dicoccon | Wheat | 2 | 7 | | | | | | | |
| Pst/Psh_11 | Triticum monococcum | Wheat | 1 | 8 | | | | | | | |
| Pst/Psh_11 | Triticum monococcum | Wheat | 2 | 8 | | | | | | | |
| Pst/Psh_11 | Vilmorin 23 | Wheat | 1 | 8 | | | | | | | |
| Pst/Psh_11 | Vilmorin 23 | Wheat | 2 | 8 | | | | | | | |
| Pst/Psh_11 | VPM1 | Wheat | 1 | 8 | | | | | | | |
| Pst/Psh_11 | VPM1 | Wheat | 2 | 8 | | | | | | | |

**Table S6.** Median infection type (IT) values used for heatmap generation. Median IT values were calculated for each isolate × host combination from infection type scores recorded on the first and second leaves at 18 days post-inoculation using the 0–9 scale described in Table S5. When the median fell between two adjacent classes, the midpoint value was retained (e.g. 6.5).

| Isolate | Line | Host Group | Median IT |
|---|---|---|---|
| Psh | Abed Binder | Barley | 0 |
| Psh | Aegilops tauschii | Other | 0 |
| Psh | Afzal | Barley | 7 |
| Psh | Astrix | Barley | 0 |
| Psh | Avocet S | Wheat | 0 |
| Psh | Avocet SP | Wheat | 0 |
| Psh | Avocet Yr1 | Wheat | 0 |
| Psh | Avocet Yr15 | Wheat | 0 |
| Psh | Avocet Yr17 | Wheat | 0 |
| Psh | Avocet Yr24 | Wheat | 0 |
| Psh | Avocet Yr27 | Wheat | 0 |
| Psh | Avocet Yr32 | Wheat | 0 |
| Psh | Avocet Yr5 | Wheat | 0 |
| Psh | Avocet Yr6 | Wheat | 0 |
| Psh | Avocet Yr7 | Wheat | 0 |
| Psh | Avocet Yr8 | Wheat | 0 |
| Psh | Avocet Yr9 | Wheat | 0 |
| Psh | Bancroft | Barley | 0 |
| Psh | Bigo | Barley | 0 |
| Psh | Chinese 166 | Wheat | 0 |
| Psh | Compair | Wheat | 0 |
| Psh | Elymus repens | Other | 0 |
| Psh | Emir | Barley | 7 |
| Psh | Heils Franken | Barley | 0 |
| Psh | Hiproly | Barley | 0 |
| Psh | Hordeum spontaneum | Barley | 7 |
| Psh | Hybrid 46 | Wheat | 0 |
| Psh | I5 | Barley | 0 |
| Psh | Kalyansona | Wheat | 0 |
| Psh | Lee | Wheat | 0 |
| Psh | Moro | Wheat | 0 |
| Psh | Morocco | Wheat | 0 |
| Psh | Sleipner | Wheat | 0 |
| Psh | Suwon/Omar | Wheat | 0 |
| Psh | TP 981 | Wheat | 0 |
| Psh | Triticum dicoccon | Wheat | 0 |
| Psh | Triticum monococcum | Wheat | 0 |
| Psh | Trumpf | Barley | 0 |

| | | | |
|---|---|---|---|
| Psh | VPM1 | Wheat | 0 |
| Psh | Varanda | Barley | 7 |
| Psh | Vilmorin 23 | Wheat | 0 |
| Psh/Pst_1 | Abed Binder | Barley | 0 |
| Psh/Pst_1 | Aegilops tauschii | Other | 0,5 |
| Psh/Pst_1 | Afzal | Barley | 7 |
| Psh/Pst_1 | Astrix | Barley | 0 |
| Psh/Pst_1 | Avocet S | Wheat | 2 |
| Psh/Pst_1 | Avocet SP | Wheat | 0 |
| Psh/Pst_1 | Avocet Yr1 | Wheat | 1 |
| Psh/Pst_1 | Avocet Yr15 | Wheat | 0 |
| Psh/Pst_1 | Avocet Yr17 | Wheat | 0 |
| Psh/Pst_1 | Avocet Yr24 | Wheat | 0 |
| Psh/Pst_1 | Avocet Yr27 | Wheat | 0 |
| Psh/Pst_1 | Avocet Yr32 | Wheat | 0,5 |
| Psh/Pst_1 | Avocet Yr5 | Wheat | 0 |
| Psh/Pst_1 | Avocet Yr6 | Wheat | 0,5 |
| Psh/Pst_1 | Avocet Yr7 | Wheat | 1 |
| Psh/Pst_1 | Avocet Yr8 | Wheat | 1 |
| Psh/Pst_1 | Avocet Yr9 | Wheat | 0,5 |
| Psh/Pst_1 | Bancroft | Barley | 0 |
| Psh/Pst_1 | Bigo | Barley | 0 |
| Psh/Pst_1 | Chinese 166 | Wheat | 7 |
| Psh/Pst_1 | Compair | Wheat | 0 |
| Psh/Pst_1 | Elymus repens | Other | 0 |
| Psh/Pst_1 | Emir | Barley | 7 |
| Psh/Pst_1 | Heils Franken | Barley | 0 |
| Psh/Pst_1 | Hiproly | Barley | 0 |
| Psh/Pst_1 | Hordeum spontaneum | Barley | 6,5 |
| Psh/Pst_1 | Hybrid 46 | Wheat | 0 |
| Psh/Pst_1 | I5 | Barley | 0 |
| Psh/Pst_1 | Kalyansona | Wheat | 0,5 |
| Psh/Pst_1 | Lee | Wheat | 2 |
| Psh/Pst_1 | Moro | Wheat | 0 |
| Psh/Pst_1 | Morocco | Wheat | 2 |
| Psh/Pst_1 | Sleipner | Wheat | 0,5 |
| Psh/Pst_1 | Suwon/Omar | Wheat | 0 |
| Psh/Pst_1 | TP 981 | Wheat | 0 |
| Psh/Pst_1 | Triticum dicoccon | Wheat | 0 |
| Psh/Pst_1 | Triticum monococcum | Wheat | 0 |
| Psh/Pst_1 | Trumpf | Barley | 0 |
| Psh/Pst_1 | VPM1 | Wheat | 0 |
| Psh/Pst_1 | Varanda | Barley | 6 |
| Psh/Pst_1 | Vilmorin 23 | Wheat | 0 |

| | | | |
|---|---|---|---|
| Psh/Pst_13 | Abed Binder | Barley | 0 |
| Psh/Pst_13 | Aegilops tauschii | Other | 0 |
| Psh/Pst_13 | Afzal | Barley | 7 |
| Psh/Pst_13 | Astrix | Barley | 0 |
| Psh/Pst_13 | Avocet S | Wheat | 3 |
| Psh/Pst_13 | Avocet SP | Wheat | 0 |
| Psh/Pst_13 | Avocet Yr1 | Wheat | 0 |
| Psh/Pst_13 | Avocet Yr15 | Wheat | 0 |
| Psh/Pst_13 | Avocet Yr17 | Wheat | 0 |
| Psh/Pst_13 | Avocet Yr24 | Wheat | 0 |
| Psh/Pst_13 | Avocet Yr27 | Wheat | 0 |
| Psh/Pst_13 | Avocet Yr32 | Wheat | 0,5 |
| Psh/Pst_13 | Avocet Yr5 | Wheat | 0 |
| Psh/Pst_13 | Avocet Yr6 | Wheat | 0,5 |
| Psh/Pst_13 | Avocet Yr7 | Wheat | 0,5 |
| Psh/Pst_13 | Avocet Yr8 | Wheat | 1 |
| Psh/Pst_13 | Avocet Yr9 | Wheat | 4 |
| Psh/Pst_13 | Bancroft | Barley | 0 |
| Psh/Pst_13 | Bigo | Barley | 0 |
| Psh/Pst_13 | Chinese 166 | Wheat | 0 |
| Psh/Pst_13 | Compair | Wheat | 0 |
| Psh/Pst_13 | Elymus repens | Other | 0 |
| Psh/Pst_13 | Emir | Barley | 7 |
| Psh/Pst_13 | Heils Franken | Barley | 0,5 |
| Psh/Pst_13 | Hiproly | Barley | 0 |
| Psh/Pst_13 | Hordeum spontaneum | Barley | 6,5 |
| Psh/Pst_13 | Hybrid 46 | Wheat | 0 |
| Psh/Pst_13 | I5 | Barley | 0 |
| Psh/Pst_13 | Kalyansona | Wheat | 3 |
| Psh/Pst_13 | Lee | Wheat | 1,5 |
| Psh/Pst_13 | Moro | Wheat | 0 |
| Psh/Pst_13 | Morocco | Wheat | 4 |
| Psh/Pst_13 | Sleipner | Wheat | 0,5 |
| Psh/Pst_13 | Suwon/Omar | Wheat | 0 |
| Psh/Pst_13 | TP 981 | Wheat | 0 |
| Psh/Pst_13 | Triticum dicoccon | Wheat | 0 |
| Psh/Pst_13 | Triticum monococcum | Wheat | 0 |
| Psh/Pst_13 | Trumpf | Barley | 0 |
| Psh/Pst_13 | VPM1 | Wheat | 0 |
| Psh/Pst_13 | Varanda | Barley | 6,5 |
| Psh/Pst_13 | Vilmorin 23 | Wheat | 0 |
| Psh/Pst_2 | Abed Binder | Barley | 0 |
| Psh/Pst_2 | Aegilops tauschii | Other | 0,5 |
| Psh/Pst_2 | Afzal | Barley | 7 |

| | | | |
|---|---|---|---|
| Psh/Pst_2 | Astrix | Barley | 0 |
| Psh/Pst_2 | Avocet S | Wheat | 2,5 |
| Psh/Pst_2 | Avocet SP | Wheat | 0 |
| Psh/Pst_2 | Avocet Yr1 | Wheat | 1,5 |
| Psh/Pst_2 | Avocet Yr15 | Wheat | 0 |
| Psh/Pst_2 | Avocet Yr17 | Wheat | 0 |
| Psh/Pst_2 | Avocet Yr24 | Wheat | 0 |
| Psh/Pst_2 | Avocet Yr27 | Wheat | 0 |
| Psh/Pst_2 | Avocet Yr32 | Wheat | 1,5 |
| Psh/Pst_2 | Avocet Yr5 | Wheat | 0 |
| Psh/Pst_2 | Avocet Yr6 | Wheat | 1,5 |
| Psh/Pst_2 | Avocet Yr7 | Wheat | 0 |
| Psh/Pst_2 | Avocet Yr8 | Wheat | 1,5 |
| Psh/Pst_2 | Avocet Yr9 | Wheat | 1 |
| Psh/Pst_2 | Bancroft | Barley | 0 |
| Psh/Pst_2 | Bigo | Barley | 0 |
| Psh/Pst_2 | Chinese 166 | Wheat | 7 |
| Psh/Pst_2 | Compair | Wheat | 0 |
| Psh/Pst_2 | Elymus repens | Other | 0 |
| Psh/Pst_2 | Emir | Barley | 7 |
| Psh/Pst_2 | Heils Franken | Barley | 0 |
| Psh/Pst_2 | Hiproly | Barley | 0 |
| Psh/Pst_2 | Hordeum spontaneum | Barley | 6,5 |
| Psh/Pst_2 | Hybrid 46 | Wheat | 0 |
| Psh/Pst_2 | I5 | Barley | 0 |
| Psh/Pst_2 | Kalyansona | Wheat | 2 |
| Psh/Pst_2 | Lee | Wheat | 1 |
| Psh/Pst_2 | Moro | Wheat | 0 |
| Psh/Pst_2 | Morocco | Wheat | 2 |
| Psh/Pst_2 | Sleipner | Wheat | 0,5 |
| Psh/Pst_2 | Suwon/Omar | Wheat | 0 |
| Psh/Pst_2 | TP 981 | Wheat | 0 |
| Psh/Pst_2 | Triticum dicoccon | Wheat | 0 |
| Psh/Pst_2 | Triticum monococcum | Wheat | 0 |
| Psh/Pst_2 | Trumpf | Barley | 0 |
| Psh/Pst_2 | VPM1 | Wheat | 0 |
| Psh/Pst_2 | Varanda | Barley | 6 |
| Psh/Pst_2 | Vilmorin 23 | Wheat | 0 |
| Psh/Pst_3 | Abed Binder | Barley | 0 |
| Psh/Pst_3 | Aegilops tauschii | Other | 0,5 |
| Psh/Pst_3 | Afzal | Barley | 7 |
| Psh/Pst_3 | Astrix | Barley | 0 |
| Psh/Pst_3 | Avocet S | Wheat | 2 |
| Psh/Pst_3 | Avocet SP | Wheat | 0 |

| | | | |
|---|---|---|---|
| Psh/Pst_3 | Avocet Yr1 | Wheat | 1,5 |
| Psh/Pst_3 | Avocet Yr15 | Wheat | 0 |
| Psh/Pst_3 | Avocet Yr17 | Wheat | 0 |
| Psh/Pst_3 | Avocet Yr24 | Wheat | 0 |
| Psh/Pst_3 | Avocet Yr27 | Wheat | 0 |
| Psh/Pst_3 | Avocet Yr32 | Wheat | 0,5 |
| Psh/Pst_3 | Avocet Yr5 | Wheat | 0 |
| Psh/Pst_3 | Avocet Yr6 | Wheat | 1,5 |
| Psh/Pst_3 | Avocet Yr7 | Wheat | 1 |
| Psh/Pst_3 | Avocet Yr8 | Wheat | 2 |
| Psh/Pst_3 | Avocet Yr9 | Wheat | 1 |
| Psh/Pst_3 | Bancroft | Barley | 0 |
| Psh/Pst_3 | Bigo | Barley | 0 |
| Psh/Pst_3 | Chinese 166 | Wheat | 7 |
| Psh/Pst_3 | Compair | Wheat | 0 |
| Psh/Pst_3 | Elymus repens | Other | 0 |
| Psh/Pst_3 | Emir | Barley | 7 |
| Psh/Pst_3 | Heils Franken | Barley | 0 |
| Psh/Pst_3 | Hiproly | Barley | 0 |
| Psh/Pst_3 | Hordeum spontaneum | Barley | 7 |
| Psh/Pst_3 | Hybrid 46 | Wheat | 0 |
| Psh/Pst_3 | I5 | Barley | 0 |
| Psh/Pst_3 | Kalyansona | Wheat | 1 |
| Psh/Pst_3 | Lee | Wheat | 1,5 |
| Psh/Pst_3 | Moro | Wheat | 0 |
| Psh/Pst_3 | Morocco | Wheat | 2 |
| Psh/Pst_3 | Sleipner | Wheat | 0 |
| Psh/Pst_3 | Suwon/Omar | Wheat | 0 |
| Psh/Pst_3 | TP 981 | Wheat | 0 |
| Psh/Pst_3 | Triticum dicoccon | Wheat | 0 |
| Psh/Pst_3 | Triticum monococcum | Wheat | 0 |
| Psh/Pst_3 | Trumpf | Barley | 0 |
| Psh/Pst_3 | VPM1 | Wheat | 0 |
| Psh/Pst_3 | Varanda | Barley | 6,5 |
| Psh/Pst_3 | Vilmorin 23 | Wheat | 0 |
| Psh/Pst_4 | Abed Binder | Barley | 0 |
| Psh/Pst_4 | Aegilops tauschii | Other | 0 |
| Psh/Pst_4 | Afzal | Barley | 7 |
| Psh/Pst_4 | Astrix | Barley | 0 |
| Psh/Pst_4 | Avocet S | Wheat | 3,5 |
| Psh/Pst_4 | Avocet SP | Wheat | 0 |
| Psh/Pst_4 | Avocet Yr1 | Wheat | 0 |
| Psh/Pst_4 | Avocet Yr15 | Wheat | 0 |
| Psh/Pst_4 | Avocet Yr17 | Wheat | 0 |

| | | | |
|---|---|---|---|
| Psh/Pst_4 | Avocet Yr24 | Wheat | 0 |
| Psh/Pst_4 | Avocet Yr27 | Wheat | 0 |
| Psh/Pst_4 | Avocet Yr32 | Wheat | 1 |
| Psh/Pst_4 | Avocet Yr5 | Wheat | 0 |
| Psh/Pst_4 | Avocet Yr6 | Wheat | 1,5 |
| Psh/Pst_4 | Avocet Yr7 | Wheat | 0 |
| Psh/Pst_4 | Avocet Yr8 | Wheat | 0 |
| Psh/Pst_4 | Avocet Yr9 | Wheat | 2,5 |
| Psh/Pst_4 | Bancroft | Barley | 0 |
| Psh/Pst_4 | Bigo | Barley | 0 |
| Psh/Pst_4 | Chinese 166 | Wheat | 0 |
| Psh/Pst_4 | Compair | Wheat | 0 |
| Psh/Pst_4 | Elymus repens | Other | 0 |
| Psh/Pst_4 | Emir | Barley | 7 |
| Psh/Pst_4 | Heils Franken | Barley | 0 |
| Psh/Pst_4 | Hiproly | Barley | 0,5 |
| Psh/Pst_4 | Hordeum spontaneum | Barley | 5,5 |
| Psh/Pst_4 | Hybrid 46 | Wheat | 0 |
| Psh/Pst_4 | I5 | Barley | 0,5 |
| Psh/Pst_4 | Kalyansona | Wheat | 5 |
| Psh/Pst_4 | Lee | Wheat | 2 |
| Psh/Pst_4 | Moro | Wheat | 0 |
| Psh/Pst_4 | Morocco | Wheat | 7 |
| Psh/Pst_4 | Sleipner | Wheat | 0,5 |
| Psh/Pst_4 | Suwon/Omar | Wheat | 0 |
| Psh/Pst_4 | TP 981 | Wheat | 0 |
| Psh/Pst_4 | Triticum dicoccon | Wheat | 0 |
| Psh/Pst_4 | Triticum monococcum | Wheat | 0 |
| Psh/Pst_4 | Trumpf | Barley | 0,5 |
| Psh/Pst_4 | VPM1 | Wheat | 0 |
| Psh/Pst_4 | Varanda | Barley | 6 |
| Psh/Pst_4 | Vilmorin 23 | Wheat | 0 |
| Pst | Abed Binder | Barley | 0 |
| Pst | Aegilops tauschii | Other | 2 |
| Pst | Afzal | Barley | 0 |
| Pst | Astrix | Barley | 0 |
| Pst | Avocet S | Wheat | 7 |
| Pst | Avocet SP | Wheat | 0 |
| Pst | Avocet Yr1 | Wheat | 7 |
| Pst | Avocet Yr15 | Wheat | 0 |
| Pst | Avocet Yr17 | Wheat | 7 |
| Pst | Avocet Yr24 | Wheat | 0 |
| Pst | Avocet Yr27 | Wheat | 4,5 |
| Pst | Avocet Yr32 | Wheat | 7 |

| | | | |
|---|---|---|---|
| Pst | Avocet Yr5 | Wheat | 0 |
| Pst | Avocet Yr6 | Wheat | 7 |
| Pst | Avocet Yr7 | Wheat | 7 |
| Pst | Avocet Yr8 | Wheat | 7 |
| Pst | Avocet Yr9 | Wheat | 7 |
| Pst | Bancroft | Barley | 0 |
| Pst | Bigo | Barley | 0 |
| Pst | Chinese 166 | Wheat | 7 |
| Pst | Compair | Wheat | 2,5 |
| Pst | Elymus repens | Other | 0 |
| Pst | Emir | Barley | 0 |
| Pst | Heils Franken | Barley | 0 |
| Pst | Hiproly | Barley | 0 |
| Pst | Hordeum spontaneum | Barley | 0 |
| Pst | Hybrid 46 | Wheat | 0 |
| Pst | I5 | Barley | 0 |
| Pst | Kalyansona | Wheat | 7 |
| Pst | Lee | Wheat | 7 |
| Pst | Moro | Wheat | 0 |
| Pst | Morocco | Wheat | 7 |
| Pst | Sleipner | Wheat | 7 |
| Pst | Suwon/Omar | Wheat | 0,5 |
| Pst | TP 981 | Wheat | 7 |
| Pst | Triticum dicoccon | Wheat | 0 |
| Pst | Triticum monococcum | Wheat | 7 |
| Pst | Trumpf | Barley | 0 |
| Pst | VPM1 | Wheat | 7 |
| Pst | Varanda | Barley | 0 |
| Pst | Vilmorin 23 | Wheat | 7 |
| Pst/Psh_10 | Abed Binder | Barley | 0 |
| Pst/Psh_10 | Aegilops tauschii | Other | 0 |
| Pst/Psh_10 | Afzal | Barley | 7 |
| Pst/Psh_10 | Astrix | Barley | 0 |
| Pst/Psh_10 | Avocet S | Wheat | 3 |
| Pst/Psh_10 | Avocet SP | Wheat | 0 |
| Pst/Psh_10 | Avocet Yr1 | Wheat | 0,5 |
| Pst/Psh_10 | Avocet Yr15 | Wheat | 0 |
| Pst/Psh_10 | Avocet Yr17 | Wheat | 0 |
| Pst/Psh_10 | Avocet Yr24 | Wheat | 0 |
| Pst/Psh_10 | Avocet Yr27 | Wheat | 0 |
| Pst/Psh_10 | Avocet Yr32 | Wheat | 0,5 |
| Pst/Psh_10 | Avocet Yr5 | Wheat | 0 |
| Pst/Psh_10 | Avocet Yr6 | Wheat | 0,5 |
| Pst/Psh_10 | Avocet Yr7 | Wheat | 0,5 |

| | | | |
|---|---|---|---|
| Pst/Psh_10 | Avocet Yr8 | Wheat | 1 |
| Pst/Psh_10 | Avocet Yr9 | Wheat | 5 |
| Pst/Psh_10 | Bancroft | Barley | 0 |
| Pst/Psh_10 | Bigo | Barley | 0 |
| Pst/Psh_10 | Chinese 166 | Wheat | 0 |
| Pst/Psh_10 | Compair | Wheat | 0,5 |
| Pst/Psh_10 | Elymus repens | Other | 0 |
| Pst/Psh_10 | Emir | Barley | 7 |
| Pst/Psh_10 | Heils Franken | Barley | 0 |
| Pst/Psh_10 | Hiproly | Barley | 0 |
| Pst/Psh_10 | Hordeum spontaneum | Barley | 5,5 |
| Pst/Psh_10 | Hybrid 46 | Wheat | 0 |
| Pst/Psh_10 | I5 | Barley | 0 |
| Pst/Psh_10 | Kalyansona | Wheat | 5 |
| Pst/Psh_10 | Lee | Wheat | 2 |
| Pst/Psh_10 | Moro | Wheat | 0 |
| Pst/Psh_10 | Morocco | Wheat | 7 |
| Pst/Psh_10 | Sleipner | Wheat | 0,5 |
| Pst/Psh_10 | Suwon/Omar | Wheat | 0 |
| Pst/Psh_10 | TP 981 | Wheat | 0,5 |
| Pst/Psh_10 | Triticum dicoccon | Wheat | 0 |
| Pst/Psh_10 | Triticum monococcum | Wheat | 0 |
| Pst/Psh_10 | Trumpf | Barley | 0,5 |
| Pst/Psh_10 | VPM1 | Wheat | 0 |
| Pst/Psh_10 | Varanda | Barley | 6 |
| Pst/Psh_10 | Vilmorin 23 | Wheat | 0 |
| Pst/Psh_11 | Abed Binder | Barley | 0 |
| Pst/Psh_11 | Aegilops tauschii | Other | 0 |
| Pst/Psh_11 | Afzal | Barley | 7 |
| Pst/Psh_11 | Astrix | Barley | 0 |
| Pst/Psh_11 | Avocet S | Wheat | 2 |
| Pst/Psh_11 | Avocet SP | Wheat | 0 |
| Pst/Psh_11 | Avocet Yr1 | Wheat | 0,5 |
| Pst/Psh_11 | Avocet Yr15 | Wheat | 0 |
| Pst/Psh_11 | Avocet Yr17 | Wheat | 0 |
| Pst/Psh_11 | Avocet Yr24 | Wheat | 0 |
| Pst/Psh_11 | Avocet Yr27 | Wheat | 0 |
| Pst/Psh_11 | Avocet Yr32 | Wheat | 0,5 |
| Pst/Psh_11 | Avocet Yr5 | Wheat | 0 |
| Pst/Psh_11 | Avocet Yr6 | Wheat | 2 |
| Pst/Psh_11 | Avocet Yr7 | Wheat | 0,5 |
| Pst/Psh_11 | Avocet Yr8 | Wheat | 1 |
| Pst/Psh_11 | Avocet Yr9 | Wheat | 4 |
| Pst/Psh_11 | Bancroft | Barley | 0 |

| | | | |
|---|---|---|---|
| Pst/Psh_11 | Bigo | Barley | 0 |
| Pst/Psh_11 | Chinese 166 | Wheat | 7 |
| Pst/Psh_11 | Compair | Wheat | 0,5 |
| Pst/Psh_11 | Elymus repens | Other | 0 |
| Pst/Psh_11 | Emir | Barley | 7 |
| Pst/Psh_11 | Heils Franken | Barley | 1 |
| Pst/Psh_11 | Hiproly | Barley | 0 |
| Pst/Psh_11 | Hordeum spontaneum | Barley | 4 |
| Pst/Psh_11 | Hybrid 46 | Wheat | 0 |
| Pst/Psh_11 | I5 | Barley | 0 |
| Pst/Psh_11 | Kalyansona | Wheat | 2 |
| Pst/Psh_11 | Lee | Wheat | 1,5 |
| Pst/Psh_11 | Moro | Wheat | 0 |
| Pst/Psh_11 | Morocco | Wheat | 5,5 |
| Pst/Psh_11 | Sleipner | Wheat | 0 |
| Pst/Psh_11 | Suwon/Omar | Wheat | 0 |
| Pst/Psh_11 | TP 981 | Wheat | 0 |
| Pst/Psh_11 | Triticum dicoccon | Wheat | 0 |
| Pst/Psh_11 | Triticum monococcum | Wheat | 0 |
| Pst/Psh_11 | Trumpf | Barley | 0 |
| Pst/Psh_11 | VPM1 | Wheat | 0 |
| Pst/Psh_11 | Varanda | Barley | 6 |
| Pst/Psh_11 | Vilmorin 23 | Wheat | 0 |
| Pst/Psh_12 | Abed Binder | Barley | 0 |
| Pst/Psh_12 | Aegilops tauschii | Other | 0 |
| Pst/Psh_12 | Afzal | Barley | 7 |
| Pst/Psh_12 | Astrix | Barley | 0 |
| Pst/Psh_12 | Avocet S | Wheat | 1,5 |
| Pst/Psh_12 | Avocet SP | Wheat | 0 |
| Pst/Psh_12 | Avocet Yr1 | Wheat | 0 |
| Pst/Psh_12 | Avocet Yr15 | Wheat | 0 |
| Pst/Psh_12 | Avocet Yr17 | Wheat | 0 |
| Pst/Psh_12 | Avocet Yr24 | Wheat | 0 |
| Pst/Psh_12 | Avocet Yr27 | Wheat | 0 |
| Pst/Psh_12 | Avocet Yr32 | Wheat | 0,5 |
| Pst/Psh_12 | Avocet Yr5 | Wheat | 0 |
| Pst/Psh_12 | Avocet Yr6 | Wheat | 1 |
| Pst/Psh_12 | Avocet Yr7 | Wheat | 1 |
| Pst/Psh_12 | Avocet Yr8 | Wheat | 1 |
| Pst/Psh_12 | Avocet Yr9 | Wheat | 4 |
| Pst/Psh_12 | Bancroft | Barley | 0 |
| Pst/Psh_12 | Bigo | Barley | 0 |
| Pst/Psh_12 | Chinese 166 | Wheat | 0,5 |
| Pst/Psh_12 | Compair | Wheat | 0 |

| | | | |
|---|---|---|---|
| Pst/Psh_12 | Elymus repens | Other | 0 |
| Pst/Psh_12 | Emir | Barley | 7 |
| Pst/Psh_12 | Heils Franken | Barley | 1 |
| Pst/Psh_12 | Hiproly | Barley | 0 |
| Pst/Psh_12 | Hordeum spontaneum | Barley | 6,5 |
| Pst/Psh_12 | Hybrid 46 | Wheat | 0 |
| Pst/Psh_12 | I5 | Barley | 0,5 |
| Pst/Psh_12 | Kalyansona | Wheat | 3,5 |
| Pst/Psh_12 | Lee | Wheat | 2 |
| Pst/Psh_12 | Moro | Wheat | 0 |
| Pst/Psh_12 | Morocco | Wheat | 7 |
| Pst/Psh_12 | Sleipner | Wheat | 1 |
| Pst/Psh_12 | Suwon/Omar | Wheat | 0 |
| Pst/Psh_12 | TP 981 | Wheat | 0 |
| Pst/Psh_12 | Triticum dicoccon | Wheat | 0 |
| Pst/Psh_12 | Triticum monococcum | Wheat | 0 |
| Pst/Psh_12 | Trumpf | Barley | 0 |
| Pst/Psh_12 | VPM1 | Wheat | 0 |
| Pst/Psh_12 | Varanda | Barley | 0 |
| Pst/Psh_12 | Vilmorin 23 | Wheat | 0 |
| Pst/Psh_14 | Abed Binder | Barley | 0 |
| Pst/Psh_14 | Aegilops tauschii | Other | 0 |
| Pst/Psh_14 | Afzal | Barley | 7 |
| Pst/Psh_14 | Astrix | Barley | 0 |
| Pst/Psh_14 | Avocet S | Wheat | 2 |
| Pst/Psh_14 | Avocet SP | Wheat | 0 |
| Pst/Psh_14 | Avocet Yr1 | Wheat | 0,5 |
| Pst/Psh_14 | Avocet Yr15 | Wheat | 0 |
| Pst/Psh_14 | Avocet Yr17 | Wheat | 0 |
| Pst/Psh_14 | Avocet Yr24 | Wheat | 0 |
| Pst/Psh_14 | Avocet Yr27 | Wheat | 0 |
| Pst/Psh_14 | Avocet Yr32 | Wheat | 0 |
| Pst/Psh_14 | Avocet Yr5 | Wheat | 0 |
| Pst/Psh_14 | Avocet Yr6 | Wheat | 2 |
| Pst/Psh_14 | Avocet Yr7 | Wheat | 0 |
| Pst/Psh_14 | Avocet Yr8 | Wheat | 2 |
| Pst/Psh_14 | Avocet Yr9 | Wheat | 4 |
| Pst/Psh_14 | Bancroft | Barley | 0 |
| Pst/Psh_14 | Bigo | Barley | 0 |
| Pst/Psh_14 | Chinese 166 | Wheat | 0 |
| Pst/Psh_14 | Compair | Wheat | 0 |
| Pst/Psh_14 | Elymus repens | Other | 0 |
| Pst/Psh_14 | Emir | Barley | 7 |
| Pst/Psh_14 | Heils Franken | Barley | 0 |

| | | | |
|---|---|---|---|
| Pst/Psh_14 | Hiproly | Barley | 0 |
| Pst/Psh_14 | Hordeum spontaneum | Barley | 6 |
| Pst/Psh_14 | Hybrid 46 | Wheat | 0 |
| Pst/Psh_14 | I5 | Barley | 0 |
| Pst/Psh_14 | Kalyansona | Wheat | 6 |
| Pst/Psh_14 | Lee | Wheat | 2 |
| Pst/Psh_14 | Moro | Wheat | 0 |
| Pst/Psh_14 | Morocco | Wheat | 7 |
| Pst/Psh_14 | Sleipner | Wheat | 0 |
| Pst/Psh_14 | Suwon/Omar | Wheat | 0 |
| Pst/Psh_14 | TP 981 | Wheat | 0 |
| Pst/Psh_14 | Triticum dicoccon | Wheat | 0 |
| Pst/Psh_14 | Triticum monococcum | Wheat | 0 |
| Pst/Psh_14 | Trumpf | Barley | 0 |
| Pst/Psh_14 | VPM1 | Wheat | 0,5 |
| Pst/Psh_14 | Varanda | Barley | 0 |
| Pst/Psh_14 | Vilmorin 23 | Wheat | 0 |
| Pst/Psh_15 | Abed Binder | Barley | 0,5 |
| Pst/Psh_15 | Aegilops tauschii | Other | 0 |
| Pst/Psh_15 | Afzal | Barley | 7 |
| Pst/Psh_15 | Astrix | Barley | 0 |
| Pst/Psh_15 | Avocet S | Wheat | 1 |
| Pst/Psh_15 | Avocet SP | Wheat | 0 |
| Pst/Psh_15 | Avocet Yr1 | Wheat | 0 |
| Pst/Psh_15 | Avocet Yr15 | Wheat | 0 |
| Pst/Psh_15 | Avocet Yr17 | Wheat | 0 |
| Pst/Psh_15 | Avocet Yr24 | Wheat | 0 |
| Pst/Psh_15 | Avocet Yr27 | Wheat | 0 |
| Pst/Psh_15 | Avocet Yr32 | Wheat | 0 |
| Pst/Psh_15 | Avocet Yr5 | Wheat | 0 |
| Pst/Psh_15 | Avocet Yr6 | Wheat | 1 |
| Pst/Psh_15 | Avocet Yr7 | Wheat | 0 |
| Pst/Psh_15 | Avocet Yr8 | Wheat | 0 |
| Pst/Psh_15 | Avocet Yr9 | Wheat | 3 |
| Pst/Psh_15 | Bancroft | Barley | 0 |
| Pst/Psh_15 | Bigo | Barley | 0 |
| Pst/Psh_15 | Chinese 166 | Wheat | 0,5 |
| Pst/Psh_15 | Compair | Wheat | 0 |
| Pst/Psh_15 | Elymus repens | Other | 0 |
| Pst/Psh_15 | Emir | Barley | 7 |
| Pst/Psh_15 | Heils Franken | Barley | 0 |
| Pst/Psh_15 | Hiproly | Barley | 0 |
| Pst/Psh_15 | Hordeum spontaneum | Barley | 6 |
| Pst/Psh_15 | Hybrid 46 | Wheat | 0 |

| | | | |
|---|---|---|---|
| Pst/Psh_15 | I5 | Barley | 0 |
| Pst/Psh_15 | Kalyansona | Wheat | 7 |
| Pst/Psh_15 | Lee | Wheat | 2 |
| Pst/Psh_15 | Moro | Wheat | 0 |
| Pst/Psh_15 | Morocco | Wheat | 7 |
| Pst/Psh_15 | Sleipner | Wheat | 0 |
| Pst/Psh_15 | Suwon/Omar | Wheat | 0 |
| Pst/Psh_15 | TP 981 | Wheat | 0 |
| Pst/Psh_15 | Triticum dicoccon | Wheat | 0 |
| Pst/Psh_15 | Triticum monococcum | Wheat | 0 |
| Pst/Psh_15 | Trumpf | Barley | 0 |
| Pst/Psh_15 | VPM1 | Wheat | 0 |
| Pst/Psh_15 | Varanda | Barley | 2,5 |
| Pst/Psh_15 | Vilmorin 23 | Wheat | 0 |
| Pst/Psh_16 | Abed Binder | Barley | 0 |
| Pst/Psh_16 | Aegilops tauschii | Other | 0 |
| Pst/Psh_16 | Afzal | Barley | 7 |
| Pst/Psh_16 | Astrix | Barley | 0,5 |
| Pst/Psh_16 | Avocet S | Wheat | 2 |
| Pst/Psh_16 | Avocet SP | Wheat | 0 |
| Pst/Psh_16 | Avocet Yr1 | Wheat | 0 |
| Pst/Psh_16 | Avocet Yr15 | Wheat | 0 |
| Pst/Psh_16 | Avocet Yr17 | Wheat | 0 |
| Pst/Psh_16 | Avocet Yr24 | Wheat | 0 |
| Pst/Psh_16 | Avocet Yr27 | Wheat | 0 |
| Pst/Psh_16 | Avocet Yr32 | Wheat | 0,5 |
| Pst/Psh_16 | Avocet Yr5 | Wheat | 0 |
| Pst/Psh_16 | Avocet Yr6 | Wheat | 2 |
| Pst/Psh_16 | Avocet Yr7 | Wheat | 1 |
| Pst/Psh_16 | Avocet Yr8 | Wheat | 0 |
| Pst/Psh_16 | Avocet Yr9 | Wheat | 5 |
| Pst/Psh_16 | Bancroft | Barley | 0 |
| Pst/Psh_16 | Bigo | Barley | 0 |
| Pst/Psh_16 | Chinese 166 | Wheat | 0,5 |
| Pst/Psh_16 | Compair | Wheat | 0 |
| Pst/Psh_16 | Elymus repens | Other | 0 |
| Pst/Psh_16 | Emir | Barley | 7 |
| Pst/Psh_16 | Heils Franken | Barley | 0 |
| Pst/Psh_16 | Hiproly | Barley | 0 |
| Pst/Psh_16 | Hordeum spontaneum | Barley | 6 |
| Pst/Psh_16 | Hybrid 46 | Wheat | 0 |
| Pst/Psh_16 | I5 | Barley | 0 |
| Pst/Psh_16 | Kalyansona | Wheat | 1,5 |
| Pst/Psh_16 | Lee | Wheat | 2 |

| | | | |
|---|---|---|---|
| Pst/Psh_16 | Moro | Wheat | 0 |
| Pst/Psh_16 | Morocco | Wheat | 5,5 |
| Pst/Psh_16 | Sleipner | Wheat | 1 |
| Pst/Psh_16 | Suwon/Omar | Wheat | 0 |
| Pst/Psh_16 | TP 981 | Wheat | 0,5 |
| Pst/Psh_16 | Triticum dicoccon | Wheat | 0 |
| Pst/Psh_16 | Triticum monococcum | Wheat | 0 |
| Pst/Psh_16 | Trumpf | Barley | 0 |
| Pst/Psh_16 | VPM1 | Wheat | 0 |
| Pst/Psh_16 | Varanda | Barley | 3 |
| Pst/Psh_16 | Vilmorin 23 | Wheat | 0 |
| Pst/Psh_17 | Abed Binder | Barley | 0 |
| Pst/Psh_17 | Aegilops tauschii | Other | 0 |
| Pst/Psh_17 | Afzal | Barley | 7 |
| Pst/Psh_17 | Astrix | Barley | 0 |
| Pst/Psh_17 | Avocet S | Wheat | 3 |
| Pst/Psh_17 | Avocet SP | Wheat | 0 |
| Pst/Psh_17 | Avocet Yr1 | Wheat | 0,5 |
| Pst/Psh_17 | Avocet Yr15 | Wheat | 0 |
| Pst/Psh_17 | Avocet Yr17 | Wheat | 0 |
| Pst/Psh_17 | Avocet Yr24 | Wheat | 0 |
| Pst/Psh_17 | Avocet Yr27 | Wheat | 0 |
| Pst/Psh_17 | Avocet Yr32 | Wheat | 0,5 |
| Pst/Psh_17 | Avocet Yr5 | Wheat | 0 |
| Pst/Psh_17 | Avocet Yr6 | Wheat | 3 |
| Pst/Psh_17 | Avocet Yr7 | Wheat | 0,5 |
| Pst/Psh_17 | Avocet Yr8 | Wheat | 0,5 |
| Pst/Psh_17 | Avocet Yr9 | Wheat | 5 |
| Pst/Psh_17 | Bancroft | Barley | 0 |
| Pst/Psh_17 | Bigo | Barley | 0,5 |
| Pst/Psh_17 | Chinese 166 | Wheat | 0 |
| Pst/Psh_17 | Compair | Wheat | 0 |
| Pst/Psh_17 | Elymus repens | Other | 0 |
| Pst/Psh_17 | Emir | Barley | 7 |
| Pst/Psh_17 | Heils Franken | Barley | 0 |
| Pst/Psh_17 | Hiproly | Barley | 0 |
| Pst/Psh_17 | Hordeum spontaneum | Barley | 7 |
| Pst/Psh_17 | Hybrid 46 | Wheat | 0 |
| Pst/Psh_17 | I5 | Barley | 0 |
| Pst/Psh_17 | Kalyansona | Wheat | 1 |
| Pst/Psh_17 | Lee | Wheat | 2 |
| Pst/Psh_17 | Moro | Wheat | 0 |
| Pst/Psh_17 | Morocco | Wheat | 5 |
| Pst/Psh_17 | Sleipner | Wheat | 0,5 |

| | | | |
|---|---|---|---|
| Pst/Psh_17 | Suwon/Omar | Wheat | 0 |
| Pst/Psh_17 | TP 981 | Wheat | 0 |
| Pst/Psh_17 | Triticum dicoccon | Wheat | 0 |
| Pst/Psh_17 | Triticum monococcum | Wheat | 0 |
| Pst/Psh_17 | Trumpf | Barley | 0,5 |
| Pst/Psh_17 | VPM1 | Wheat | 0 |
| Pst/Psh_17 | Varanda | Barley | 0 |
| Pst/Psh_17 | Vilmorin 23 | Wheat | 0 |
| Pst/Psh_5 | Abed Binder | Barley | 0 |
| Pst/Psh_5 | Aegilops tauschii | Other | 0 |
| Pst/Psh_5 | Afzal | Barley | 7 |
| Pst/Psh_5 | Astrix | Barley | 0,5 |
| Pst/Psh_5 | Avocet S | Wheat | 3,5 |
| Pst/Psh_5 | Avocet SP | Wheat | 0 |
| Pst/Psh_5 | Avocet Yr1 | Wheat | 0,5 |
| Pst/Psh_5 | Avocet Yr15 | Wheat | 0 |
| Pst/Psh_5 | Avocet Yr17 | Wheat | 0 |
| Pst/Psh_5 | Avocet Yr24 | Wheat | 0 |
| Pst/Psh_5 | Avocet Yr27 | Wheat | 0 |
| Pst/Psh_5 | Avocet Yr32 | Wheat | 0,5 |
| Pst/Psh_5 | Avocet Yr5 | Wheat | 0 |
| Pst/Psh_5 | Avocet Yr6 | Wheat | 3 |
| Pst/Psh_5 | Avocet Yr7 | Wheat | 0,5 |
| Pst/Psh_5 | Avocet Yr8 | Wheat | 0 |
| Pst/Psh_5 | Avocet Yr9 | Wheat | 3,5 |
| Pst/Psh_5 | Bancroft | Barley | 0 |
| Pst/Psh_5 | Bigo | Barley | 0 |
| Pst/Psh_5 | Chinese 166 | Wheat | 0 |
| Pst/Psh_5 | Compair | Wheat | 0 |
| Pst/Psh_5 | Elymus repens | Other | 0 |
| Pst/Psh_5 | Emir | Barley | 7 |
| Pst/Psh_5 | Heils Franken | Barley | 0,5 |
| Pst/Psh_5 | Hiproly | Barley | 0 |
| Pst/Psh_5 | Hordeum spontaneum | Barley | 5,5 |
| Pst/Psh_5 | Hybrid 46 | Wheat | 0 |
| Pst/Psh_5 | I5 | Barley | 0 |
| Pst/Psh_5 | Kalyansona | Wheat | 6 |
| Pst/Psh_5 | Lee | Wheat | 1,5 |
| Pst/Psh_5 | Moro | Wheat | 0 |
| Pst/Psh_5 | Morocco | Wheat | 7 |
| Pst/Psh_5 | Sleipner | Wheat | 0 |
| Pst/Psh_5 | Suwon/Omar | Wheat | 0 |
| Pst/Psh_5 | TP 981 | Wheat | 0,5 |
| Pst/Psh_5 | Triticum dicoccon | Wheat | 0 |

| | | | |
|---|---|---|---|
| Pst/Psh_5 | Triticum monococcum | Wheat | 0 |
| Pst/Psh_5 | Trumpf | Barley | 0 |
| Pst/Psh_5 | VPM1 | Wheat | 0,5 |
| Pst/Psh_5 | Varanda | Barley | 5 |
| Pst/Psh_5 | Vilmorin 23 | Wheat | 0 |
| Pst/Psh_7 | Abed Binder | Barley | 0 |
| Pst/Psh_7 | Aegilops tauschii | Other | 0 |
| Pst/Psh_7 | Afzal | Barley | 7 |
| Pst/Psh_7 | Astrix | Barley | 0 |
| Pst/Psh_7 | Avocet S | Wheat | 2 |
| Pst/Psh_7 | Avocet SP | Wheat | 0 |
| Pst/Psh_7 | Avocet Yr1 | Wheat | 0,5 |
| Pst/Psh_7 | Avocet Yr15 | Wheat | 0 |
| Pst/Psh_7 | Avocet Yr17 | Wheat | 0 |
| Pst/Psh_7 | Avocet Yr24 | Wheat | 0 |
| Pst/Psh_7 | Avocet Yr27 | Wheat | 0 |
| Pst/Psh_7 | Avocet Yr32 | Wheat | 0,5 |
| Pst/Psh_7 | Avocet Yr5 | Wheat | 0 |
| Pst/Psh_7 | Avocet Yr6 | Wheat | 0,5 |
| Pst/Psh_7 | Avocet Yr7 | Wheat | 0,5 |
| Pst/Psh_7 | Avocet Yr8 | Wheat | 1,5 |
| Pst/Psh_7 | Avocet Yr9 | Wheat | 3 |
| Pst/Psh_7 | Bancroft | Barley | 0 |
| Pst/Psh_7 | Bigo | Barley | 0 |
| Pst/Psh_7 | Chinese 166 | Wheat | 7 |
| Pst/Psh_7 | Compair | Wheat | 0 |
| Pst/Psh_7 | Elymus repens | Other | 0 |
| Pst/Psh_7 | Emir | Barley | 7 |
| Pst/Psh_7 | Heils Franken | Barley | 0 |
| Pst/Psh_7 | Hiproly | Barley | 0 |
| Pst/Psh_7 | Hordeum spontaneum | Barley | 7 |
| Pst/Psh_7 | Hybrid 46 | Wheat | 0 |
| Pst/Psh_7 | I5 | Barley | 0 |
| Pst/Psh_7 | Kalyansona | Wheat | 6 |
| Pst/Psh_7 | Lee | Wheat | 2 |
| Pst/Psh_7 | Moro | Wheat | 0 |
| Pst/Psh_7 | Morocco | Wheat | 7 |
| Pst/Psh_7 | Sleipner | Wheat | 0,5 |
| Pst/Psh_7 | Suwon/Omar | Wheat | 0 |
| Pst/Psh_7 | TP 981 | Wheat | 0 |
| Pst/Psh_7 | Triticum dicoccon | Wheat | 0 |
| Pst/Psh_7 | Triticum monococcum | Wheat | 0 |
| Pst/Psh_7 | Trumpf | Barley | 0 |
| Pst/Psh_7 | VPM1 | Wheat | 0 |

| | | | |
|---|---|---|---|
| Pst/Psh_7 | Varanda | Barley | 3 |
| Pst/Psh_7 | Vilmorin 23 | Wheat | 0 |
| Pst/Psh_8 | Abed Binder | Barley | 0,5 |
| Pst/Psh_8 | Aegilops tauschii | Other | 0 |
| Pst/Psh_8 | Afzal | Barley | 7 |
| Pst/Psh_8 | Astrix | Barley | 0 |
| Pst/Psh_8 | Avocet S | Wheat | 3,5 |
| Pst/Psh_8 | Avocet SP | Wheat | 0 |
| Pst/Psh_8 | Avocet Yr1 | Wheat | 3 |
| Pst/Psh_8 | Avocet Yr15 | Wheat | 0 |
| Pst/Psh_8 | Avocet Yr17 | Wheat | 0 |
| Pst/Psh_8 | Avocet Yr24 | Wheat | 0 |
| Pst/Psh_8 | Avocet Yr27 | Wheat | 0 |
| Pst/Psh_8 | Avocet Yr32 | Wheat | 2 |
| Pst/Psh_8 | Avocet Yr5 | Wheat | 0 |
| Pst/Psh_8 | Avocet Yr6 | Wheat | 3 |
| Pst/Psh_8 | Avocet Yr7 | Wheat | 1 |
| Pst/Psh_8 | Avocet Yr8 | Wheat | 2 |
| Pst/Psh_8 | Avocet Yr9 | Wheat | 5 |
| Pst/Psh_8 | Bancroft | Barley | 0 |
| Pst/Psh_8 | Bigo | Barley | 0 |
| Pst/Psh_8 | Chinese 166 | Wheat | 7 |
| Pst/Psh_8 | Compair | Wheat | 0,5 |
| Pst/Psh_8 | Elymus repens | Other | 0 |
| Pst/Psh_8 | Emir | Barley | 7 |
| Pst/Psh_8 | Heils Franken | Barley | 0 |
| Pst/Psh_8 | Hiproly | Barley | 0 |
| Pst/Psh_8 | Hordeum spontaneum | Barley | 5 |
| Pst/Psh_8 | Hybrid 46 | Wheat | 0 |
| Pst/Psh_8 | I5 | Barley | 0 |
| Pst/Psh_8 | Kalyansona | Wheat | 2 |
| Pst/Psh_8 | Lee | Wheat | 2 |
| Pst/Psh_8 | Moro | Wheat | 0 |
| Pst/Psh_8 | Morocco | Wheat | 6 |
| Pst/Psh_8 | Sleipner | Wheat | 1 |
| Pst/Psh_8 | Suwon/Omar | Wheat | 0 |
| Pst/Psh_8 | TP 981 | Wheat | 0 |
| Pst/Psh_8 | Triticum dicoccon | Wheat | 0 |
| Pst/Psh_8 | Triticum monococcum | Wheat | 0 |
| Pst/Psh_8 | Trumpf | Barley | 0 |
| Pst/Psh_8 | VPM1 | Wheat | 0 |
| Pst/Psh_8 | Varanda | Barley | 5 |
| Pst/Psh_8 | Vilmorin 23 | Wheat | 0 |
| Pst/Psh_9 | Abed Binder | Barley | 0 |

| | | | |
|---|---|---|---|
| Pst/Psh_9 | Aegilops tauschii | Other | 0 |
| Pst/Psh_9 | Afzal | Barley | 7 |
| Pst/Psh_9 | Astrix | Barley | 0 |
| Pst/Psh_9 | Avocet S | Wheat | 2 |
| Pst/Psh_9 | Avocet SP | Wheat | 0 |
| Pst/Psh_9 | Avocet Yr1 | Wheat | 0,5 |
| Pst/Psh_9 | Avocet Yr15 | Wheat | 0 |
| Pst/Psh_9 | Avocet Yr17 | Wheat | 0 |
| Pst/Psh_9 | Avocet Yr24 | Wheat | 0 |
| Pst/Psh_9 | Avocet Yr27 | Wheat | 0 |
| Pst/Psh_9 | Avocet Yr32 | Wheat | 0,5 |
| Pst/Psh_9 | Avocet Yr5 | Wheat | 0 |
| Pst/Psh_9 | Avocet Yr6 | Wheat | 0 |
| Pst/Psh_9 | Avocet Yr7 | Wheat | 0,5 |
| Pst/Psh_9 | Avocet Yr8 | Wheat | 0 |
| Pst/Psh_9 | Avocet Yr9 | Wheat | 1 |
| Pst/Psh_9 | Bancroft | Barley | 0 |
| Pst/Psh_9 | Bigo | Barley | 0 |
| Pst/Psh_9 | Chinese 166 | Wheat | 7 |
| Pst/Psh_9 | Compair | Wheat | 0 |
| Pst/Psh_9 | Elymus repens | Other | 0 |
| Pst/Psh_9 | Emir | Barley | 7 |
| Pst/Psh_9 | Heils Franken | Barley | 0 |
| Pst/Psh_9 | Hiproly | Barley | 0 |
| Pst/Psh_9 | Hordeum spontaneum | Barley | 5 |
| Pst/Psh_9 | Hybrid 46 | Wheat | 0 |
| Pst/Psh_9 | I5 | Barley | 0 |
| Pst/Psh_9 | Kalyansona | Wheat | 1,5 |
| Pst/Psh_9 | Lee | Wheat | 2 |
| Pst/Psh_9 | Moro | Wheat | 0 |
| Pst/Psh_9 | Morocco | Wheat | 4 |
| Pst/Psh_9 | Sleipner | Wheat | 0,5 |
| Pst/Psh_9 | Suwon/Omar | Wheat | 0 |
| Pst/Psh_9 | TP 981 | Wheat | 0 |
| Pst/Psh_9 | Triticum dicoccon | Wheat | 0 |
| Pst/Psh_9 | Triticum monococcum | Wheat | 0 |
| Pst/Psh_9 | Trumpf | Barley | 0 |
| Pst/Psh_9 | VPM1 | Wheat | 0 |
| Pst/Psh_9 | Varanda | Barley | 0 |
| Pst/Psh_9 | Vilmorin 23 | Wheat | 0 |
